# A unique symbiosome in an anaerobic single-celled eukaryote

**DOI:** 10.1101/2023.03.03.530753

**Authors:** Jon Jerlström-Hultqvist, Lucie Gallot-Lavallée, Dayana E. Salas-Leiva, Bruce A. Curtis, Kristína Záhonová, Ivan Čepička, Courtney W. Stairs, Shweta Pipaliya, Joel B. Dacks, John M. Archibald, Andrew J. Roger

**Affiliations:** Department of Biochemistry and Molecular Biology, Dalhousie University, Halifax, Nova Scotia, Canada; Department of Biochemistry, University of Cambridge, Cambridge, United Kingdom; Department of Zoology, Faculty of Science, Charles University, Prague, Czech Republic; Division of Infectious Disease, Department of Medicine, Faculty of Medicine and Dentistry, University of Alberta, Edmonton, Alberta, Canada; Department of Parasitology, Faculty of Science, Charles University, BIOCEV, Vestec, Czech Republic; Institute of Parasitology, Biology Centre, Czech Academy of Sciences, České Budějovice (Budweis), Czech Republic; Life Science Research Centre, Department of Biology and Ecology, Faculty of Science, University of Ostrava, Ostrava, Czech Republic; Microbiology Group, Department of Biology, Lund University, Lund, Sweden

**Keywords:** ectosymbiosis, syntrophy, hydrogenosome, anaerobiosis, protists, evolution

## Abstract

Symbiotic relationships drive evolutionary change and are important sources of novelty. Here we demonstrate a highly structured syntrophic symbiosis between species of the anaerobic protist *Anaeramoeba* (Anaeramoebae, Metamonada) and bacterial ectosymbionts. We dissected this symbiosis with long-read metagenomics, transcriptomics of host and symbiont cells coupled with fluorescent in situ hybridization (FISH), and microscopy. Genome sequencing, phylogenomic analyses and FISH show that the symbionts belong to the *Desulfobacteraceae* and were acquired independently in two different *Anaeramoeba* species. We show that ectosymbionts likely reside deep within cell surface invaginations in a symbiosomal membrane network that is tightly associated with cytoplasmic hydrogenosomes. Metabolic reconstructions based on the genomes and transcriptomes of the symbionts suggest a highly evolved syntrophic interaction. Host hydrogenosomes likely produce hydrogen, acetate, and propionate that are consumed by the symbionts dissimilatory sulfate reduction, Wood-Ljungdahl and methylmalonyl pathways, respectively. Because the host genome sequences encode several vitamin B12-dependent enzymes but appear to lack the ability to biosynthesize this vitamin, we hypothesize that the symbionts supply their hosts with B12. We detected numerous lateral gene transfers from diverse bacteria to *Anaeramoeba*, including genes involved in oxygen defense and anaerobic metabolism. Gene families encoding membrane-trafficking components that regulate the phagosomal maturation machinery are notably expanded in *Anaeramoeba* spp. and may be involved in organizing and/or stabilizing the symbiosomal membrane system. Overall, the Anaeramoebae have evolved a dynamic symbiosome comprised of a vacuolar system that facilitates positioning and maintenance of sulfate-reducing bacterial ectosymbionts.

## Introduction

Syntrophic interactions between eukaryotes and prokaryotes are important drivers of evolutionary change. In oxygen-poor environments, eukaryotic cells have repeatedly evolved specialized symbiotic organs to increase syntrophic efficiency. These interactions are often centered on the exchange of metabolites produced by the host’s mitochondrion-related organelles (MROs), that perform metabolism adapted to anaerobic conditions. H_2_ produced by these MROs or ‘hydrogenosomes’, is an important currency in these interactions and is the end-product of the anaerobic ATP-producing metabolic pathway. Removal of H_2_ by prokaryotes likely increases the flux in these host pathways by avoiding product build-up and inhibition of anaerobic metabolism. As a result, H_2_-consuming symbionts are sometimes found intimately associated with host MROs, an arrangement that increases their access to substrate (Rotterová et al. 2022). Although the symbiotic organs of some animals have been comparatively well-studied, examples of structured symbioses in unicellular organisms are poorly understood.

The Anaeramoebae is a newly described phylum of anaerobic protists from the supergroup Metamonada (Stairs et al. 2021; Táborský, Pánek, and Čepička 2017). Anaeramoebid cells are predominantly amoeboid with a large mass of symbionts positioned close to the nucleus. The symbiont mass is tightly intercalated with double-membraned electron-dense organelles, identified as hydrogenosomes (Stairs et al. 2021; Táborský, Pánek, and Čepička 2017). The symbionts are not in direct contact with the hydrogenosomes; they are bound by a host-derived membrane (Táborský, Pánek, and Čepička 2017). Metabolic reconstruction of the *Anaeramoeba* hydrogenosomes showed that they likely produce H_2_, acetate, and propionate as end-products (Stairs et al. 2021). Here, we combine long-read whole genome sequencing, transcriptome sequencing of host and symbionts, phylogenomic analyses, and a variety of microscopy and *in situ* localization approaches to (i) dissect the *Anaeramoeba* holobiont, (ii) characterize the bacterial symbionts, and (iii) probe the host-symbiont metabolic and cellular interactions. We find that membrane-trafficking components shown in other organisms to regulate the phagosomal maturation machinery have expanded in *Anaeramoeba*. The interconnected vacuolar system within which the symbionts reside is a symbiotic organelle that allows *Anaeramoeba* species to capture, position and maintain symbionts.

## Results

### The Anaeramoeba symbionts are directly connected to the extracellular milieu

Actively feeding *Anaeramoeba* cells have a fan-shaped hyaline zone and trailing projections (Figure 1A). Near the nucleus in the bulbous cell body lies a large mass of symbionts apparently housed in vacuoles tightly positioned close to, but not in direct contact with, hydrogenosomes (Figure 1B-E) (Táborský, Pánek, and Čepička 2017). The symbionts are stably maintained throughout the cell cycle and segregated in an organized fashion during cell division (Figure S1). These observations indicate that the clusters are stable structures but do not reveal if the *Anaeramoeba* symbionts are endosymbionts completely enclosed within the host or are directly connected to the extracellular media. To distinguish between these two possibilities, we conducted live-cell pulse-labeling experiments with fluorescent-labeled Wheat Germ Agglutinin (WGA), a lectin that stains sialic acid and N-acetyl glucosamine in bacterial cell walls. The *A. flamelloides* BUSSELTON2 symbiont showed clear WGA labeling, with staining intensity comparable to free-living bacteria (Figure S2). This suggests the symbionts are in direct contact with the outside environment. Furthermore, TEM images of *A. ignava* BMAN show symbiont cells in pockets at the host cell membrane with connections to the surface (Figure 1F). We thus conclude that the symbionts of *Anaeramoeba* spp. reside in a network of deep, anastamosing invaginations of the cell’s membrane with extracellular openings.

**Figure 1:**
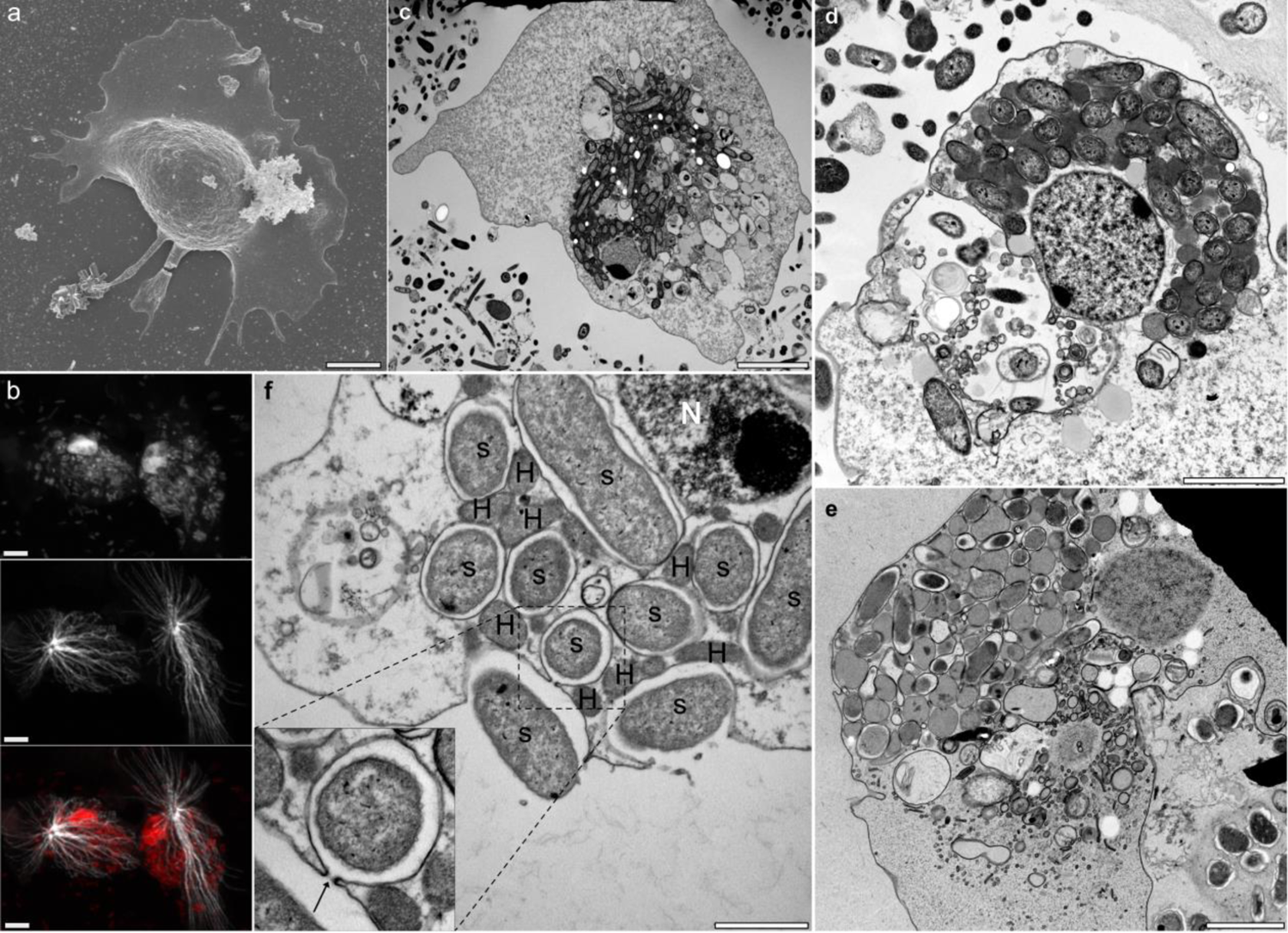
*Anaeramoeba* and symbionts. **a,** Scanning electron micrograph (SEM) of an *Anaeramoeba flamelloides* BUSSELTON2 amoebae showing a hyaline front and posterior trailing projections. Scale bar 5 µm. **b,** Immunolocalization of alpha-tubulin (TAT1 antibody, 1:200) in *A. flamelloides* BUSSELTON2 cells using laser scanning confocal microscopy. Host nuclei and symbiont DNA were stained using DAPI (top panel) and the acentriolar centrosome and radiating microtubules by alpha-tubulin TAT1 antibody (middle panel) with overlay (bottom panel). Scale bar 5 µm. **c,** TEM of chemically fixed *A. flamelloides* BUSSELTON2. Scale bar 5 µm. **d,** TEM of chemically fixed *A. flamelloides* BUSSELTON2. Scale bar 2 µm. **e,** TEM of cryo-fixed *A. flamelloides* BUSSELTON2. Scale bar 2 µm. **f,** Transmission electron micrograph (TEM) of *A. ignava* BMAN showing vesicle-bound symbionts (S) and host hydrogenosomes (H) in close proximity. The boxed inlay shows narrow openings connecting outside media to the vesicles housing the symbionts. Nucleus (N). Scale bar 1 µm.

### Anaeramoeba symbionts belong to Desulfobacteraceae and were acquired independently in different host species

We sequenced the nuclear genomes of three *Anaerameoba* isolates (*A. ignava* BMAN, *A. flamelloides* BUSSELTON2, and SCHOONER1) and the genomes of the associated prokaryotes using Nanopore long-read and Illumina short-read technologies (Table S1). The most abundant prokaryotes in the *A. flamelloides* and *A. ignava* genomic datasets were *Desulfobacteraceae*, hereafter referred to as Sym_BUSS2, Sym_SCH1, and Sym_BMAN. In *A. flamelloides*, where amoeba separation from suspended bacteria was found to be the most efficient (Figure S3), the symbiont lineages were detected almost exclusively in the amoeba enrichment fraction and were virtually non-detectable in the culture supernatant (Figure S3). Fluorescence *in situ* hybridization using unique probes designed against the 16S rDNA genes in the genomes of Sym_BMAN (Figure 2A-D) and Sym_BUSS2/Sym_SCH1 (Figure 2F-L) (as well as less specific probes for broader encompassing taxonomic groups (Figure S4)) confirmed the identity of the symbionts. The average number of symbionts per host based on symbiont genome coverage relative to nuclear genome coverage were estimated to be 35.3-36.5 for *A. flamelloides* (Sym_BUSS2 & Sym_SCH1) and 12.6 for *A. ignava* (Sym_BMAN) (the ploidy of the host nuclear genome is unknown but posited to be haploid). All three symbiont genomes are large (4.97-6.06 Mbp) and relatively gene-rich (3,823-5,286 intact genes) (Table S2). The Sym_BUSS2 and Sym_SCH1 genomes are 99.7% identical at the nucleotide level but show large differences in synteny (Figure S5), whereas the Sym_BMAN genome is highly divergent relative to the two *A. flamelloides* symbionts (average nucleotide identity of 74.1-74.2%). A second less abundant *Desulfobacteraceae* genome, Sym_BMAN2, was also detected in the *A. ignava* dataset but its status as a symbiont could not be established using FISH. Thus, for *A. ignava* we focused on Sym_BMAN in subsequent analyses.

**Figure 2:**
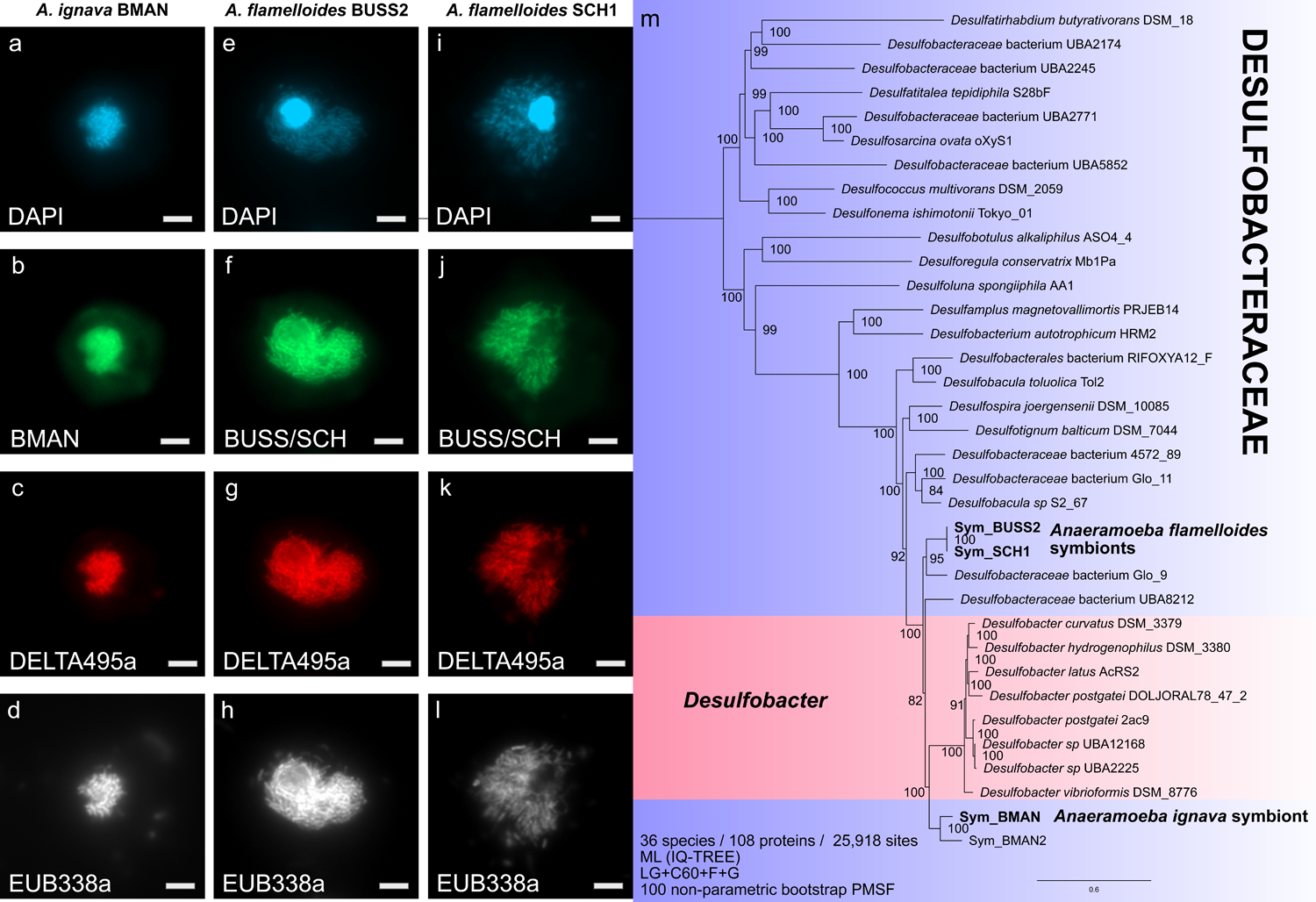
*Anaeramoeba* symbionts are closely related to *Desulfobacter* and were acquired in separate events. **a-d,** *A. ignava* BMAN stained with **a,** DAPI and hybridized with **b,** probe DSBA355-BMN-488, **c,** probe Delta495a-Atto 550 and **d,** probe EUB338a-Atto 633. **e-h,** *A. flamelloides* BUSSELTON2 stained with **e,** DAPI and hybridized with **f,** probe BUSS/SCH-BMN-488, **g,** probe Delta495a-Atto 550 and **h,** probe EUB338a-Atto 633. **i-l,** *A. flamelloides* SCHOONER1 stained with **i,** DAPI and hybridized with **j,** probe BUSS/SCH-BMN-488, **k,** probe Delta495a-Atto 550 and **l,** probe EUB338a-Atto 633. Scale bar 1 µm. **m,** The phylogenomic analysis was based on 36 taxa, 108 proteins, and 25,918 sites with IQTree v2.2.0.3 (LG+C60+F+G model of evolution). Bipartition support values are derived from 100 non-parametric bootstraps under the PMSF model. Scale bar indicates inferred number of substitutions per site. Tree files and alignments are available at FigShare: https://doi.org/10.6084/m9.figshare.20375619

Phylogenetic analysis of 15 ribosomal proteins from 165 *Desulfobacterales* genomes placed the symbionts as members of the family *Desulfobacteraceae* (Figure S6), closely related to the genus *Desulfobacter*. To improve resolution, we performed phylogenetic analysis on a data set of 108 proteins from a phylogenetically restricted set of Desulfobacteraceae. Sym_BMAN and Sym_BMAN2 belong to a clade that branches sister to a large *Desulfobacter* group (Figure 2M) whereas Sym_BUSS2/SCH1 branches separately, forming a well-supported group (BS 95%) with an environmental isolate (Glo_9) recovered from a metagenome of the benthic foraminiferan *Globobulimina* spp. The latter clade forms a sister group to the *Desulfobacter* – Sym_BMAN clade. The *Desulfobacter*-like symbionts of *A. ignava* and *A. flamelloides* thus appear to have been acquired independently by their respective hosts.

### The Anaeramoeba-Desulfobacter symbiosis was recently established

The Sym_BUSS2 and Sym_SCH1 genomes were found to have 922 and 1,000 pseudogenes, respectively (Table S2, S4), and >600 Insertion Sequences (IS) per genome (Table S5). In contrast, pseudogene and IS element quantities in Sym_BMAN are more than an order of magnitude lower and fall within the range of free-living *Desulfobacteraceae* species (Table S2). The IS elements of Sym_BUSS2 and Sym_SCH1, many of which have an intact transposase and are likely active, showed no strong evidence of clustering (Figure S7A,B). However, >200 IS elements were found to be close (<1,500 bp) to the edges of syntenic blocks, suggesting that the apparent high frequency of rearrangements is connected to IS element activity (Figure S7C,D). The Sym_Buss2 and Sym_SCH1 genomes are enriched in pseudogenes with predicted gene ontologies related to in signal transduction (T) and amino acid metabolism & transport (E) functional categories, whereas translation (J) and transcription (K) classes showed the opposite trend (Figure S8). The genomes of these two symbionts have flagellar operons (Figure S9), type IV pili, and CRISPR systems that are all extensively pseudogenized; both symbionts also appear to be impaired in their ability to decorate lipid A with O-antigen. In contrast, the Sym_BMAN genome encodes an intact type I-F CRISPR system with an array of 25 spacers as well as two convergent, 25 gene operons for a type VI secretion system (T6SS) (Figure S10). The greater degree of genome degeneration seen in the Sym_BUSS2 and Sym_SCH1 genomes relative to the largely intact Sym_BMAN genome strongly suggests that the former are ‘older’ symbionts and that they are unable to survive without the host.

However, the relatively large genomes of all *Anaeramoeba* symbionts and other genomic features are reminiscent of early-stage, recently-acquired symbionts (Moran and Plague 2004). Constitutive overexpression of the chaperonin GroEL/S is regarded as a critical factor in stabilizing endosymbionts in a wide range of insect symbioses (Kupper et al. 2014). Notably, metatranscriptomic analysis of the *A. ignava* BMAN and *A. flamelloides* BUSSELTON2 cultures show that the *groEL, groES* and HSP70-encoding *dnaK* genes are highly expressed in Sym_BUSS2 but moderately to lowly expressed in Sym_BMAN, in line with their respective degrees of genome erosion (Table S6).

### Anaeramoeba symbionts are metabolically poised to use hydrogenosome metabolites

The hydrogenosomes of both *Anaeramoeba* species are predicted to produce H_2_, acetate, and propionate as end-products (Stairs et al. 2021). To investigate whether the symbionts express genes predicted to be important for H_2_-uptake, we performed metatranscriptomics of the *A. ignava* BMAN and *A. flamelloides* BUSSELTON2 cultures. Metatranscriptomics of Sym_BUSS2 showed that the most prominently symbiont-expressed genes include those involved in dissimilatory sulfate reduction (DSR), the group 1b uptake [Ni/Fe] hydrogenase (*hynAB*), the Wood-Ljungdahl (W-L) pathway, and ATP synthase (Figure 3) (Table S6). Similarly, in Sym_BMAN, the *dsr* genes, *aprAB* pathway and *hynA*B were highly expressed, while W-L pathway and ATP synthase genes are less expressed than in Sym_BUSS2 (Table S6). Sym_BMAN shows high expression of an acetate permease (*actP*) that might act to bring in host-derived acetate to feed the W-L pathway. Although we could not identify acetate transporter genes (*actP* or *satP*) in the *A. flamelloides* symbiont genomes, we suspect these symbionts might be able to acquire acetic acid by diffusion across the membrane as reported in other systems (Grime et al. 2008). The symbionts appear to be able to activate propionate via propionyl-CoA synthase (*prpE*), which is highly expressed in Sym_BUSS2 and Sym_BMAN (Table S6). Propionyl-CoA likely feeds into the methylmalonyl-CoA (MMA) pathway to produce pyruvate that, via oxidative decarboyxlation by pyruvate:ferredoxin oxidoreductase, generates acetyl-CoA that could enter the W-L pathway (Figure 3). Most enzymes of the MMA pathway are highly and moderately expressed in Sym_BUSS2 and Sym_BMAN, respectively. Because Sym_BUSS2 is closely related to the *Desulfobacter* Glo_9 denitrifying symbiont of *Globobulimina* spp. (discussed in (Woehle et al. 2018; 2022)), we investigated whether the *Anaeramoeba:*Sym_BUSS2 symbiotic system could be based on denitrification. However, we failed to find denitrification-related genes in the host (*nrt*, *nirK*, *nor*) or symbionts (*napA*, *nozA*)

**Figure 3:**
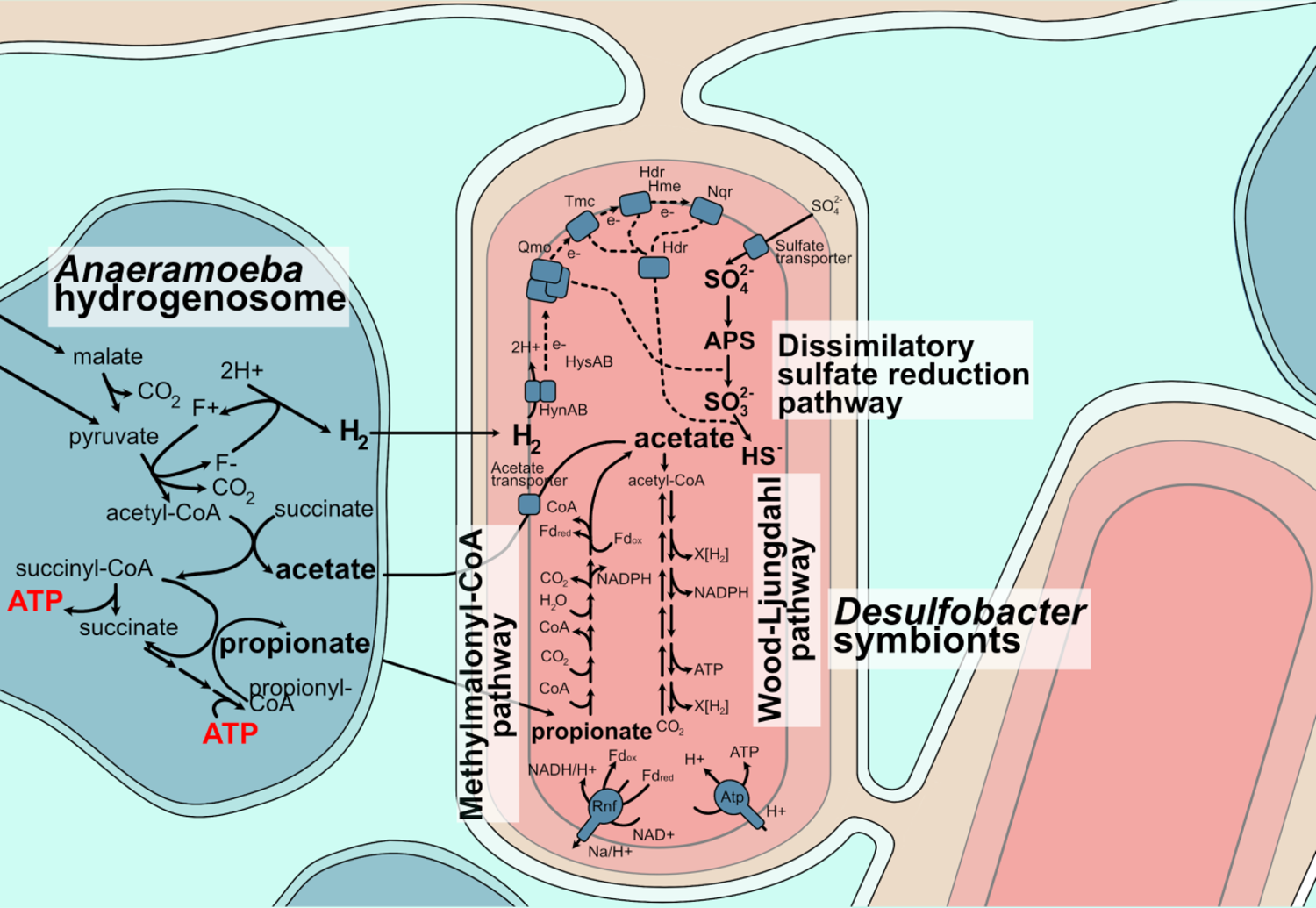
*Anaeramoeba* symbionts are metabolically poised for syntrophic interaction with hydrogenosomes. Suggested syntrophic interactions between *Anaeramoeba* hydrogenosomes (blue) and symbionts (pink/salmon) based on metabolic reconstruction from transcriptomic and genomic evidence. The ATP-producing hydrogenosomes generate H2, acetate, and propionate as end-products (in bold). Based on metatranscriptomic data, the symbionts use the hydrogenosome products by prominently expressing the dissimilatory sulfate reduction (DSR), methylmalonyl-CoA, and the Wood-Ljungdahl pathways. The symbionts are in deep membrane-pits with a connection to the cell surrounding that gives ready access to sulfate (gold). Abbreviations: HynAB - periplasmic [NiFe] hydrogenase, HysAB - [NiFeSe] hydrogenase, Qmo - QmoABC complex, Tmc - TmcABCD complex, Hdr - Heterodisulfide reductase, Hme - DsrMKJOP complex, Nqr - NADH:ubiquinone oxidoreductase, Rnf - Rnf complex, Atp - ATP synthase, APS - adenosine 5’-phosphosulfate, CoA - Coenzyme A, NAD - Nicotinamide adenine dinucleotide, NADPH - Nicotinamide adenine dinucleotide phosphate, Fdred/ox - Ferredoxin reduced/oxidized, ATP - adenosine triphosphate

Given that many of the gene products described above are sensitive to oxygen, we examined the genomes of the symbionts for oxygen or reactive oxygen species (ROS) defense systems. Genes encoding the superoxide detoxification (superoxide reductase and rubredoxin) and hydrogen peroxide detoxficiation (rubrerythrin) were highly expressed by Sym_BUSS2 and Sym_BMAN (Table S6).

### Anaeramoeba nuclear genomes are A+T rich and have extensively expanded gene families

The nuclear genomes of Anaeramoebae are A+T-rich, with the *A. ignava* BMAN nuclear genome (80.65% A+T) and intron (91.86% A+T) sequences being among the highest ever reported in eukaryotes (see for example; *P. falciparum* A+T genome (80.6%), intron (86.5%)(Gardner et al. 2002)) (Table S1,S3A). Whereas the assembled genomes of *A. flamelloides* are almost six times larger than for *A. ignava*, the number of protein coding genes is only two times higher (Table S1). The sequence divergence between the *Anaeramoeba* species is substantial with only 37.3% average amino acid identity between orthologous genes (Figure S11). In contrast, the *A. flamelloides* BUSSELTON2 and SCHOONER1 strains are much more similar (94.8% identity between orthologs).

To investigate the evolution of gene content in Anaeramoebae, we compared the gene families of the three *Anaeramoeba* genomes to nine other microbial eukaryotes including those from Parabasalia (*T. vaginalis*), Oxymonadida (*M. exilis*), Fornicata (*C. frisia*, *C. membranifera*, *K. bialata*, *G. intestinalis*, *S. salmonicida*,), Heterolobosea (*N. gruberi*), and Amorphea (*D. discoideum*). Across all genomes, a total of 565 and 4,704 families were deemed core and accessory, respectively (Table S7A-C). 1,806 gene families are taxon-specific, and of these, 67, 74 and 107 are specific to BUSSELTON2, SCHOONER1 and BMAN, respectively (Table S7D). When comparing *Anaeramoeba* to other metamonads, *N. gruberi* and *D. discoideum*, 92 families were considered specific to *Anaeramoeba* (Table S7E). Note that only 557 core and 1,557 accessory families were retained for the protein family expansion/contraction analysis across all taxa after filtering. Expansions ≥ 5-fold were observed in 110 core and 448 accessory families, whereas contractions were only detected in 18 accessory families (Table S7F,G). When comparing only *A. flamelloides* to *A. ignava*, we identified 2,597 core and 688 accessory families but only 2,578 and 89 of those were retained for the analysis (Table S7F). Of these, expansions were detected in 330 core families while contractions were found in 13 core families (Table S7H). The contractions seen in *A. flamelloides* likely correspond to expansions in *A. ignava* rather than contractions in *A. flamelloides* (Table S7F,H). Together with the presence of BUSSELTON2, SCHOONER1 and BMAN-specific families already mentioned, these differences highlight the striking differences among these relatively closely related taxa.

Many gene family expansions occur specifically in *A. flamelloides* and are enriched in genes involved in RNA and DNA metabolism and membrane-trafficking systems. One striking feature is the massive expansion of ribosomal proteins with more than 1,100 genes each in *A. flamelloides* compared to the 82 genes in *A. ignava*, a more typical number for a eukaryote (Suppl. Text, Table S7I-K; Figure S12). However, expansions of gene families connected to nutrient exchange were also noted. For example, *A. flamelloides* and *A. ignava* have sizable expansions of an Amt/MEP ammonium transporter gene, whose products likely function in the uptake or secretion of ammonia, a primary nitrogen source for cells (X.-D. Li et al. 2006).

### Lateral gene transfer is ongoing in Anaeramoeba

Lateral gene transfer (LGT) is increasingly recognized as a factor in eukaryotic evolution (Sibbald et al. 2020; Cote-L’Heureux, Maurer-Alcalá, and Katz 2022). We performed a phylogenomic analysis to infer LGTs from prokaryotes and viruses to *Anaeramoeba*. We identified 612, 1,414 and 1,359 putative foreign genes in the BMAN, BUSSELTON2, and SCHOONER1 genomes, respectively (4.1%, 4.7%, 4.5% of the total number of genes per genome). Accounting for gene duplications after acquisition, this corresponds to 388, 781 and 777 gene families (Table 1). These are some of the highest LGT proportions ever reported for protists (Van Etten and Bhattacharya 2020). Interestingly, most of the LGTs were inferred to have happened after the divergence between *A. ignava* and *A. flamelloides* (Table S8, Figure 4). 169 putative LGTs occurred in a common ancestor of the two species, 577 LGTs map to the base of the divergence between the *A. flamelloides* BUSSELTON2 and SCHOONER1 strains, and 220 were acquired in a more recent ancestor of *A. ignava* (this pattern may also be explained in part by differential loss of LGT genes). Approximately 30 genes were inferred to have been acquired in the two *A. flamelloides* strains after their divergence.

**Figure 4:**
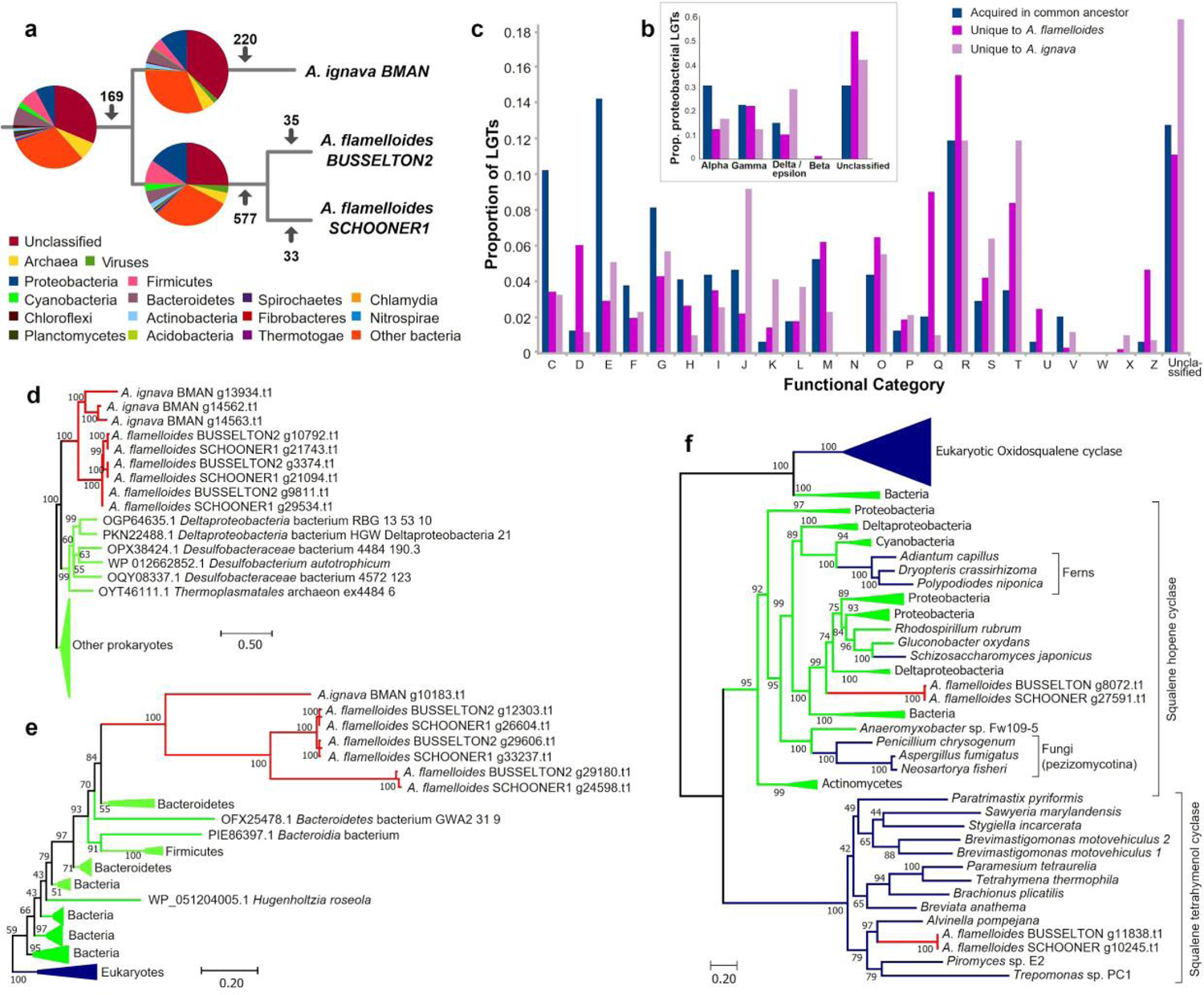
Taxonomic range and functional categories of LGT donors in *Anaeramoeba*. **a**, Taxonomy of LGT donors for genes inferred to have been acquired in the common ancestor of *Anaeramoeba* and separately in *A. flamelloides* and *A. ignava*. The numbers on the branches are the inferred number of LGTs on each position of the tree. **b,** Breakdown of LGTs from proteobacterial donors. **c,** Functional categories of genes inferred to have been acquired in the *Anaeramoeba* common ancestor, in *A. flamelloides* and *A. ignava*. Color coding is the same as for part b. **d,** Maximum-likelihood phylogeny of FAD-dependent oxidoreductases. **e,** Maximum-likelihood phylogeny of vitamin B12-dependent methionine synthase MetH. The trees shown in **d** and **e** were produced by our LGT-detection pipeline (see text). **f,** Maximum-likelihood phylogeny of oxidosqualene cyclase (OSC), squalene-hopene cyclase (SHC), and squalene-tetrahymanol cyclase (STC). The tree shown stems from analyses based on previously published datasets (Takishita et al. 2017; Bouwknegt et al. 2021). Sequences were aligned with MAFFT, sites were selected using BMGE, and the phylogeny inferred using IQTree model C20+G4 with 1,000 ultrafast bootstraps. *Anaeramoeba* sequences are in red, eukaryotic sequences in blue, and prokaryotic sequences in bright green. Scale bars indicate the inferred number of amino acid substitutions per site. Abbreviations: INFORMATION STORAGE AND PROCESSING: [J] Translation, ribosomal structure and biogenesis; [K] Transcription; [L] Replication, recombination and repair; CELLULAR PROCESSES AND SIGNALING: [D] Cell cycle control, cell division, chromosome partitioning; [V] Defense mechanisms; [T] Signal transduction mechanisms; [M] Cell wall/membrane/envelope biogenesis; [N] Cell motility; [Z] Cytoskeleton; [W] Extracellular structures; [U] Intracellular trafficking, secretion, and vesicular transport; [O] Posttranslational modification, protein turnover, chaperones. METABOLISM: [C] Energy production and conversion; [G] Carbohydrate transport and metabolism; [E] Amino acid transport and metabolism; [F] Nucleotide transport and metabolism; [H] Coenzyme transport and metabolism; [I] Lipid transport and metabolism; [P] Inorganic ion transport and metabolism; [Q] Secondary metabolites biosynthesis, transport and catabolism. POORLY CHARACTERIZED: [R] General function prediction only; [X] Mobilome: prophages, transposons; [S] Function unknown.

**Table 1:** Predicted LGTs in the genomes of *Anaeramoeba* species.

|  | Predicted proteins | Ortho groups (OGs) | OGs including at least one prokaryotic or viral sequence | Trees | Inferred LGT events | Total LGT-derived genes |
| --- | --- | --- | --- | --- | --- | --- |
| *A. ignava* BMAN | 15,022 | 6,168 | 2,224 | 1,743 | 388 | 612 |
| *A. flamelloides* BUSSELTON2 | 29,927 | 14,259 | 3,701 | 3,156 | 781 | 1,414 |
| *A. flamelloides* SCHOONER1 | 30,041 | 14,246 | 3,679 | 3,115 | 777 | 1,359 |

Most of the LGTs for which a taxonomic origin could be inferred are bacterial, although archaeal and viral contributions to *Anaeramoeba* are also apparent (Figure 4A). Proteobacteria, Firmicutes and Bacteroidetes (Figure 4A) are the most represented donor phyla contributing genes to the ancestor of *A. ignava* and *A. flamelloides*, and the two species appear to have acquired more proteobacterial genes since they diverged from one another. Interestingly, Deltaproteobacteria is not the most represented class, despite the taxonomic affinity of *Anaeramoeba*’s symbionts (Figure 2). That said, several dozen genes with amino acid identities >60% (Table S8D) are inferred to have been transferred from Deltaproteobacteria to *Anaeramoeba*, such as an aspartate ammonia ligase (in *A. flamelloides*) and an FAD-dependent oxidoreductase (in both species) (Figure 4D). A few *Anaeramoeba* genes have discernable homologs only in the symbionts and thus might represent symbiont-to-host LGTs. Curiously, these LGTs encode long repetitive proteins (up to 4,100 amino acids). The predicted relative contributions of Alpha- and Gammaproteobacteria are similar for *A. ignava* and *A. flamelloides* (Figure 4B).

### Laterally transferred genes shape Anaeramoeba biology

The predicted functions of LGTs in *Anaeramoeba* differ between those acquired in the common ancestor of *A. ignava* and *A. flamelloides* and those acquired after the divergence of the two species (Figure 4C) (Suppl. Text). Several genes acquired in the *Anaeramoeba* common ancestor appear to be related to accommodating sulfate-reducing symbionts. For example, genes encoding a cysteine synthase K and D-3-phosphoglycerate dehydrogenase are LGT-derived (Figures S13D,E); together with SerC phosphoserine aminotransferase (which was not detected in our LGT screen but was found in the genomes), these proteins link glycolysis to the recycling of SH^-^ produced by the symbionts and the generation of acetate that might be used by the symbiont (Figure S14). A prokaryotic acetate transporter gene (Figure S13F) also appears to have been acquired in the *Anaeramoeba* ancestor and might have played a role in the establishment of the symbiosis (the symbionts are predicted to use acetate produced by *Anaeramoeba*). The foreign acetate transporter gene is present in >10 copies in each *Anaeramoeba* genome.

We scanned the *Anaeramoeba* LGTs for putative host-symbiont recognition factors, which often have repetitive domains (Hinzke et al. 2019). Amongst the LGT candidates, we identified 38 gene families that previously have been associated with host-symbiont interactions (Table S8). Some of these gene families are highly amplified, have predicted signal peptide-encoding regions, and show differential presence/abundance between *A. ignava* and *A. flamelloides*, indicating that they might traffic in the secretory pathway and could potentially mediate symbiont interactions. Several foreign genes in *A. ignava* BMAN and *A. flamelloides* BUSSELTON2 are clearly involved in anaerobic life, including (i) MRO metabolism (Stairs et al. 2021), (see Table S8), (ii) oxygen detoxification, (iii) ATP production in the absence of oxidative phosphorylation (e.g., pyruvate phosphate dikinase; Figure S13G), and (iv) cell membrane composition modification (acquisition of a bacterial squalene hopene cyclase (SHC) involved in oxygen-free biosynthesis of hopanoids (Suppl. Text, Figure 4F). Several of these genes appear to have been acquired on multiple occasions from various donors (Suppl. Text; e.g. Figure S13M,O).

Of the LGT-derived genes that are amenable to functional / metabolic prediction, very few make up a complete “module” (Aramaki et al. 2020), suggesting that most LGT-derived proteins function in concert with host-origin enzymes in mosaic pathways. The sole exception is a complete set of enzymes (four) constituting a putative Leloir pathway in *A. flamelloides*, which suggests that the organism can metabolize galactose (Suppl. Text). Interestingly, while a complete Leloir pathway is present in the BMAN symbiont, *galE* appears to have been lost in the *A. flamelloides* symbionts.

### Anaeramoeba might acquire vitamin B12 from its symbionts

Vitamin (vit)B12 is one of the most complex coenzymes known. Its biosynthesis involves ∼30 enzymatic steps and is confined to certain bacterial and archaeal species (Raux, Schubert, and Warren 2000). The symbionts associated with both *Anaeramoeba* species encode a complete pathway for vitB12 synthesis (except for alpha-ribazol phosphatase (*cobC*); below). Interestingly, our genome screens detected up to six vitB12-dependent enzymes, a surprising number and a record among eukaryotes examined thus far, encoded in the *Anaeramoeba* host genomes. This includes two enzymes in *A. flamelloides* (Suppl. Text) that have not been found in a eukaryote before. Additionally, both *Anaeramoeba* species have acquired a class II ribonucleotide reductase (RNR) and a methionine synthase (*metH*) (Figure 4E) from bacteria and have a eukaryotic methylmalonyl-CoA mutase and its associated GTPase MeaB. These observations suggest that vitB12—or cobalamin—is produced by the symbionts and consumed by the host (both *Anaeramoeba* species encode a cobalamin adenosyl transferase that catalyzes the conversion of cobalamin into adenosylcobalamin, the form used as a cofactor by vitB12-dependent methylmalonyl-CoA mutase). The exchange of vitB12 between prokaryotic producers and B12-auxotrophic algae was hypothesized elsewhere (Croft et al. 2005; Grant et al. 2014). Curiously, we found a laterally acquired gene encoding CobC (Figure S13W) in *A. ignava.* This enzyme catalyzes the last step of cobalamin synthesis, which is not predicted to occur in the symbionts. The *cobC* gene has also been acquired in some vitB12 auxotrophic diatoms (Vancaester et al. 2020). Our analyses strongly suggest that vitB12 exchange is part of the host-symbiont metabolic interaction in *Anaeramoeba*, a hypothesis that can be tested in the future.

### Expansions of endosome/phagosome modulating membrane-trafficking proteins in Anaeramoeba

The vesicle bound symbiont mass in these Anaeramoebae, which is connected to the extracellular environment, is reminiscent of phagosomes that have been delayed or frozen in their maturation process. The general expansions in genes involved in membrane-trafficking systems in both *Anaeramoeba* spp. identified above prompted us to specifically investigate several sets of proteins, which in other eukaryotes have been implicated in regulation of phagosomal maturation. Certain Rab GTPases are essential in this process, mediating early to late endosomal conversion, as well as endosomal function and are involved in other regulatory aspects of the maturation process (Homma, Hiragi, and Fukuda 2021). Consequently, we investigated not only Rabs but also their GTPase Activating Proteins (GAPs) and the Vps-C complexes HOPS and CORVET that mediate endosomal Rab conversion, with the former having recently been implicated in regulating phagosomal-lysosomal fusion (Jeschke and Haas 2018).

The *Anaeramoeba* genomes encode many Rab GTPases, almost 400 in *A. flamelloides*. Rabs are small proteins, which made resolved classification by phylogenetic analysis intractable. Orthofinder (Emms and Kelly 2019) was thus used for initial classification followed by targeted phylogenies of related Rab sub-families (when necessary). While 75-80% of Rab proteins could not be confidently classified, clear trends were nevertheless apparent (Table S10). We observed relatively low numbers of Rabs 6, 7, and 18, and no clear candidate orthologs for eight Rab sub-families were found, including some endosomal Rabs such as Rab 4 or 22. By contrast, Rab GTPase families 1, 2, 5, 7, 11, 14 and 20 were found to be highly expanded (Figure S15A-C). We were also able to confidently classify most Tre-2/Bub2/Cdc16 (TBC) proteins that serve as GTP-activating proteins for Rabs. Although there are only one or several members of each of the TBC-I, M, and N subfamilies, the TBC-E, F, K, and Q subfamilies are notably expanded (Figure S15D). Finally, we observed expansions in the complement of both HOPS and CORVET subunits, most obviously with increased numbers of the HOPS-specific subunits (Table S10).

We searched for evidence of gene duplications within the Rab and TBC subfamilies that might predate the divergence of the *Anaeramoeba* spp., and which could indicate *Anaeramoeba*-specific machinery involved in capture and regulated retention/digestion of symbionts. Rab1, or the metazoan-specific paralog Rab35 (Diekmann et al. 2011), has been directly implicated in phagosome maturation dynamics in mammalian cells (Haley and Zhou 2021) and in the amoebozoan parasite *Entamoeba* (Verma and Datta 2017). Although the phylogenetic resolution between the subfamilies is poor, in the case of the Rab1 sub-family (Figure S15C), and its regulator TBC-E (Figure S15D), tree topologies are consistent with gene duplication and diversification prior to the split of the *Anaeramoeba* spp. We also observed three strongly supported clades within TBC-K (Figure S15D) that predate the *Anaeramoeba* split. TBC-K regulates another GTPase Arf6 (Falace et al. 2010), which was recently shown in metazoan systems to regulate phagosomal maturation. Notably Rab35 and Arf6 operate in multiple mammalian systems in a cooperatively mutually antagonistic mechanism (Chesneau et al. 2012; Kobayashi and Fukuda 2012).

Collectively, these analyses show expansions of the complement of membrane-trafficking components in *Anaeramoeba* spp. that, in other organisms, regulate phagosomal maturation. But this is not true of all membrane-trafficking machinery. A recent study (Maciejowski et al. forthcoming) of the vesicle coat forming machinery (including coats that act within the endosomal system) found these proteins to be largely encoded in single copy in A. *ignava*, resembling the canonical eukaryotic complement. Overall, our findings are consistent with the existence of a specialized modulation and control system for the symbiont-containing vacuoles that allows *Anaeramoeba* spp. to regulate the acquisition and maintenance of its symbiont mass.

## Discussion

Symbiotic organs have evolved on numerous occasions across the tree of life and their origins are associated with massive changes to host cell biology and physiology (Fronk and Sachs 2022). Our analyses suggest that the ancestor of *Anaeramoeba* spp. evolved a subcellular symbiosome to house *Desulfobacter*-related symbionts in a way that has drastically reshaped their cell biology and metabolic capacities. The notable expansion of some membrane-trafficking genes in the *Anaeramoeba* common ancestor suggests that the membranous ‘organ’ housing the symbionts is a highly evolved structure that might allow them to selectively manage their captured symbionts and position them in tight association with their hydrogenomes. To our knowledge the only vaguely similar system occurs in certain anaerobic ciliates (*Scuticociliatia* and *Karyorelictea*) (Rotterová et al. 2022; Beinart et al. 2018), the heterolobosean *Psalteriomonas lanterna*, and obligately symbiotic parabasalids (*Trichonympha* and *Spirotrichonympha*), which are found in the termite digestive tract (Sato et al. 2009; Kuwahara et al. 2017). These organisms form elaborate relationships and metabolic dependencies with endosymbiotic and ectosymbiotic prokaryotes, with some of the latter being housed in host-derived cell membrane invaginations with hydrogenosomes in close proximity (Sato et al. 2009; Kuwahara et al. 2017). While the symbionts in *Anaeramoeba* are almost completely compartmentalized, they nevertheless retain connections to the external environment, presumably due to metabolic constraints such as the need to access sulfate, as seen in the *Desulfovibrio* symbionts of *Trichonympha* (Kuwahara et al. 2017; Takeuchi et al. 2020). Brown-pigmented karyorelictid ciliates in anoxic sediments house several symbiont consortia including a member of *Desulfobacteraceae* that are not closely related to the Anaeramoebae symbionts (Figure S6). These ciliate symbionts heavily transcribe genes needed for dissimilatory sulfate reduction even though they appear to be endosymbionts residing in double-membraned vacuoles with no obvious mechanism for sulfate provisioning (Beinart et al. 2018).

The specificity of symbiotic organs for colonizing prokaryotes can be multifactorial, influenced by host and symbiont metabolic complementarity, receptor recognition, and host secondary metabolite production, which inhibits disruptive colonizers or pathogens (Clay 2014). Intriguingly, the *Anaeramoeba* genomes sequenced herein have many LGT-derived genes previously implicated in protein-protein interactions and cell-cell adhesion in host-symbiont systems (Hinzke et al. 2019), as well as several non-ribosomal peptide synthases (Table S9) whose products might be influencing which bacteria can successfully colonize the symbiosome.

Hydrogen depletion is likely to be an important reason for host dependence on the *Anaeramoeba* symbionts. However, the large repertoire of host-encoded vitB12-dependent enzymes includes essential proteins involved in hydrogenosome metabolism and nucleotide scavenging, suggesting that a degree of metabolic complementarity exists and serves as a stabilizing factor. The lateral acquisition of CobC, the final enzyme in vitB12 biosynthesis, in *A. ignava* BMAN may help to optimize the synthesis of vitB12. Diatoms have been shown to strongly induce vitB12-binding proteins under limiting growth conditions (Bertrand et al. 2012) and modeling predicts active vitB12 release from bacteria in co-cultures (Grant et al. 2014). How vitB12 is harvested from bacteria by microbial eukaryotes is not clear but directed release and lysis have both been suggested (Grant et al. 2014; Nef et al. 2022).

Whereas evidence for the obligate nature of the *A. flamelloides* symbionts is strong (i.e. the high degree of genome degeneration), the degree of autonomy of the *A. ignava* symbiont is less clear. Having only examined three *Anaeramoeba* strains and their symbionts, we do not know whether all species in this genus exclusively establish symbioses with *Desulfobacteraceae* or whether they can harbor other compatible strains or consortia that deplete hydrogen and provide vitB12 biosynthesis. Hydrogen consumption may be carried out by other taxa such as *Arcobacter* spp. (Hamann et al. 2016), some of which are predicted to also be vitB12 prototrophs, or additional Deltaproteobacteria that are present in our *Anaeramoeba* cultures (Figure S3). If *Anaeramoeba* species are truly dependent on symbiont-provided vitB12, then this might also mean that cyclical symbiont-replacement, as seen in the ciliate *Euplotes* (Boscaro et al. 2022), is necessary to ensure that they receive a steady source of vitB12 in the face of symbiont genome erosion. Our LGT analyses indicate that no single bacterial lineage has disproportionally contributed to the gene repertoire of *Anaeramoeba*, which is reminiscent of the shopping bag model of gene acquisitions in the presence of symbiont-derived organelles (Howe et al. 2008). Although we detected several cases of transfers from *Desulfobacteraceae,* they are not obviously from the resident symbiont lineages. Some of the LGTs we observed might also be derived from past symbionts that have since been replaced by the current lineages.

The numbers of predicted foreign genes in *Anaeramoeba* is higher than for most microbial eukaryote genomes investigated (Van Etten and Bhattacharya 2020). Many of these LGTs appear to be related to adaptation to life in low-oxygen environments, and only a few show obvious links to the establishment and maintenance of the symbiosis with their *Desulfobacteraceae* symbionts. The fact that the *Desulfobacteraceae* are not found to be major LGT donors suggests that the phagocytic capacity of *Anaeramoeba* gives rise to more foreign genes than does the symbiosis itself; symbiosome-associated bacteria may only rarely be fully internalized and digested, thus providing few opportunities for gene transfer.

We have provided deep insight into the evolution of a unique, elaborate symbiotic organ in an anaerobic protist lineage. We have revealed some of the metabolic interactions and selective pressures that led to its establishment, and the evolutionary consequences on the genomes of the partner organisms. Future studies will focus on dissecting the fine-scale structure and function of the *Anaeramoeba* symbiosomes and host-symbiont interactions.

## Materials and methods

An expanded detailed account of the materials and methods can be found in the Supplementary text.

### Culturing and harvesting of Anaeramoeba

*A. flamelloides* BUSSELTON2 and SCHOONER1 and *A. ignava* BMAN cells were propagated as described in (Táborský, Pánek, and Čepička 2017) and cultures were scaled up and maintained as described in (Suppl. Text). Large-scale cultures of *Anaeramoeba* were harvested by decanting the culture supernatant and rinsing the amoeba monolayer in each flask. Cells were detached by cold-shock followed by percussive force to ensure efficient cell detachment. The resulting cell suspensions were centrifuged and cell pellets were processed further for RNA or DNA extraction (see Suppl. Text).

### RNA extraction and sequencing

*A. flamelloides* BUSSELTON2 and *A. ignava* BMAN cells were harvested as above and total RNA was extracted using TRIzol™ Reagent (Thermo Fisher Scientific) according to the manufacturer’s recommendations. Ten µg of total RNA was treated by the TURBO DNA-free™ Kit (Thermo Fisher Scientific) then treated with the DNase inactivation reagent. The total RNA was submitted to Génome Québec for sequencing. Libraries were made using the Illumina TruSeq RNA strand-specific sequencing kit and were sequenced on an Illumina HiSeq 4000 using 100 bp paired reads. Illumina reads were quality checked using FastQC v.0.11.5 (http://www.bioinformatics.babraham.ac.uk/projects/fastqc) and trimmed using Trimmomatic v0.36 (Bolger, Lohse, and Usadel 2014).

### Enrichment of prokaryotic mRNA and sequencing

Total RNA from enriched *A. ignava* BMAN and *A. flamelloides* BUSSELTON2 samples were treated by Terminator Exonuclease (Epicentre) according to the manufacturer’s instructions. After phenol extraction, ethanol precipitation and resuspension in distilled water, the total RNA samples were submitted to Génome Québec where polyA+ selection was performed to remove eukaryote mRNAs. One µl of supernatant from these RNA selections was used as input into the fragmentation step ahead of first-strand cDNA synthesis in the TruSeq protocol. The final library was sequenced on the Illumina NovaSeq 6000 S2 using 150 bp paired reads. Illumina reads were quality checked and trimmed as above.

### DNA extraction and short-read sequencing

Large-scale cultures of *Anaeramoeba* were harvested as described above and in Suppl. Text and DNA was purified using the MagAttract HMW gDNA kit (Qiagen) using the tissue lysis protocol. DNA extracts were further purified on the GenomicTip G/20 column (Qiagen) by the manufacturer’s protocol. Sample quality and quantity were assessed by agarose gel electrophoresis, UV specrophotometry and the Qubit™ dsDNA BR Assay Kit (Thermo Fisher Scientific).

DNA samples for Illumina short-read sequencing were submitted to Génome Québec for construction of shotgun and PCR-free shotgun libraries (see Table S11) using the Illumina TruSeq LT kit. The libraries were sequenced on an Illumina HiSeq X using 150 bp paired reads. Illumina reads were quality checked and trimmed as above.

### Long-read sequencing and basecalling

Genomic DNAs (1-5 µg) prepared as above were processed using Oxford Nanopore LSK108, LSK109 or LSK308 kits to construct sequencing libraries. The libraries were loaded on FLO-MIN106 (R9.4 or R9.4.1 pore) or FLO-MIN107 (R9.5 pore) flow cells and sequenced on the MinION Mk2 nanopore sequencer (Oxford Nanopore) running the MinKNOW control software. The fast5 files were basecalled to fastq format using Guppy v2.3.5 (Oxford Nanopore). The fastq reads were trimmed using Porechop v0.2.3_seqan2.1.1 (https://github.com/rrwick/Porechop) with the --discard_middle flag.

### Assembly and correction

The *A. ignava* BMAN read set was assembled using ABruijn (v1.0) (Lin et al. 2016) using default parameters. *A. flamelloides* BUSSELTON2 and SCHOONER1 read sets were assembled in metagenomics mode (--meta) using Flye (v2.4) with 3,000 bp min overlap and Flye (v2.4.2) (Kolmogorov et al. 2020) with 1,500 bp min overlap respectively.

Read mapping steps for Illumina short-reads were performed by bowtie2 (v2.3.1) (Langmead and Salzberg 2012) and nanopore reads were mapped by minimap2 (v2.10-r761) (H. Li 2018) with parameters (`-ax map-ont`).

The BMAN assembly was corrected by three rounds of Racon (v0.5) (Vaser et al. 2017) followed by Nanopolish v0.8.4 (nanopolish-git-dec-18-2017) (Loman, Quick, and Simpson 2015). The final BMAN assembly was obtained by two rounds of Pilon (v1.22) polishing employing (--mindepth 5 –fix bases) parameters (Walker et al. 2014).

The BUSSELTON2 and SCHOONER1 assemblies were corrected by four rounds of Racon (v1.4.13) with settings: `-u -m 8 -x -6 -g -8 -w 500`. Draft genomes were polished using Medaka v0.6.2 (https://github.com/nanoporetech/medaka) using the (`-m r941_trans`) model. The final assemblies were generated by five rounds of Pilon (v1.23) (Walker et al. 2014) polishing employing (--mindepth 1 --fix bases,amb) parameters.

### Genome classification and binning

Genome binning was performed manually utilizing the combined evidence of mapped polyA+ selected RNA sequencing data, sequence similarity searches (blastn and blastx) against the NCBI nr database, long-read coverage information, and GC-content of contigs. Chimeric and/or misassembles were identified by consistency of long-read mapping and split manually at the read-mapping border to retain the eukaryotic part of the contig. The presence of spliceosomal introns with GT-AG boundaries in genes was the main criterion for assigning a contig as being eukaryotic. Several analyses were used to confirm that a single isolate was present in the culture (see Suppl. Text).

RNAseq data for *A. ignava* BMAN and *A. flamelloides* BUSSELTON2 were mapped using Hisat2 v2.1.0 (Kim et al. 2019) using (`--rna-strandness RF --phred33 --max-intronlen 10000 -k 2`) flags. For *A. flamelloides* SCHOONER1, the BUSSELTON2 dataset was mapped with Hisat2 v2.1.0 (Kim et al. 2019) using relaxed parameters (`-- phred33 --max-intronlen 10000 -k 2 --mp 1,1 --sp 20`).

The GC% for each contig was calculated by `countgc.sh` in the BBMap package v38.20 (sourceforge.net/projects/bbmap/).

### Hybrid assembly of symbiont genomes

Long- and short sequence reads mapping to each *Desulfobacteraceae* genome were extracted and re-assembled using the hybrid-assembler Unicycler. For *A. flamelloides* SCHOONER1, the long-read data was assembled using Flye (v2.4.2 with --meta flag) (Kolmogorov et al. 2020). A detailed description of symbiont genome assemblies can be found in the Suppl. Text. ANI values were calculated using the OAT software (Lee et al. 2016).

### Prokaryotic annotation

The reassembled symbiont genomes were annotated by Rapid Annotation using Subsystem Technology v2.0 (RAST: (Aziz et al. 2008)) (Sym_BMAN; 2294.7, Sym_BUSS2, 2294.12; Sym_SCH1 2294.13). Insertion sequences were predicted using the ISsaga v2.0 webserver (http://issaga.biotoul.fr/ISsaga2/). Pseudogene candidates were identified using Pseudofinder v0.11 (Syberg-Olsen et al. 2021) and additionally refined by manual curation. Synteny was calculated using Sibelia v3.0.7 (Minkin et al. 2013). Circos plots were created with Circa (http://omgenomics.com/circa).

### 16S rDNA amplicon typing

The *Anaeramoeba*-enriched cell material was prepared by cold-shock and monolayer rinsing as described above. The supernatant samples were obtained by pelleting 10 mL of culture supernatant poured off prior to monolayer rinsing. The samples were extracted using the DNeasy PowerSoil Pro Kit (Qiagen) according to the manufacturer’s instructions.

The V4V5 (Bacteria) or V6V8 (Archaea) regions of the 16S rRNA genes were amplified and sequenced at the Integrated Microbiome Resource (IMR) at Dalhousie University, as described in (Comeau, Douglas, and Langille 2017). Bioinformatic processing of raw reads was carried by IMR (see (Comeau, Douglas, and Langille 2017)), updated in the Standard Operating Procedures (SOPs) for creation of ASV in QIIME2 (v2019.7) as outlined on the current MicrobiomeHelper website (https://github.com/LangilleLab/microbiome_helper/wiki).

### Phylogenomics

Phylogenomic analysis was performed on the symbiont genomes and genomes from *Desulfobacterales* (NCBI:txid213118) supplemented by data from outgroup taxa. The initial selection of representative genomes and marker genes / proteins was done using phyloSkeleton v1.1 (Guy 2017) from 353 *Desulfobacterales* genomes from NCBI. One representative genome per genus was selected in *Desulfobacteraceae* except for *Desulfobacter* where all available genomes were selected. The Bac109 protein markers were identified with phyloSkeleton v1.1 (Guy 2017) and only genomes with >80% of the marker genes were selected for further analyses. The UPF0081 marker was removed since it was only present in the out-group genome but not in any of the in-group genomes. Each of the 108 remaining marker proteins was aligned by MAFFT-linsi v7.458 (Katoh and Standley 2013) and trimmed by BMGE v1.12 (Criscuolo and Gribaldo 2010), using the BLOSUM30 matrix and stationary-based character trimming. Alignments were concatenated and a phylogenetic tree was inferred by IQTree v2.0.3 (Minh et al. 2020) using the LG4X (Le, Dang, and Gascuel 2012) substitution matrix with 1,000 ultrafast bootstraps (Hoang et al. 2018). The resulting tree was inspected, closely related taxa were removed, and the procedure was repeated. The final dataset consisted of 35 genome-derived sequences: *Desulfovibrio desulfuricans* ND132 (1 genome), *Desulfobacteraceae* (23 genomes), *Desulfobacter* (8 genomes) and the *Anaeramoeba* symbionts (3 genomes). A maximum-likelihood tree was inferred using IQ-TREE v2.0.3 (Minh et al. 2020) using the LG+C60+F+Gamma mixture model (Si Quang, Gascuel, and Lartillot 2008). The PMSF model (H.-C. Wang et al. 2018) generated from the above guide-tree and fitted model was used to perform 100 nonparametric bootstrap replicates.

### Protein-coding gene family expansions and contractions

We employed the PANTHER family database (v15) (Mi et al. 2017) to examine which protein families may have undergone expansions or contractions. Assignment of proteins to families was done with direct hidden Markov models (HMM) searches and the Panther Score tool (Mi et al. 2019) (set for hmmsearch with HMMER v3.1b2 (Eddy 2011)) using the PANTHER family database with e-value cut-off < 1e-5. Once classified, proteins assigned to each PANTHER family were counted to investigate relative family expansion or contraction in any *Anaeramoeba* species relative to other groups. (Protein families that were taxon-specific, exclusive to each group being compared, apparently heavily expanded in diplomonads, *Trichomonas, Dictyostelium* or *Naegleria,* and those with median counts of zero for any compared group were not considered in the expansion/contraction analysis.) To detect signatures of expansion/contraction, outlier values within each group were replaced by the median value of their respective group when the value had a 1.5 > z-score < −1.5. Data were normalized by the median value of their respective families and log_2_FC among groups was calculated. To identify what metabolic pathways were relevant for further analyses, we investigated families with signature of expansions or contractions ≥ 5-fold (2.35 > log_2_FC or log_2_FC < −2.35). Relative contractions and expansions are reported for *Anaeramoeba* taxa and protein families were considered ‘core’ if they were present in all studied proteomes and ‘accessory’ if they were missing in at least one. Possible expansions/contractions were visualized with Circos (Krzywinski et al. 2009). A large proportion of the protein families expanded beyond ≥ 5-fold belong to RNA synthesis, DNA synthesis and repair, and membrane trafficking metabolisms. Hence, we carried out pathway reconstruction for RNA synthesis and membrane trafficking (see below) by validating protein orthology and recording absence/presence patterns. In the case of RNA systems, we first classified query proteins of interest (from yeast or human) with PANTHER Score (as above) to identify their PANTHER family. Query proteins and proteins identified for each PANTHER family of interest were then aligned together with a random taxonomic sample from the same family of interest from the PANTHER database. Alignments were trimmed to carry out phylogenetic reconstructions as described in (Salas-Leiva et al. 2021).

For membrane-trafficking proteins, *Anaeramoeba* datasets were searched by BLAST using queries from previous studies (Elias et al. 2012; Gabernet-Castello et al. 2013; Klinger, Klute, and Dacks 2013). Identified Rab and TBC proteins were subjected to orthologous clustering by OrthoFinder v2.0.0 (Emms and Kelly 2019) including proteins from human, cnidarian *Nematostella vectensis*, and the heterolobosean *Naegleria gruberi*. Phylogenetic analyses for selected Rab sub-families and TBC proteins were conducted. Sequences were aligned by MAFFT v7.458 (Katoh and Standley 2013) under L-INS-i strategy, and poorly aligned positions were removed by trimAl v1.4.rev15 (Capella-Gutierrez, Silla-Martinez, and Gabaldon 2009) using -gt 0.8. Maximum-likelihood trees were inferred by IQ-TREE v1.6.8 (Nguyen et al. 2015) using the PMSF method (H.-C. Wang et al. 2018) and the LG+C20+F+G model, with the guide tree inferred under the LG+F+G model. UFBOOT branch supports were obtained with 1,000 replicates.

### Lateral gene transfer in Anaeramoeba

In order to assess LGT from prokaryotic and viral donors, we conducted a large-scale screen of the predicted proteomes of the three *Anaeramoeba* genomes. Initial clustering was performed using Orthofinder v2.5.4 (Emms and Kelly 2019) with the following outgroup proteomes: *Arabidopsis thaliana*, *Dictyostelium discoideum*, *Acanthamoeba castellanii* str. Neff, *Giardia intestinalis*, *Homo sapiens*, *Kipferlia bialata*, *Monocercomonoides* sp., *Naegleria gruberi*, *Trepomonas* PC1, *Trypanosoma brucei*, *Trichomonas vaginalis*, *Saccharomyces cerevisiae*. In some instances, Orthofinder appeared to cluster proteins that were assigned different PANTHER annotations obtained in the Protein-coding gene family expansions and contractions analysis, indicating they were unlikely to be true orthologs. In such cases, clusters were further split according to the PANTHER annotations.

Proteins for phylogenetic trees were inferred by gathering homologous sequences in the nr database using blastp using an E-value cutoff of 1×10^-5^. All *Anaeramoeba* orthologs/paralogs constituting a cluster together with the 500 best database hits to each were grouped. Clusters containing at least one prokaryotic or viral protein together with *Anaeramoeba* orthologs/paralogs were aligned using MAFFT v7.310 (Katoh and Standley 2013) with the default setting and sites were selected using BMGE v1.0 (Criscuolo and Gribaldo 2010). Initial phylogenetic reconstructions were done using FastTree v1.0.1 (Price, Dehal, and Arkin 2009). These phylogenies were used to reduce the taxonomic redundancy of the initial sequence files using in-house scripts. The reduced files were realigned using MAFFT v7.310 (Katoh and Standley 2013) with the accurate option (L-INS-i), and sites were selected using BMGE v1.0 (Criscuolo and Gribaldo 2010). Phylogenetic reconstructions were done using IQ-TREE multicore v1.5.5 (Nguyen et al. 2015) with the LG4X model and 1000 ultrafast bootstraps (UFBOOT) for alignments ≥80 sites (shorter alignments were treated separately, see below). Phylogenetic trees were then parsed to find putative LGTs, using the following criteria:

1. When *Anaeramoeba* sequence(s) and prokaryotic or viral sequences constituted a clade supported by an UFBOOT value of ≥70% the *Anaeramoeba* homologs were considered as possible LGTs. To allow for mis-annotation, one other sequence in that clade could be eukaryotic.
2. When no clades were well-supported, if ≥95% of the sequences in the tree were prokaryotic and/or viral, then this was counted as an LGT.
3. The taxonomic group of the donor could be inferred when the >50% of the taxa in a clade containing the *Anaeramoeba* protein(s) were from a particular taxonomic group.
4. For alignments <80 sites, no trees were constructed. However, when all sequences constituting the cluster, besides *Anaeramoeba*, were prokaryotic or viral, it was counted as LGT.

### FISH probe design and testing

The symbiont 16S rDNA sequences were extracted from the RAST annotations and aligned by MAFFT v7 (https://mafft.cbrc.jp/alignment/software/) to selected full-length 16S rDNA sequences selected from the Ribosomal Database Project (RDP).

Probes targeting Sym_BUSS2 and Sym_SCH1 to the exclusion of outgroups in the 16S rDNA alignment were designed using Decipher (Wright et al. 2014). The probes were evaluated by matching against the SILVA and RDP databases and did not match any other sequences in the database at 1 mismatch. The probes were ordered from biomers.net GmBH. Probe sequences, fluorophores and hybridization conditions can be found in Table S12.

### Fluorescence in situ hybridization (FISH)

*Anaeramoeba* cell cultures in 10 mL slanted culture tubes were decanted and the adhered cells were resuspended in the remaining volume of culture media (typically 200 µl) by percussive shock. The suspended cells were applied to Teflon multi-well microscope slides (Electron Microscopy Sciences (ER-264)) for 5 min at room-temperature. The cells were fixed for 10 min at room temperature by adding a suitable volume of 16% methanol-free formaldehyde (w/v) (Pierce-Thermo Fisher Scientific, Cat No 28908) to give a final concentration of 4% paraformaldehyde in ASW. The cells were rinsed in 2×50 mL distilled water for a total of 3 min and then air-dried. The slide was immersed in 100% ethanol for 5 min and air-dried again. Hybridization buffer (20 mM Tris-HCl, pH 7.6, 0.01% SDS, 900 mM NaCl) with an appropriate concentration of formamide and probe (5 ng/µl) (Table 12) was added to dried cells and incubated for 2-3 hours at 47 °C in a moisture chamber. Post-hybridization, the slides were rinsed with buffer (20 mM Tris-HCl, pH 7.6, 0.01% SDS, 5 mM EDTA and an appropriate NaCl concentration depending on the %FA in the hybridization buffer; see Table S12) and incubated with 50 mL for 45 min at 48 °C, finally rinsed in water for 40 s and air-dried. The slides were mounted in ProLong™ Diamond Antifade Mountant with DAPI (Thermo Fisher Scientific, Cat No P36971) using #1.5H cover slips (Ibidi, Cat No 10811). The slides were incubated at room temperature ≥12 hours before imaging.

Images from the slides were acquired using either wide-field microscopy on the Zeiss Axio Imager Z2 or by confocal microscopy on Zeiss LSM 710 or Leica SP8. Pseudo color, merging of channels and projections were made in Zeiss ZEN v3.1 or BioImageXD (Kankaanpää et al. 2012).

### Scanning electron microscopy

*A. flamelloides* BUSSELTON2 cells were cultured and harvested as described above with cells finally being resuspended in 500 µL of ice-cold filtered growth media. 50 µL of cells were transferred onto poly-L-lysine-coated 12 mm round coverslips, left to adhere for 5 min at room temperature and immediately fixed with a drop of 25% glutaraldehyde and OsO_4_ vapor for 1 hour. After fixation, the coverslips were washed three times in filtered ASW, and then subjected to a dehydration series of ethanol-water mixtures, as follows: 30%, 50%, 70%, 80%, 90%, 95%, 100% (three times). This was followed by critical-point drying with CO_2_ on a Leica EM CPD300, then a ∼15 nm Au-Pd coat was added with a Leica EM ACE200 sputter-coater. Samples were imaged on a Hitachi S4700 scanning electron microscope.

### Transmission electron microscopy

Fixation (both chemical and cryofixation), embedding, sectioning and TEM was conducted as described in detail by (Táborský, Pánek, and Čepička 2017).

### Tubulin staining

For tubulin staining the cells were harvested and fixed for 10 min in 4% paraformaldehyde according to the procedure described in the ‘Fluorescence in situ hybridization (FISH)’ section. The fixative was removed, and the cells were washed twice with ASW and once with PBS. Cells were then incubated 15 min in PBS with 50 mM NH_4_Cl to quench remaining aldehyde fixative. The cells were washed twice with PBS and permeabilized for 15 min in PBS with 0.1% Triton X100. Cells were blocked in antibody dilution buffer (ADB - 1% BSA-c (Aurion) in PBS with 0.1%Triton X100) for 1 hour at room temperature. The cells were then incubated with primary antibodies diluted in ADB (TAT1; 1:200 dilution or KMX-1; 1:200 dilution) overnight at 4 °C in a moisture chamber. The slides were droplet-washed six times using excess ADB and incubated 1 hour at room temperature with secondary antibody (goat anti-mouse Alexa Fluor 594, 1:250 dilution, Thermo Fisher Scientific, Cat No A-11032). The cells were washed six times using ADB, twice with PBS, and then mounted in ProLong™ Diamond Antifade Mountant with DAPI (Thermo Fisher Scientific, Cat No P36971) using #1.5H cover slips (Ibidi, Cat No 10811).

Images were acquired using wide-field microscopy on the Zeiss Axio Imager Z2 or by confocal microscopy on Zeiss LSM 710 or Leica SP8. Pseudo color, merging of channels and projections were made in Zeiss ZEN or BioImageXD (Kankaanpää et al. 2012).

### Wheat Germ Agglutinin (WGA) staining

For WGA staining two different protocols were used, the first was a live stain protocol, and the second relied on staining after aldehyde fixation. In both protocols, cells were harvested and attached to multi-well slides according to the FISH procedure (above). Live stain cells were washed twice using 100 µl ASW and then incubated for 10 min in 50 µg/ml WGA-CF633 conjugate (Biotium, Cat No #29024-1) in ASW. The stain was removed, and cells were washed twice in 100 µl ASW and then post-fixed in 4% formaldehyde in ASW for 10 min at room-temperature.

For the aldehyde fixation protocol, cells were fixed in 4% formaldehyde in ASW immediately after attachment and the first two washes. The fixative was removed, and the cells were washed twice in 100 µl ASW. Each slide was rinsed twice in a large volume of distilled water for 1 min 30 s each and air-dried. The slides were incubated 5 min in 100% ethanol, air-dried and the slide was mounted and imaged according to the procedure described in the FISH section.

## Data availability

Sequencing reads and the annotated genomes of *A. flamelloides* BUSSELTON2, *A. flamelloides* SCHOONER1, *A. ignava* BMAN, Sym_BUSS2, Sym_SCH1 and Sym_BMAN were deposited to NCBI under the BioProject number PRJNA634776.

## Supporting information

Table S12: FISH probes

Table S13: Defense systems in Desulfobacteraceae

Table S14: Amino acid pathways and unique proteins in the symbionts

Table S15: Symbiont DNA repair

Table S1-S3: Sequencing data and metrics, Symbiont genome characteristics, Introns statistics

Table S4: Pseudogenes in Sym_BUSS2 and Sym_SCH1

Table S5: IS elements in the symbionts

Table S6: Metatranscriptomics of Sym_BUSS2 and Sym_BMAN

Table S7: Gene families in Anaeramoeba

Table S8: LGT analysis

Table S9: Anti-SMASH predictions for Anaeramoeba predicted proteomes

Table S10: Summary table of membrane-trafficking components Anaeramoeba

Table S11: Sequencing libraries

## Acknowledgements

Gordon Lax and Yana Eglit are acknowledged for help during SEM samples preparation and imaging. The majority of the work and J.J.H. were supported by a Foundation grant (FRN-142349) from the Canadian Institutes of Health Research awarded to A.J.R. J.J.H was additionally supported a grant from Vetenskapsrådet, (VR-NT grant 2022-04490). Archibald Lab contributions to this study were supported by the Gordon and Betty Moore Foundation (GBMF5782) and a Discovery Grant from the Natural Sciences and Engineering Research Council of Canada (RGPIN-2019-05058). Work from the Čepička lab was funded by Czech Science Foundation grant no. 21-30563S. Computational resources were supplied by the project “e-Infrastruktura CZ” (e-INFRA CZ LM2023054) supported by the Ministry of Education, Youth and Sports of the Czech Republic.

