## Supplemental text for "A unique symbiosome in an anaerobic single-celled eukaryote"

**Supplementary text**

**Supplementary Materials and Methods**

*Cultivation*

*A. flamelloides* BUSSELTON2 and SCHOONER1 and *A. ignava* BMAN cells were maintained in 1x ATCC medium: 1525 Seawater 802 (SW802) medium as in (Táborský, Pánek, and Čepička 2017)). Cultivation was done in capped tubes (not under strict anoxic conditions).

Large-scale cultures of *Anaeramoeba* were performed in 0.25x SW802 media in fully filled 550 mL Cellstar® T175 low profile flasks (Grenier Bio-One GmbH). Cells were dislodged with a cell scraper and split 1:1 with fresh media every 2-4 days for *A. flamelloides* BUSSELTON2 and SCHOONER1 or 7-10 days for *A. ignava* BMAN.

### Large-scale cell culturing and harvesting

Large-scale cultures of *Anaeramoeba* were harvested by decanting the culture supernatant and rinsing the amoeba monolayer in each flask with 250 mL 1x Artificial Seawater (ASW) (per liter medium: 24.72 g NaCl, 0.67 g KCl, 1.364 g CaCl_2_×2H_2_O, 4.66 g MgCl_2_×6H_2_O, 6.29 g MgSO_4_×7H_2_O, 0.18 g NaHCO_3_-). Cells were detached by cold-shock by adding 35 mL of ice-cold 1x ASW and immersing the flasks in ice-slush for 15 min. The flasks were inspected for cell detachment and percussive force was used to ensure efficient cell detachment. The cell suspensions were centrifuged at 500×g for 8 min at 4 °C and the combined cell pellets were resuspended in 10 mL ice-cold 1x ASW. Centrifugation was repeated as above, and the cell pellet was processed further for RNA or DNA extraction.

*Confirming that only a single Anaeramoeba isolate was in each genome assembly.*

Firstly, unique 18S rRNA and 28S rRNA sequences of each *Anaeramoeba* genome were retrieved with Metaxa2 (Bengtsson-Palme et al. 2015). Secondly, the up/down stream areas of highly amplified RP gene families were investigated to rule out the presence of multiple highly similar contigs that could be due to mixture of closely related strains in the culture. Both of these analyses suggested that there was only a single *Anaeramoeba* isolate in each genome assembly.

*Enrichment of amoebae*

16S rDNA sequencing data of replicate enrichments confirmed the metagenomic sequencing results with distinct profiles in the culture supernatant and the amoeba-enriched cell material (Figure S3B,C). *Bacteroidales and Pseudoalteromonadaceae* were dominant in the culture supernatant of *A. ignava,* whereas *Vibrio* was most abundant in the *A. flamelloides* sample. These lineages were depleted in the amoeba-enriched samples (Figure S3B,C). The obtained relative level of enrichment for the *Desulfobacteraceae* lineage was higher in *A. flamelloides* (Figure S3B) than in *A. ignava* (Figure S3C). We mapped long-reads from individual sequencing runs representing distinct DNA preparations back to the assembled contigs to estimate the respective abundance of each organism from the average contig depth coverage. The data was normalized to the coverage of the *Anaeramoeba* nuclear genome to establish the number of prokaryotic cells / amoeba genome equivalents (Figure S17).

*IS element activity*

The number of IS elements recorded in SRBs is variable, but some free-living *Desulfobacteraceae* genomes encode high numbers (for example, 130 copies in *Desulfobacula toluolica* Tol2). We found high numbers of transposable elements in both of the *A. flamelloides* symbionts (> 700 copies/genome) (Table S5), while the BMAN symbiont encodes 60 IS elements, a comparable number to that of free-living species. The IS elements showed no strong evidence of clustering overall (Figure S7A,B) but we found that >200 IS elements were close (<1500 bp) to edges of syntenic blocks, indicating the frequent genome rearrangements might indeed be connected to IS element activity (Figure S7C,D). Many of the high-copy-number IS elements had a complete transposase indicating that they were recently active. The two most abundant IS elements in both genomes were IS256 (Sym_BUSS2 172 copies, Sym_SCH1 194 copies) and IS4_ssgr_IS4Sa (Sym_BUSS2 129 copies, Sym_SCH1 185 copies). In total we detected 13 families in Sym_SCH1 and 14 families in Sym_BUSS2, and about half of them, six families in Sym_SCH1 and seven families in Sym_BUSS2, were considered abundant (>40 copies) (Table S5).

*Pseudogene curation in A. flamelloides symbionts*

Automatic methods for pseudogene detection (PGAP and pseudofinder) both indicated an elevated number of pseudogene candidates in the Sym_BUSS2 and Sym_SCH1 genomes compared to Sym_BMAN (Table S2), however the detected numbers were starkly different between the two methods. Pseudogenes were manually curated for Sym_BUSS2 and Sym_SCH1 guided by synteny evidence and the automatic pseudogene predictions.

*A. flamelloides symbionts have lost flagella and have impaired cell wall assembly machinery*

Flagella are crucial to many cells found in fluctuating environments but may become dispensable in stable environments encountered by symbionts (Toft and Fares 2008). Sym_BMAN shows no degradation of its flagellar operon with 54 full-length genes and no IS element insertions. Strikingly Sym_SCH1 and Sym_BUSS2 both have a ten gene deletion that includes (*flgB*, *flgC*, *fliE*, *fliF*) and a 20 kb inversion at the 3’end compared to Sym_BMAN. Their flagellar loci are further degraded by pseudogenization (Sym_BUSS2, 14 Ψ / 44 genes; Sym_SCH1, 7 Ψ / 22 genes) and multiple IS element insertions (Sym_BUSS2, 4 insertions; Sym_SCH1, 3 insertions). In addition, Sym_SCH1 has a 27 kbp deletion that deletes 24 genes (Figure S9). We conclude that both *A. flamelloides* symbionts have lost the ability to produce flagella, whereas the *A. ignava* symbiont still retains the capacity to produce flagella.

The three symbionts retain the machinery to produce core lipid A and Kdo. However, Sym_BUSS2 and Sym_SCH1 have pseudogenes of *waaL* that encode O-antigen ligase. Sym_BUSS2 further has a defunct copy of *wzyC*, which encodes O-antigen polymerase. *A. flamelloides* symbionts also might be impaired in decorating lipid A with O-antigen. Having reduced O-antigen on lipid A decrease the permeability barrier at the outer membrane (Whitfield, Williams, and Kelly 2020), and it could additionally lead to better adherence between host and symbiont.

*Adherence*

Type IV pili help pathogens such as *Pseudomonas aeruginosa* to colonize a host. The *pilMNOPQ* loci encode factors important for both T4P assembly and twitching motility. The Sym_BMAN and *D. toluolica* loci are syntenic and show no evidence of degradation. The Sym_BUSS2 and Sym_SCH1 loci both have a pseudogenized *pilQ* gene. The gene is frame-shifted in both strains, with Sym_SCH1 having a further two IS element insertions. Further, Sym_BUSS2 and Sym_SCH1 have pseudogenized copies of *pilB* and one of the two copies of *pilU/T*. Sym_BMAN further encodes an incomplete Tad (tight adherence) gene cluster which is not present in Sym_BUSS2 and Sym_SCH1. This gene cluster has shown to be essential for colonization of surfaces by the human pathogen *Actinobacillus actinomycetemcomitans* (Planet et al. 2003). Components of the Tad cluster code for a secretion system that exports and assembles bundled Flp pili (fibrils).

*DNA repair systems of the symbiont*

The genome degeneration in the *A. flamelloides* symbionts could indicate ineffective DNA repair systems. We failed to find genes encoding endonuclease IV that binds to and excises apurinic-apyrimidinic sites and excises them to direct DNA repair in both *A. flamelloides* symbionts. This protein is found in Sym_BMAN and free-living members of *Desulfobacteraceae* (Table S15) We also found pseudogenized copies of deoxyribodipyrimidine photo-lyase in the *A. flamelloides* symbionts making them less able to respond to UV-induced stress. However, this gene appears to be missing from other *Desulfobacteraceae* genomes including Sym_BMAN (Table S15). Several genes whose products are prominent members in DNA replication and repair pathways (*dinB*, *uvrD, and dnaE*) were found to be pseudogenized in Sym_BUSS2 and Sym_SCH1 (Table S4). The significance of these gene losses is not clear since most of these are multi-copy genes, and their function is likely to be compensated by the presence of a second redundant copy. Mutator phenotypes have been recovered in natural settings as well as evolution experiments where they can have a selective advantage to find adaptive mutations under relaxed selective conditions. Mutators are predicted to be favored during nascent symbioses where the chance of essential genes being perturbed is lower than in later stages in symbiont evolution (McCutcheon, Boyd, and Dale 2019).

*The A. ignava symbionts encode type VI secretion systems and anti-viral defenses*

We detected the presence of several potential protein secretion systems in the symbionts. All three are predicted to encode the Sec-SRP general secretion system while the twin-arginine translocation (TAT) pathway appears compromised in Sym_BUSS2 and Sym_SCH1 by the pseudogenization of *tatB*. Sym_BMAN encodes a Type II secretion and a type VI secretion system (T6SS) where the latter consists of two convergent transcription units of 25 genes, with most genes showing evidence of expression (Figure S10). The first transcription unit includes 12 genes (*tssJKLM,* *tagF*, *tssA*, *tssBC*, *tssEFGH*) that are conserved in most T6SS and two hypothetical proteins. The second transcription unit encodes the two conserved T6SS factors, *vgrG*, *hcp,* and upstream of these; we find a forkhead associated domain (FHA) protein, a TPR repeat protein, a carbohydrate esterase 4 (CE4) superfamily protein. The downstream genes include a DUF4280 family protein previously suggested as potential spike (PAAR) proteins (Lays, Tannier, and Henry 2016). Directly downstream of *vgrG,* we identified a DUF4123 protein, which is generally encoded upstream of putative T6SS effectors (Liang et al. 2015). The four effector candidates encoded between the DUF4123 domain protein and DUF4280 domain protein have no known function, but putative homologs are present in both free-living and bacterial symbiont genomes.

Phage and plasmid infections can be controlled by a growing list of defense systems and multiple systems are usually found in bacterial and archaeal genomes (Payne et al. 2022). We investigated the repertoire of phage and plasmid defense systems in the symbionts and compared it to repertoires of free-living *Desulfobacteraceae* (same set of organisms as used in the phylogenetic analysis in Figure 2A) using the tool PADLOC (Payne et al. 2022). We detected very few defense components (Sym_SCH1 – 3 systems, Sym_BUSS2 – 2 systems) in the *A. flamelloides* symbionts (Table S13). This is in contrast to most free-living *Desulfobacteraceae* which tend to encode multiple defense systems (average 13.9, MAGS excluded 14.3). Upon close inspection two each of the detected systems of Sym_BUSS2 and Sym_SCH1 encode pseudogenized CRISPR systems of type I-C. The leading *cas2* and terminating *cas3’’* genes are truncated, and no associated CRISPR protospacer array was identified. Sym_BMAN appears more similar to free-living *Desulfobacteraceae* in that it encodes 9 defense systems including an intact type I-F CRISPR system with an array of 25 spacers. None of the spacers were found to be self-matching or belong to known phages or plasmids.

*Membrane complexes and transporters*

Several highly conserved membrane complexes direct electron transfers in DSR organisms (Pereira 2008). In Sym_BUSS2, we detected high expression (dsrJ – 768 RPKM, dsrO – 751 RPKM, dsrP – 633 RPKM, dsrK – 536 RPKM, dsrM – 390 RPKM) of the DsrMJKOP transmembrane electron carrier complex that shuttles electrons to sulfite reductase. By contrast, this membrane complex is not as prominently expressed in Sym_BMAN (dsrJ – 136 RPKM, dsrO – 109 RPKM, dsrP – 107 RPKM, dsrK – 77 RPKM, dsrM – 136 RPKM). The Rnf respiratory membrane complex is highly expressed in both symbionts. This complex acts a H+/Na+ channel that oxidizes reduced ferredoxin and reduces NAD+ coupled to ion transport. The most highly expressed gene in both Sym_BMAN and Sym_BUSS2 is an outer membrane porin of unknown function. The outer membrane protein OmpA is particularly highly expressed in Sym_BUSS2. In contrast, an Oxalate:formate antiporter is the second highest expressed transporter in Sym_BMAN. Additional transporters with prominent expression include nickel-, ammonium (Sym_BUSS2) and molybdenum transporters (Table S6).

*Amino acids metabolism in the symbionts*

All the three symbionts, Sym_BMAN, Sym_BUSS2 and Sym_SCH1, maintain the ability to synthesize the 20 standard amino acids. Complete pathways were predicted for 16 amino acids (L-arginine, L-asparagine, L-cysteine, L-glutamine, L-glycine, L-isoleucine, L-leucine, L-lysine, L-methionine, L-phenylalanine, L-proline, L-serine, L-threonine, L-tryptophan, L-tyrosine, L-valine) as well as chorismate biosynthesis (Table S15). Histidine synthesis contains two gaps where hisI and hisN activities are unaccounted for. These are known pathway gaps known to be present in free-living *Desulfobacteraceae* species (*Desulfobacter vibrioformis* DSM 8776, *Desulfobacula toluolica* Tol2) known to perform L-histidine biosynthesis. L-alanine can be synthesized from cysteine, L-lysine is made using the diaminopimelate-aminotransferase variant pathway, L-proline can be made from glutamate or ornithine and L-glycine from threonine and serine. Isoleucine biosynthesis is also possible via ferredoxin-dependent reductive carboxylation of propanoyl-CoA to 2-oxobutanoate propanoate. Multiple transaminases of different classes are present to likely serve the need for synthesis of L-alanine, L-aspartate and L-glutamate. There are some symbiont-specific differences in the amino acid metabolism;

The *A. flamelloides* symbionts differ in their amino acid pathways. For example, 8 out of 29 Sym_SCH1 and 5 out of 37 Sym_BUSS2 specific proteins respectively function in amino acid pathways (Table S15). Sym_SCH1 encodes several additional enzymes involved in Methionine biosynthesis (Homoserine O-acetyltransferase (EC 2.3.1.31), O-acetylhomoserine sulfhydrylase (EC 2.5.1.49), O-succinylhomoserine sulfhydrylase (EC 2.5.1.48). The enzyme complement of the arginine and ornithine degradation pathway is also variable between the two symbionts. Sym_BUSS2 and Sym_SCH1 can assimilate ammonia via glutamate-ammonia ligase and glutamate synthase. Sym_BUSS2 encodes threonine hydratase to convert threonine to isoleucine via 2-oxobutanoate. Sym_BUSS2 and Sym_SCH1 encode the L-arginine degradation pathway that proceeds via arginase.

The amino acid metabolic pathways are more extensive in Sym_BMAN compared to the *A. flamelloides* symbionts. Sym_BMAN encodes 36 and 37 additional genes in amino acid biosynthesis pathways compared to Sym_SCH1 and Sym_BUSS2 respectively (Supplemental table). For example, this includes expanded capabilities in alanine, glycine, arginine and polyamine biosynthesis. They also encode a bigger set of genes in ammonia assimilation via glutamate and aspartate as well as in proline uptake. Sym_BMAN can make L-asparagine from aspartic acid and L-homocysteine from L-homoserine. Genes for utilization of histidine as well as arginine and ornithine are also present. We detected inactivated copies histidine ammonia-lyase and urocanate hydratase, involved in histidine catabolism, in Sym_BUSS2 and Sym_SCH1.

*Selenocysteine and pyrrolysine*

Selenoproteins are well represented in organisms inhabiting certain anaerobic environments where their unique properties can be advantageous (Lacourciere and Stadtman 1999; Arnér 2010; Reich and Hondal 2016). Selenocysteine might be better able to withstand oxidation better than cysteine and could offer an advantage in fluctuating anaerobic environments (Maroney and Hondal 2018). The sulfate-oxidizing and sulfate-reducing symbionts of the gutless worm *Olavius algarvensis* was previously shown to encode a large complement of both selenoproteins and pyrrolysine-containing proteins (Zhang and Gladyshev 2007). The *Anaeramoeba* symbiont genomes encode many selenoproteins (BUSS2 – 19, Sch – 20, BMAN – 25). Selenocysteine is predicted to be synthesized from Se^0^ and incorporated in proteins in all three *Anaeramoeba* symbionts using the standard bacterial pathway. Enzymes predicted to use selenocysteine includes Selenide, water dikinase and prominent redox-proteins (CoB--CoM-reducing hydrogenase (Sec) delta subunit, DsrK-like, NAD-dependent formate dehydrogenase alpha subunit, Uptake hydrogenase large subunit).

Pyrrolysine is known as the 22nd amino acid that can be incorporated into proteins during translation. Sym_BMAN encodes the necessary enzymatic pathway (proline 2-methylase (pylB), pyrrolysine synthetase (pylC), proline reductase (pylD), pyrrolysine-tRNA(Pyl) ligase large and small subunits) to synthesize and incorporate pyrrolysine into proteins. Trimethylamine--corrinoid protein Co-methyltransferase (MttB) was found to be have an in-frame stopcodon to guide Pyl-insertion. The target for MttB-directed methylation in Sym_BMAN is unclear since no clear target is present.

*Nitrogenase*

Nitrogen sources are often limited in microbial systems even though molecular nitrogen is abundant in the atmosphere. A limited number of microbes have the ability to use molecular nitrogen as a nitrogen source using an enzyme called nitrogenase. The exotic molybdenum-iron cluster in nitrogenase is able to fix nitrogen directly into ammonia by ATP expenditure. Sym_BMAN encodes a Mo-nitrogenase and associated genes (nifH, nifD, nifN, nifB, nifV, 2Fe) that allows the fixation of N_2_ into ammonia using with the expenditure of ATP.

*Genome erosion buffering and symbiont stability*

GroEL/S is believed to buffer the deleterious changes that accrue in symbiont proteins as a result of genome erosion. Constitutive overexpression of GroEL/S is regarded as a critical factor in stabilizing endosymbionts in a wide range of insect symbioses (Kupper et al. 2014) and has been shown to buffer mutations in mutator strains of *E. coli* (Fares et al. 2002). In organisms experiencing genome erosion such as *Buchnera aphidicola* GroEL/S can amount to 10% of the total cellular protein (Baumann, Baumann, and Clark 1996). Interestingly we found that GroEL/S are highly expressed (Ranked 24 – 2044 RPKM and 39 – 1353 RPKM) in Sym_BUSS2 while moderately expressed in Sym_BMAN (Ranked 436 – 167 RPKM and 572 – 133 RPKM) in line with their respective degrees of genome erosion. In addition to this, the chaperone DnaK in Sym_BUSS2 is highly expressed (rank 93 – 663 RPKM) whereas it being expressed at a low level (rank 1734 - 43 RPKM) in Sym_BMAN.

We found additional cellular defense systems in the symbionts to be highly activated. Superoxide reductase and Rubredoxin defends the cell against highly reactive and toxic superoxide radicals (O_2_^-^) that is converted into hydrogen peroxide. Both genes are highly expressed in Sym_BUSS2 (rank 27 – 1959 RPKM and rank 66 – 524 RPKM respectively) and in Sym_BMAN (rank 27 – 1386 RPKM and 110 – 496 RPKM). The less toxic hydrogen peroxide is detoxified by the action of Rubrerythrin. The symbionts encode several copies of Rubrerythrin (8 copies in Sym_BMAN and 6 copies in Sym_BUSS2). Several rubrerythrin genes in Sym_BUSS2 show high expression and the most highly expressed rubrerythrin gene copy (rank 25 – 2043 RPKM) is transcribed from the same operon as the Per2 peroxide stress regulator which is likewise highly expressed (rank 84 – 690 RPKM). In addition, Sym_BMAN show high expression of Dodecin (rank 12 – 2136 RPKM) that can neutralize flavins and shield the cell from high flavin reactivity that can be toxic if they accumulate in an uncontrolled way. Several cold-shock proteins and additional stress sensors such as Ribosome hibernation protein YhbH, the previously mentioned Per2 peroxide stress regulator and Phage shock protein A are highly expressed. The abundant expression of cold-shock proteins might be related to the low temperatures the cells are exposed to during cell harvest. The abundant expression of stress coping genes indicates that the symbionts, and especially Sym_BUSS2, is highly active in countering stress perhaps related to accumulation slightly deleterious protein coding alleles.

*Introns in Anaeramoeba*

Among the 3 genomes analyzed, SCHOONER1 and BUSSELTON2 host genomes have a very similar intron complement in terms of the number of introns overall (Table S3A) and the size ranges (Table S3, Figure S16). The genomic context is equally similar with regard to the percentage of the genome devoted to introns, the GC% of all the introns and the number of introns per gene and per genomic 1 kb. The BMAN genome has almost as many introns (Table S3A) as SCHOONER1 and BUSSELTON2 (32,108, 39,247, 39,831 respectively) but apart from this the characteristics diverge. BMAN introns tend to be much smaller with a mode length of 76 versus 113 for SCHOONER1 and 124 for BUSSELTON2 and the longest intron found in BMAN (4168 bps) is less than half of the longest ones in the other genomes (Table S3A, Figure S16). Equally striking is the GC% of the introns. While SCHOONER1, BUSSELTON2 and BMAN all have very GC% poor genomes (23.78, 23.63 and 19.36 respectively), the introns in BMAN are even more GC% poor averaging only 8.14 GC% which is less than half of the genome globally while the GC percentage of introns in SCHOONER1 and BUSSELTON2 are close to their overall genomes (Table S3A). The AT richness of all three genomes is exemplified in the bases immediately before and after the splice sites. The surrounding bases are heavily weighted towards As and Ts (Figure S16D). Because the BMAN genome is 1/6^th^ the size of the other two genomes while having almost as many introns, its number of introns per kb is considerably higher (0.74 versus 0.15). The smaller genome size of BMAN, while having almost as many introns, also accounts for why the percentage of intronic sequence in all three genomes is very similar.

The number of introns that possess non-standard splice sites (5’ GT, 3’ AG) in the three genomes is extremely small but also in line with other sequenced genomes (). Only 55 of 32108 (0.17%) introns in BMAN had noncanonical boundaries while in SCHOONER1 and BUSSELTON2 the numbers were even lower, 6 and 7 respectively (Table S3B). All of the noncanonical intron boundaries in SCHOONER1 and BUSSELTON2 were of the GC-AG type while in BMAN 33 introns had AT-AC splice sites. Traditionally, AT-AC boundaries were considered diagnostic for U12 type introns. However, this is no longer the case and whether an intron is spliced by the major or minor (U12 type) spliceosome is determined by analyzing the branch point and critically, the first few bases in the intron (Turunen et al. 2013). U12 type introns have a highly conserved motif at the 5’ end, starting with the splice site. The first base can be either an A or a G followed by TATCCTTT. BMAN has 32 introns that start with (G/A)TATCCTTT (Table S3B) with 27 of them being AT-AC introns while in SCHOONER1 all 10 of the introns that start with the conserved motif are GT-AG introns as are the 8 in BUSSELTON2 that have the motif. The presence of the distinctive U12 5’ motif in this limited set of introns strongly suggests that this lineage has a minor spliceosome. However, there was no conserved branch point sequence among these potential U12 type introns. None of the branch points were similar to the reported conserved branch point motif (Turunen et al. 2013). However, it should be noted that the published branch point motif is based on a very limited set of introns from model organisms. The fact that *Anaeramoebae* share the 5’ motif with the model organisms but not the branch point motif suggests that only the 5’ motif is truly diagnostic for U12 type introns.

*Extended analyses of selected LGT candidates*

Stairs et al. (Stairs et al. 2021) used transcriptomics to reconstruct the metabolism of the MRO of *A. ignava* and *A. flamelloides* BUSSELTON2 and found it to be a hydrogenosome. Several hydrogenosome proteins were detected in our LGT screen and are involved in several aspects of hydrogenosome metabolism such as dealing with oxidative stress and amino acid synthesis (see Table S8). The donor taxa of the hydrogenosomal LGT-derived genes are variable. These authors also found that *Anaeramoeba* spp. have several enzymes involved in ATP production that were not found in other Metamonada (Stairs et al. 2021). Some of these enzymes were acquired through LGT. For example, a bacterial 3-hydroxybutyrate dehydrogenase is found in *A. ignava* (Figure S13A), and two bacterial methylmalonyl CoA epimerases (MME) of independent origin are found in *A. flamelloides* (Figure S13B). Interestingly, *A. flamelloides* also has a eukaryotic version of the enzyme which is the one predicted to be targeted to the MRO (*A. ignava* only has a eukaryotic MME).

We used KofamKoala (Aramaki et al. 2020) to assess the extent to which *Anaeramoeba* contains metabolic “modules” (sub-pathways) encoded entirely by LGT-derived genes. We found that strongly supported LGT-derived proteins typically form chimeric pathways with eukaryotic proteins (note that less than a third of the LGT-derived genes were successfully annotated by the software). The sole exception is an apparently intact Leloir pathway in *A. flamelloides*, which suggests that the organism can metabolize galactose. Interestingly, while the pathway is typically comprised of four stand-alone enzymes, the *A. flamelloides* Leloir pathway contains two fused proteins. BUSSELTON_g20257.t1 and SCHOONER_g30019.t1 are fusions of galactose mutarotase (GalM) and galactokinase (acetyl-CoA carboxylase, GalK), and BUSSELTON_g13285.t1 and SCHOONER_g28623.t1 are fused galactokinase (GalK), galactose 1-P uridyltransferase (GalT) and UDP-galactose 4 epimerase (GalE). Phylogenetic reconstructions of each of the four enzymes shows that members of the bacterial phylum “Candidatus Omnitrophica” branch as sister taxa and thus could be the donor lineage (Figure S13C). The *Anaeramoeba* Leloir pathway genes could be derived from an operon, which would have facilitated fusion protein formation after LGT. Due to the fragmented nature of available genomic assemblies for “Candidatus Omnitrophica”, we were not able to determine whether the Leloir pathway genes indeed reside in an operon in this organism. However, such genes are operonic in other bacteria such as *E. coli* (Semsey et al. 2007) (i.e., GalETKM). Interestingly, whereas the utilization of galactose has apparently been lost in the *A. flamelloides* symbionts due to the loss of *galE*, a complete Leloir pathway is present in Sym_BMAN.

We observed differences in the predicted high-level functions of LGTs inferred to have been acquired in the common ancestor of *A. ignava* and *A. flamelloides* and those that were acquired after the two species diverged (the possibility of secondary loss must also be acknowledged). Of the genes acquired in the *Anaeramoeba* common ancestor, functional categories E (amino acid metabolism), C (Energy metabolism) and G (Carbohydrate metabolism) are the most highly represented (Figure 4C, blue bar). This is similar to what was observed in *Mastigamoeba,* a LGT-prone eukaryote found in anaerobic environments (Žárský et al. 2021), as well as in diverse parasitic eukaryotes (Alsmark et al. 2013).

We analyzed proteins predicted to have been acquired by LGT in the common ancestor of *A. ignava* and *A. flamelloides* in order to assess how they might ‘plug in’ to the metabolism of the *Desulfobacteraceae* symbionts. The results speak strongly to the complementarity of the partner organisms. For example, a cysteine synthase K and a D-3-phosphoglycerate dehydrogenase are LGT-derived (Figure S13D,E), and together with SerC phosphoserine aminotransferase (which is itself not detected in our LGT screen but found in the genomes), these enzymes are linked to glycolysis, to the re-use of SH^-^ produced by the symbionts, and to provision of acetate to the symbionts (Figure S14). This is one example where an *Anaeramoeba* pathway is composed of eukaryotic, putatively ancestral enzymes, and LGT-derived enzymes.

Another example of metabolic complementarity involves an acetate transporter (Figure 13F) acquired in the *Anaeramoeba* common ancestor. Acquisition of this enzyme by LGT might have played a role in the establishment of the symbiosis. From the data in hand it is not clear whether this LGT directly facilitates the transport of acetate from *Anaeramoeba* to the symbiont or if it simply imports acetate from the environment that could be routed to the symbiont secondarily. In any case, it is noteworthy that the acetate transporter gene has duplicated numerous times after acquisition, which is consistent with functional specialization (*A. ignava* has seven paralogs, *A. flamelloides* BUSSELTON2 has 35, and *A. flamelloides* SCHOONER1 has 32).

Genes associated with anaerobiosis were also acquired in the common ancestor of *A. ignava* and *A. flamelloides*. For example, and as has been described in several diplomonads (Jiménez-González, Xu, and Andersson 2019) and other anaerobic protists (Slamovits and Keeling 2006), *Anaeramoeba* spp. seem able to perform pyrophosphate-dependent glycolysis, which uses pyrophosphate (PPi) instead of ATP as a phosphate donor. This capacity appears to have been enabled by the *Anaeramoeba* ancestor having two pyruvate phosphate dikinase (PPD) enzymes (Figure S13G), one shared with Parabasalia and thus likely acquired before *Anaeramoeba* and *Trichomonas* diverged from one another. The other PPD is exclusive to *Anaeramoeba*. Two other enzymes acting downstream of PPD are also found in *Anaeramoeba* spp. but were not detected in our LGT screen, due to presence of eukaryote-eukaryote LGTs after an initial acquisition from bacteria. These two enzymes are ATP-independent phosphofructokinase (Figure S13H) and phosphoglycerate mutase (Figure S13I). *A. ignava* has also acquired a second copy of phosphoglycerate mutase (Figure S13J). The chimeric nature of glycolysis in *Anaeramoeba* (involving both prokaryotic and eukaryotic enzymes) may have enhanced ATP production, which would have been crucial given their lack of a TCA cycle, which relies on oxidative phosphorylation (Mertens 1993).

Genes allowing survival in anoxic environments were also acquired multiple times in the history of *Anaeramoeba*. Strategies for mitigating the loss of the eukaryotic aerobic ribonucleotide reductase (RNR) in other metamonads include reliance on the salvaging of exogenous deoxynucleosides in *Giardia* (Baum et al. 1989), acquisition of class III RNR from prokaryotes in *Spironucleus*, *Trepomonas* (Xu et al. 2016) and *Kipferlia bialata* (Jiménez-González, Xu, and Andersson 2019; Xu et al. 2020), and acquisition of class II RNR (*Trichomonas* and *Tritrichomonas*) (Lundin et al. 2010), as well as a bacterial class I RNR in *Tritrichomonas*. *A. ignava* and *A. flamelliodes* have different RNR gene suites, none of which encode the classical oxygen sensitive eukaryotic class I RNR but instead encode several class II RNRs, which are vitamin B12 dependent and oxygen insensitive (Figure S13K). *A. flamelloides* RNRs branch with the few known eukaryotic RNR class II enzymes (monomers) that are thought to have been acquired from bacteria or viruses on several occasions (Figure S13K) (Crona et al. 2013; Lundin et al. 2010), and were also putatively transferred among eukaryotes (Lundin et al. 2010). By contrast, BMAN RNRs constitute (at least) two evolutionarily distinct branches. One shares recent common ancestry with *Trichomonas vaginalis* RNR class II and archaeal RNR class II (dimer), as well as bacterial enzymes (Figure S13K), and another branches ambiguously with diverse bacteria.

The *Anaeramoeba* common ancestor appears to have acquired enzymes to deal with oxidative stress via LGT. For example, as mentioned in (Stairs et al. 2021), several NADH oxidases are found in *Anaeramoeba.* Interestingly, they were acquired on multiple occasions in *Anaeramoeba* history from different donors (see next section for a global screening of serial acquisitions) (Figure S13L), likely including *Desulfobacteraceae* (Figure S13L). Other anaerobic protists have NADH oxidases but they do not belong to the same clade, suggesting that they were acquired independently (Leger et al. 2016; Stairs et al. 2019).

*Serially acquired genes in* Anaeramoeba *speak to continual adaptive pressures*

We found evidence of serially acquired genes in *Anaeramoeba*. A total of 45 orthologs are predicted to have been acquired multiple times in one or both *Anaeramoeba* species. Additionally, 25 orthologs were independently acquired in *A. ignava* and *A. flamelloides* (or acquired in their common ancestor and then differentially lost; see Table S8). These results are summarized below.

Orthologous genes involved in fighting oxidative stress were acquired multiple times in *Anaeramoeba* history. The phylogeny of OsmC proteins (Figure S13M) suggests that they were acquired (at least) six times, four times in *A. flamelloides* and twice in *A. ignava*, from various donors. Nitroreductases were acquired once in their common ancestor, and then twice in *A. flamelloides* and once in *A. ignava* (Figure S13N). Note that the phylogeny of nitroreductases suggests that several eukaryotes, many which are anaerobes (*Trichomonas foetus* and *vaginalis*, several species of the *Entamoeba* genus, *Stygiella incarcerata*, and *Blastocystis* subtype 1) have also acquired nitroreductases from prokaryotes—and separately from *Anaeramoeba*. Thioredoxins were also acquired independently in both *Anaeramoeba* species while a rubrerythrin gene was acquired in the *Anaeramoeba* common ancestor and once in *A. flamelloides* (Figure S13O). This suggests that LGT is an ongoing feature of *Anaeramoeba*, one that gives rise to functionally differentiated genes that impact the biology of the organism. A more complete set of LGT-derived genes involved in oxygen detoxification is presented in Table S8. Several of these genes were previously described as having been laterally acquired in several Metamonada (Jiménez-González, Xu, and Andersson 2019). Strikingly, the evolutionary origins of these genes is diverse: bacteria, archaea and viruses are all predicted donors.

Biosynthesis of sterols, which are key constituents of canonical eukaryotic membranes, requires molecular oxygen. In contrast, prokaryotic hopanoid biosynthesis does not require molecular oxygen as a substrate, and the squalene is directly cyclized by the enzyme squalene-hopene cyclase (SHC) (Bloch 1965). Several anaerobic eukaryotes have been shown to encode a bacterial-like SHC, more specifically a squalene tetrahymenol cyclase (STC) that was likely transferred between eukaryotes after an initial acquisition from bacteria (Takishita et al. 2012). More recently, the anaerobic yeast *Schizosaccharomyces japonicus* was shown to have acquired a bacterial SHC independently, allowing growth in sterol-free media under anaerobic conditions (Bouwknegt et al. 2021). Strikingly, *A. flamelloides* possess two evolutionarily distinct SHC genes. The first forms a sister branch with eukaryotic STCs, while the second one emerges among bacterial SHCs (Figure 4F). Additionally, the *A. ignava* and *A. flamelloides* ancestor appears to have acquired one of the enzymes of the mevalonate pathway from archaea, i.e., hydroxymethylglutaryl-CoA reductase (one of the pathways synthesizing the terpenoid backbone). As all the other enzymes in the pathway are eukaryotic, this represents an example of orthologous replacement of the eukaryotic enzyme by its prokaryotic counterpart, resulting in a chimeric mevalonate pathway.

Two Nim genes encoding 5-nitroimidazole reductases or related pyridoxamine 5’-phosphate oxidase were independently acquired by *A. ignava* and *A. flamelloides* from various donors (Figure S13P). Nim proteins are associated with acquired resistance to metronidazole in bacterial anaerobes (Alauzet et al. 2019). Interestingly, other anaerobic protists have acquired Nim genes from bacteria (Pal et al. 2009), including *Trichomonas vaginalis*; in this case Nim proteins are targeted to the hydrogenosome (Bradic et al. 2017) (and in another recently sequenced parabasalid, *Histomonas meleagridis* KAH0787444.1). Their exact role in metroimidazole resistance is still under investigation (Pal et al. 2009) but their independent acquisitions in several anaerobic protists suggests that they are functionally important. Interestingly, while the *Desulfobacter* spp. associated with *A. ignava* has a full-length Nim gene, the Nim genes of the epibionts of *A. flamelloides* are pseudogenes.

Three genes involved in the synthesis of lysine, belonging to each of two known pathways were acquired either independently or multiple times in *Anaeramoeba* spp. (two genes in the diaminopimelic acid pathway and one in the homocitrate-aminoadipate pathway) (Figure S13Q,R). A branch comprising eukaryotic and viral ornithine decarboxylases is present in the diaminopimelate decarboxylase tree, suggesting that viruses could have mediated a eukaryote-to-eukaryote gene transfer. Both *Anaeramoeba* species also acquired several genes involved in leucine biosynthesis. They possess stand-alone LeuC and LeuD (large and small subunits of isopropyl malate isomerase, respectively), as well as a LeuC-LeuD fusion (Figure S13T,U). These genes were seemingly acquired separately, as the trees for each subunit show that the stand-alone and fused versions do not cluster together (Figure S13T).  *A. ignava* fused LeuC-LeuD has eight introns while the standalone subunits have none, suggesting that the former was acquired earlier than the latter. In line with this hypothesis, the standalone subunits share 66.6 % identity with their best prokaryotic hit on average (Table S8) (the fused versions still have on average 57.8 % of identity with their best prokaryotic hits). Interestingly, the fused and unfused versions of the LeuC and LeuD genes in *A. ignava* BMAN are next to each other in the genome despite their independent origin (in the following order: LeuD; LeuC; LeuD-LeuC fusion). This suggests that rearrangements happened after acquisition.

Finally, the phylogeny of two FAD-dependent dehydrogenases (Figure S13V) belonging to the same COG underscores the extent to which LGT has impacted *Anaeramoeba* genomes. One LGT-derived gene is present in both species and emerges among archaeal sequences. *A. flamelloides* encodes a second FAD-dependent dehydrogenase. Its closest sister branch is an enzyme from a eukaryotic dsDNA virus and bacterial sequences. The topology of the latter clade is also compatible with a virus being an intermediate between prokaryotic species and *Anaeramoeba*.

*Enrichment of ribosomal proteins suggest specialized compartmentalization in A. flamelloides*

Ribosomes are formed by rRNA and ribosomal proteins (RPs). In eukaryotes, they consist of a small subunit (40S) formed by 18S rRNA and up to 33 RPs, and of a large (60S) subunit formed by 28S, 5.8S, and 5S rRNAs and up to 46 RPs (Ban et al. 2014). We retrieved two 18S rRNA sequences for BUSSELTON2 located in different scaffolds, one is a fragment of 1041bp and the other one is 3654bp long sharing 99% identity, and also two sequences for SCHOONER1 of 3362 bp and 3446 bp long with shared identity of 99.3%. Five and seven 28S rRNA sequences with varied lengths (i.e., 1384 to 4562 bp, with identities ranging from 90-99.8%) were identified in different SCHOONER1 and BUSSELTON2 scaffolds, respectively. We were able to validate most of the annotated RPs from 40S and 60S subunits that were flagged by our expansion analysis (Figure S12, Table S7J,K). In addition, our findings showcase *Anaeramoeba*’s RPs repertoire as the most complete among all metamonads studied, and that expansions in *A. flamelloides* correspond to a recent event that occurred in the last common ancestor of SCHOONER1 and BUSSELTON2 (Data not shown). Proteins within each family are very similar in size and percentage identity at the amino acid and nucleotide levels. Given these high identities, we queried 10 proteins located up and downstream of each RP ortholog to find out whether they were similar or shared similar functional annotations with other scaffolds along loci containing their respective RP ortholog. We found that RPs-flanking proteins belong to loci associated with varied functions that were rarely shared among the RPs-flanking regions of their respective orthologs in all comparisons. This reinforces that those expansions are bona fide and not assembly duplication artifacts. Since RPs were expanded, we looked for evidence of expansion in tRNAs with tRNA-scan-SE (Chan and Lowe 2019). This yielded 667, 682 and 111 tRNA copies in BUSSELTON2, SCHOONER1, and BMAN, respectively. The copy numbers for BUSSELTON2 and SCHOONER1 appear to be consistent with the expansion of gene families associated to translation metabolic pathways.

The near complete 40S and 60S RPs repertoire, their large abundances, and the expansion in tRNA genes may suggest that *A. flamelloides* strains are able to assemble ‘subpopulations’ of ribosomes, and given the intricate relationships that have probably been developed between the host and symbionts, it may be likely that these subpopulations of ribosomes can have unique properties that inﬂuence the functions of the proteins they produce, in a similar way as shown by new evidence from model organisms (reviewed in (Filipovska and Rackham 2013)). Hence, since ribosomal proteins provide access to unique modes of translation, it might be possible that *A. flamelloides* has developed some sort of ‘ribosome code’ as ribosomes assembled with varied ribosomal protein composition may confer specialized functions, and ribosomal proteins paralog specificity may define novel means of translational control (Filipovska and Rackham 2013; Segev and Gerst 2018; Ghulam, Catala, and Abou Elela 2020).

*Extended analyses of vitamin B12 in* Anaeramoeba *and* Desulfobacter *symbionts*

Vitamin B12 (vitB12), also known as cobalamin, is a complex Co^2+^-containing modified tetrapyrrole that acts as a cofactor. In prokaryotes, there are over 15 enzymes that have a vitB12 cofactor (Shelton et al. 2019). The ability to synthesize cobamides *de novo* involves approximately 30 steps (Warren et al. 2002) and is found in ~37% of prokaryotes (Shelton et al. 2019). No known eukaryote presents evidence for cobamide *de novo* biosynthesis. Nonetheless, some encode a few enzymes that modify cobalamin or use it as cofactor (Orłowska, Steczkiewicz, and Muszewska 2021). Only four of the latter are known (Orłowska, Steczkiewicz, and Muszewska 2021; Crona et al. 2013).  First, methylmalonyl-CoA mutase (MCM) and methylmalonyl-CoA epimerase (MCE) are involved in odd-chain fatty acid metabolism in the mitochondria of animals. Second, B12-dependent ribonucleotide reductase (RNR) is found in a few mostly anaerobic eukaryotes. There are both B12-independent and B12-dependent forms of ribonucleotide reductase (RNR) (type I & III RNR are B12 independent, type II is B12 dependent), involved in deoxyribose biosynthesis (Hamilton 1974; Carell and Seeger 1980). Most eukaryotes have the type I isoform, but this isoform is oxygen sensitive, so several anaerobes have acquired vitB12-dependent RNR from bacteria, viruses, or even other eukaryotes on several occasions (Crona et al. 2013; Helliwell et al. 2011). Third, two isozymes of methionine synthase are similarly found. MetH is B12-dependent and is found in animals, while an alternative B12-independent form of methionine synthase (*metE*) is found in land plants and fungi. Based on the presence of *metH* and the absence of functional *metE*, approximately half of all cultured eukaryotic algal species from marine and freshwater environments are predicted to require exogenously produced corrinoids for growth (Croft et al. 2005; Helliwell et al. 2011). This property appeared not to be monophyletic but rather scattered in the tree. MetH has a higher catalytic rate compared to MetE (Gonzalez et al. 1992), factors that might explain why it can be advantageous to possess *metH* despite its dependency to vitB12. Several studies suggest that vitB12 might be exchanged between prokaryotic producers and B12 auxotroph algae, which would provide the bacteria with photosynthate (Croft et al. 2005; Grant et al. 2014). Other studies predict that corrinoid requirements can be fulfilled through indirect production and release into the water column upon death and cell lysis (Droop 2007).

The initial observations that (i) *Anaeramoeba* have acquired from bacteria vitB12-dependent metH and vitB12-dependent RNR (and that the vitB12-independent isozymes are absent) (ii) the *Desulfobacter* symbionts encode some cobalamin biosynthesis genes, led us to wonder if *Anaeramoeba* have additional enzymes requiring vitB12 and if vitB12 could be exchanged between *Anaeramoeba* and *Desulfobacter*.

In order to better understand the vitB12 picture of the *Anaeramoeba* system, we:

(A) scanned the *Desulfobacter* symbionts associated with *A. ignava* and *A. flamelloides* genomes for vitB12 synthesis and vitB12 uptake genes to confirm that they can synthesize it *de novo*

(B) scanned the *Anaeramoeba* genomes for

(i) additional vitB12-dependent enzymes, besides MetH and RNR.

(ii) cobalamin synthesis genes. Indeed the acquisition from bacteria of enzymes involved in the final steps of cobalamin synthesis was described in some diatoms (Vancaester et al. 2020).

(iii) vitB12 import enzymes.

(A) The *Desulfobacter* symbionts associated with *A. ignava* and *A. flamelloides* (BUSSELTON2 and SCHOONER1) seems to encode a full pathway for vitB12 synthesis including: several cobalt import proteins, ALA synthesis, synthesis of the tetrapyrrole precursor, corrin ring synthesis, adenosylation, nucleotide loop assembly, aminopropanol linker, synthesis of lower ligand DMB, addition of alpha-ribazol phosphate. The only enzyme that is not found is CobC which is responsible for the very last step in which the phosphate is removed from the lower ligand alpha-ribazol after the later was added to the corrin ring. None of CobZ (Zayas, Woodson, and Escalante-Semerena 2006), CblX, CblY and CblZ, (Rodionov et al. 2003),  which are nonorthologous replacements of CobC were found either. However, this step might not be needed for cobalamin use in *Desulfobacter* or this step might occur through an undescribed phosphatase as (i) *Desulfobacterium autotrophicum*, another member of *Desulfobacteraceae* was shown experimentally to produce cobalamin but not to encode CobC either (Shelton et al. 2019); (ii) in some archaea (e.g., *Halobacterium*), the phosphatase has not been identified (Escalante-Semerena 2007) suggesting that other non orthologous phosphatase are yet to be described.

Despite likely producing cobalamin *de novo*, the *Desulfobacter* symbionts also have several enzymes involved in the uptake of cobalamin related to BtuB. The coexistence of the *de novo* synthesis pathway and uptake proteins is known to occur in other vitB12 producers as well (Shelton et al. 2019). The *Desulfobacter* symbionts have several vitB12-dependent enzymes so do not produce cobalamin only for *Anaeramoeba*.

(B)

(i) *A. ignava* and *A.* *flamelloides* have orthologs of the three vitB12-dependent enzymes found in eukaryotes. As previously mentioned, the type II RNR and metH were acquired from bacteria, while methylmalonyl-CoA mutase and methylmalonyl-CoA mutase-associated GTPase MeaB (MeaB is a small G-protein involved in loading coenzyme B_12_ to MCM (Padovani and Banerjee 2009) are eukaryotic.

Interestingly, *A. flamelloides* strains BUSSELTON2 and SCHOONER1 appear to possess two additional LGT-derived vitB12-dependent enzymes, which were never described to occur in eukaryotes:

A B12-dependent reductive dehalogenase opens the possibility that *A. flamelloides* might be able to metabolize halogenated compounds. The *A. flamelloides* enzymes is related to archaeal proteins. The presence of this dehalogenase in members of the *Lokiarchaeota* and *Thorarchaeota* was recently highlighted (Manoharan et al. 2019; Spang et al. 2019). The *Desulfobacter* symbionts do not encode the othologous gene but have the related vitB12-dependent epoxyqueusine reductase.

 The B12-dependent Ethanol ammonia-lyase, EutBC. Ethanolamine serves as a source of carbon and nitrogen for a variety of bacteria (Garsin 2010; Lundgren et al. 2016; Tsoy, Ravcheev, and Mushegian 2009). EutBC is the core enzyme in ethanolamine utilisation. It breaks down ethanolamine into acetaldehyde and ammonia (Tsoy, Ravcheev, and Mushegian 2009).

Crucially, the *Desulfobacter* symbionts associated with *A. flamelloides* appear not to have EutBC while the symbiont associated with *A. ignava* encodes it. A similar pattern is found for EutT, which is the corrinoid-adenosyltransferase dedicated to produce the vitB12 cofactor ethanolamine for ethanolamine ammonia lyase, and for the acetaldehyde dehydrogenase EutE. After the break down of ethanolamine by EutBC, EutE converts the acetaldehyde intermediate to acetyl-CoA, which enters the carbon pool of the cell. This suggests that the latter do rely less than the former on *Anaeramoeba* to feed the Wood-Ljungdahl pathway with acetate.

This observation is in line with a longer association between *A. flamelloides* and its symbiont compared with the *A. ignava*/*Desulfobacter* system.

The ethanolamine utilisation genes in bacteria are organised in operons that are variable in length, ranging from the core EutB and EutC only in some species to up to 17 genes in other (Tsoy, Ravcheev, and Mushegian 2009). Interestingly, *Anaeramoeba* Ethanolamine ammonia lyase consist of a fused EutB and EutC as found in some *Deltaproteobacteria* (Tsoy, Ravcheev, and Mushegian 2009) while the *A. ignava* symbiont has open reading frame for EutB and one for EutC.

Nor *A. flamelloides* BUSSELTON2 or *A. flamelloides* SCHOONER1 have EutE, suggesting that the Ethanolamine ammonia lyase is not devoted to produce acetate. Rather, ethanolamine might be used as a source of nitrogen. Interestingly, *A. flamelloides* have the alcohol dehydrogenase EutG that converts acetaldehyde into ethanol (also found in the three symbionts). In *Anaeramoeba*, the enzyme appears to have been acquired from archaea.

(ii)

- The eukaryotic cobalamin adenosyl transferase is found in the three Anaeramoebae. This enzyme catalyzes the conversion of cobalamin into adenosylcobalamin, the form used as a cofactor by Methylmalonyl-CoA mutase.

- *A. ignava* have laterally acquired an alpha-ribazol phosphatase (CobC), corresponding to the very last step of cobalamin synthesis (that was not found in the symbionts). The lateral acquisition of some steps of cobalamin synthesis including cobC has occurred in a subset of diatoms auxotroph for vitB12 (Vancaester et al. 2020). We have discussed the fact that the absence of cobC in the *Desulfobacter* might not be crucial given that cobC is not found in the symbionts. Consequently, and because *A. flamelloides* is lacking cobC, it is unclear what cobC is doing in *A. ignava*. One possibility is that the cobalamin is imported in *A. ignava* with the phosphate attached to the lower ligand and that the removal of the phosphate happens in the eukaryote.

(iii)

- none of the described prokaryotic cobalamin transporters described in (Degnan et al. 2014; Rempel et al. 2018; Rodionov et al. 2003; Zhang et al. 2009) were found in *Anaeramoeba*.

- nor was the diatom-like cobalamin binding protein CBA1 (Bertrand et al. 2012).

However, (i) not much is known about cobalamin import in eukaryotes outside of metazoan (Quadros, Nakayama, and Sequeira 2005; Zhang et al. 2009) (ii) the extreme proximity between the symbiont and *Anaeramoeba* likely favorize the import of vitB12 produced by the symbiont by *Anaeramoeba*.

The mechanism by which corrinoids are released from corrinoid-producing microbes into the environment is unclear. As yet, no active means of corrinoid export has been identified (Seth and Taga 2014).

### Naming of the symbionts

Since we have not been able to grow these organisms in pure culture, we do not fulfill the requirements of the *Bacteriological Code* (1990 revision) for a valid species description. Following the guidelines developed by SeqCode (Hedlund et al. 2022) we describe these taxa using the genome sequence as the underlying type. We describe two species that are hereafter referred to “*Desulfobacter amicus*” as the symbiont of *A. ignava* BMAN, and “*Desulfobacter proximus*” as the symbiont in *A. flamelloides* BUSSELTON2/SCHOONER1.

**Description of candidate taxa**

“*Desulfobacter* *amicus*” ([*Desulfobacteraceae*, δ-Proteobacteria] NC; NA; R; NAS [GenBank number CP054837], oligonucleotide sequence complementary to unique region of 16S rRNA is 5’- GACTTATGCCAACCGACTTTATCC-3’ [probe BMAN], S [*Anaeramoeba ignava* BMAN {Anaeramoebidae, Metamonada, Eukaryota}, vacuole]; Anaer.; M). Etymology: amicus, m [Latin] adj. supporting, propitious, helpful. Refers to the close association with its host. Additional information: Numerous cells clustered in vacuoles, tightly associated with the mitochondrion-related organelles of the host cell *Anaeramoeba ignava* (Anaeramoebidae, Metamonada, Eukaryota); phylogenetically defined to contain the sequences of strain BMAN (CP054837) but not those of *Desulfobacter proximus* (CP054838) and (JABUQY000000000); Reference strain: BMAN (inhabiting *Anaeramoeba ignava* strain BMAN). Locality of reference strain: Balikpapan, Kalimantan, Indonesia. 1°20′S, 116°50′E. Reference material: Permanent protargol preparations of the strain BMAN, deposited in the collection of the National Museum in Prague, Czech Republic, inventory numbers P6E 4142 – P6E 4148.

“*Desulfobacter proximus*” ([*Desulfobacteraceae*, δ-Proteobacteria] NC; NA; R; NAS [GenBank numbers CP054838 and JABUQY000000000], oligonucleotide sequence complementary to unique region of 16S rRNA is 5’- CTTATGCTGATCAACTTTATCCGGT-3’ [probe BUSS/SCH], S [*Anaerameoba flamelloides* BUSSELTON2 {Anaeramoebidae, Metamonada, Eukaryota}, vacuole]; Anaer.; M). Etymology: proximus, m [Latin] adj. nearest, next, neighbor. Refers to the close association with its host. Additional information: Numerous cells clustered in vacuoles, tightly associated with the mitochondrion-related organelles of the host cell *Anaeramoeba flamelloides* (Anaeramoebidae, Metamonada, Eukaryota); phylogenetically defined to contain the sequences of strain BUSSELTON2 (CP054838) and SCHOONER1 (JABUQY000000000) but not those of *Desulfobacter* *amicus* (CP054837); can be discriminated from *Desulfobacter* *amicus* and other *Desulfobacterales* by fluorescence in situ hybridization with the probe BUSS/SCH. Reference strain: BUSSELTON2 (inhabiting *Anaeramoeba flamelloides* strain BUSSELTON2). Locality of reference strain: Busselton, Western Australia, Australia. 33°38′S, 115°20′E. Reference material: Permanent protargol preparations of the strain BUSSELTON22, deposited in the collection of the National Museum in Prague, Czech Republic, inventory numbers P6E 4137 – P6E 4141.
