## Supplementary material for "A unique symbiosome in an anaerobic single-celled eukaryote": Table S1-S3: Sequencing data and metrics, Symbiont genome characteristics, Introns statistics

**Table S1** General genome statistics for *A. ignava BMAN, A. flamelloides* BUSSELTON2 *and A. flamelloides* SCHOONER1*.*

|  | *Anaeramoeba ignava* BMAN | *Anaeramoeba flamelloides* BUSSELTON2 | *Anaeramoeba flamelloides* SCHOONER1 |
| --- | --- | --- | --- |
| Genome size (Mbp) | 42,843,398 | 254,083,75 | 259,028,67 |
| # of scaffolds | 147 | 76 | 346 |
| N50 | 512,274 | 12,797,701 | 1,575,887 |
| GC% | 19.35 | 23.63 | 24.77 |
| # of genes | 14,837 | 30,401 | 30,317 |
| # of CDSs | 14,759 | 29,744 | 29,839 |
| # of tRNAs | 74 | 637 | 456 |

**Table S2** Symbiont genome characteristics.

|  | Sym_BMAN | Sym_BUSS2 | Sym_SCH1 | *Desulfobacterium autotrophicum* |
| --- | --- | --- | --- | --- |
| Chromosome size (bp) | 6,055,721 (closed circular) | 4,968,664 (closed circular) | 4,994,604  (26 contigs) | 5,657,780 |
| CDS (functional) | 5,256 | 3,840 | 3,823 | 4,768 |
| GC% | 52.96 | 47.60 | 47.50 | 48.80 |
| Pseudogenes | 37 | 922 | 1,000 | 79 |
| IS elements^1^ | 60 | 624 | 666 | 75 |
| Symbionts/amoeba | 12.6 | 35.3 | 36.5 | - |

^1^Predicted by ISsaga v2.0 (http://issaga.biotoul.fr/ISsaga2/).

**Table S3A** Introns

|  | *Anaeramoeba ignava* BMAN | *Anaeramoeba flamelloides* BUSSELTON2 | *Anaeramoeba flamelloides* SCHOONER1 |
| --- | --- | --- | --- |
| # of introns | 32,108 | 39,831 | 39,247 |
| Average length (bps) | 300 | 1,459 | 1,504 |
| Median length (bps) | 102 | 264 | 278 |
| Mode length (bps) | 76 | 124 | 113 |
| Longest (bps) | 4,168 | 10,402 | 10,385 |
| Shortest (bps) | 35 | 49 | 44 |
| Percentage of genome | 22.5 | 22.8 | 22.7 |
| Introns/genomic kb | 0.74 | 0.15 | 0.15 |
| Introns/gene | 2.13 | 1.3 | 1.3 |
| GC% of all introns (genome) | 8.14 (19.36) | 22.8 (23.63) | 22.06 (23.78) |

**Table S3B** Noncanonical intron boundaries

|  | *Anaeramoeba ignava* BMAN | *Anaeramoeba flamelloides* BUSSELTON2 | *Anaeramoeba flamelloides* SCHOONER1 |
| --- | --- | --- | --- |
| AT-AC | 33 | 0 | 0 |
| GC-AG | 20 | 7 | 6 |
| GT-AA | 1 | 0 | 0 |
| TA-AG | 1 | 0 | 0 |
| 5’ (G/A)TATCCTTT | 32 (27 A, 5 G) | 8 (8 G) | 10 (10 G) |
