## Supplementary material for "A unique symbiosome in an anaerobic single-celled eukaryote": Figure S1-17

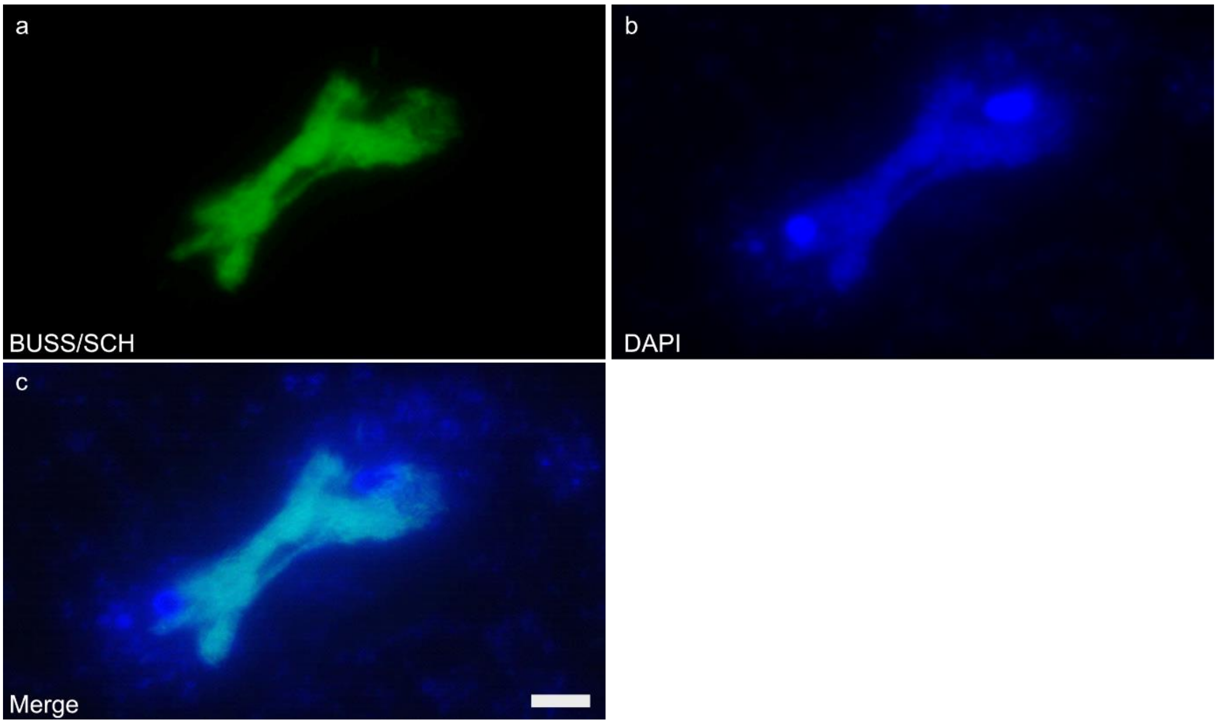


**Figure S1: *A. flamelloides* BUSSELTON2 symbionts segregating during cytokinesis.** *A. flamelloides* BUSSELTON2 hybridized with **a,** probe BUSS/SCH-BMN-488 and stained with **b,** DAPI. **c,** merged image of **a,** and **b,**. Scale bar 5 µm.

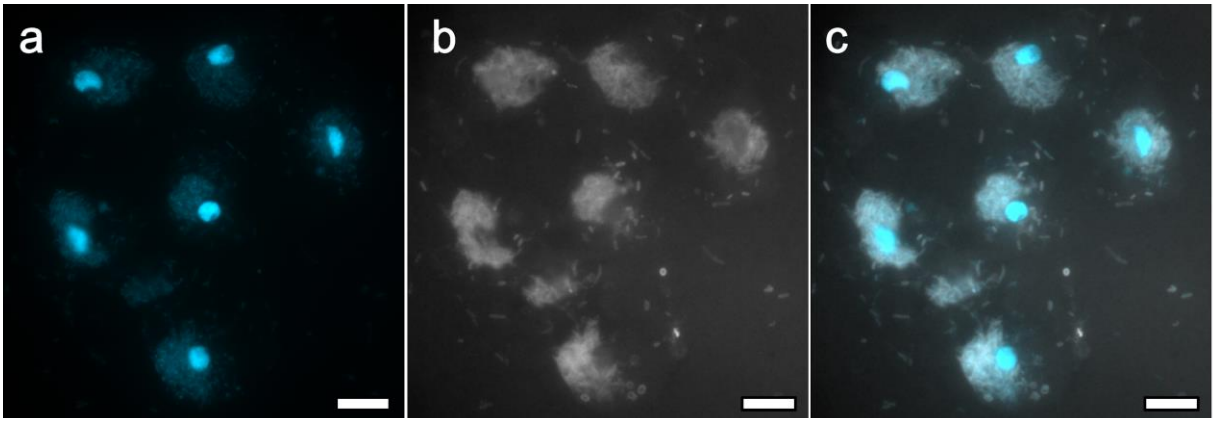


**Figure S2: Wheat germ agglutinin (WGA) staining of *Anaeramoeba* and symbionts.** *A. flamelloides* BUSSELTON2 cells were live-stained for 10 min using WGA-CF633, washed using artificial sea water and fixed in 4% formaldehyde. The cells were mounted in DAPI containing mounting media and were imaged using wide-field microscopy. The surface of the symbiont is stained by the WGA lectin at a similar intensity as free-living bacteria. **a,** *A. flamelloides* BUSSELTON2 stained with DAPI. **b,** WGA-CF633 staining, **c,** merged image of **a,** and **b,**. Scale bar 10 µm.

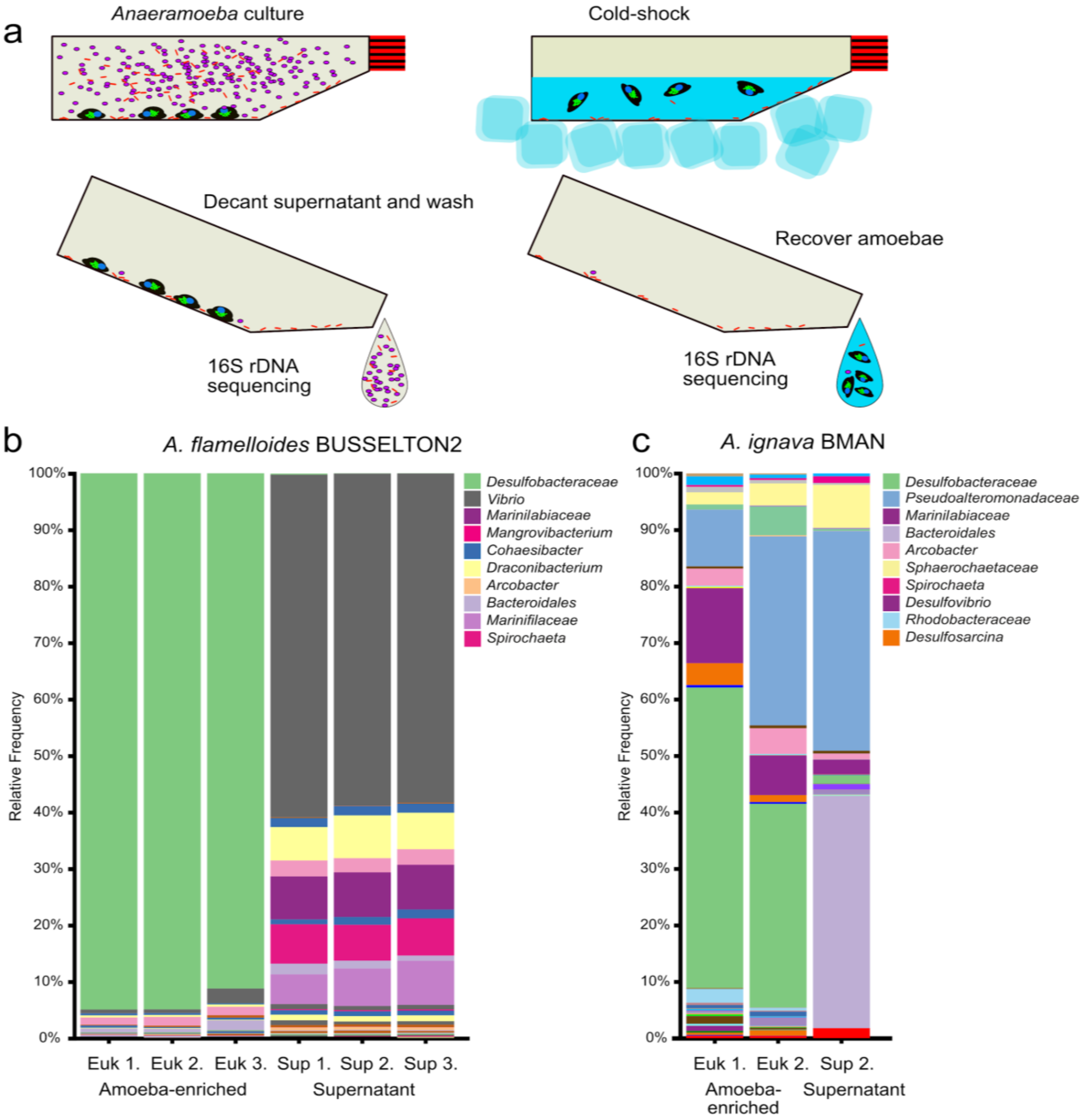


**Figure S3: Amoebae enrichment procedure and 16S rDNA sequencing. a,** The amoebae are enriched from xenic cultures by resuspending planktonic bacteria and decanting the supernatant followed by several washes using artificial sea water. The amoebae are selectively enriched from adherent bacteria using cold-shock detachment and differential centrifugation. The efficiency of enrichment is monitored by 16S rDNA sequencing. **b,** The relative frequency of bacterial taxa based on the V4 region of 16S rDNA analysis from triplicate amoeba-enriched and supernatant fractions of *A. flamelloides* BUSSELTON2. **c,** The relative frequency of bacterial taxa based on analysis of the V4 region of 16S rDNA from two amoeba-enriched and one supernatant fraction of *A. ignava* BMAN. The enrichments in **b,** and **c,** were prepared using the method described in **a,**.

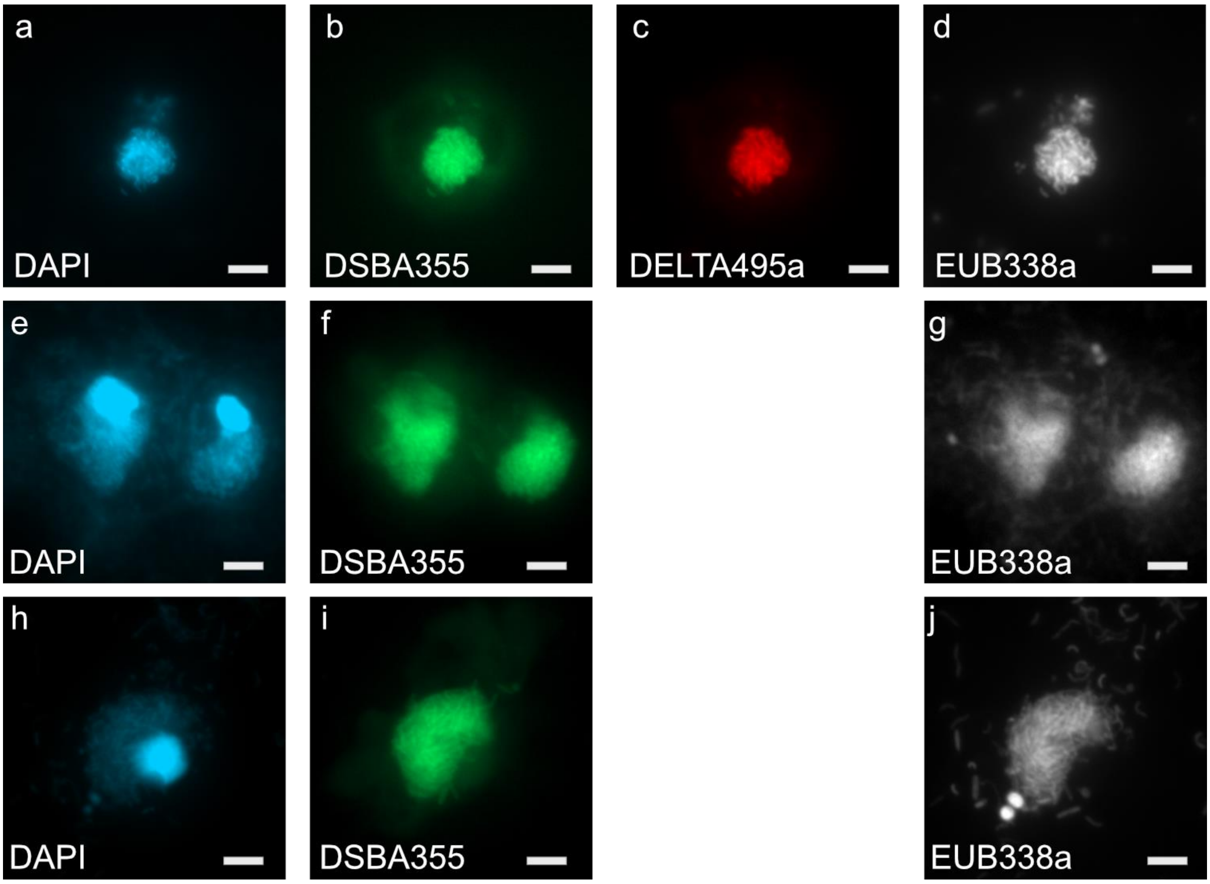


**Figure S4: Fluorescence in situ hybridization (FISH) of the symbionts in *A. ignava* BMAN and *A. flamelloides* BUSSELTON2/SCHOONER1. a-d,** *A. ignava* BMAN stained with **a,** DAPI and hybridized with **b,** probe DSBA355-BMN-488, **c,** probe Delta495a-Atto 550 and **d,** probe EUB338a-Atto 633. **e-g,** *A. flamelloides* BUSSELTON2 stained with **e,** DAPI and hybridized with **f,** probe DSBA355-BMN-488, and **g,** probe EUB338a-Atto 633. **h-j,** *A. flamelloides* SCHOONER1 stained with **h,** DAPI and hybridized with **i,** probe DSBA355-BMN-488, and **j,** probe EUB338a-Atto 633. Scale bar 1 µm.

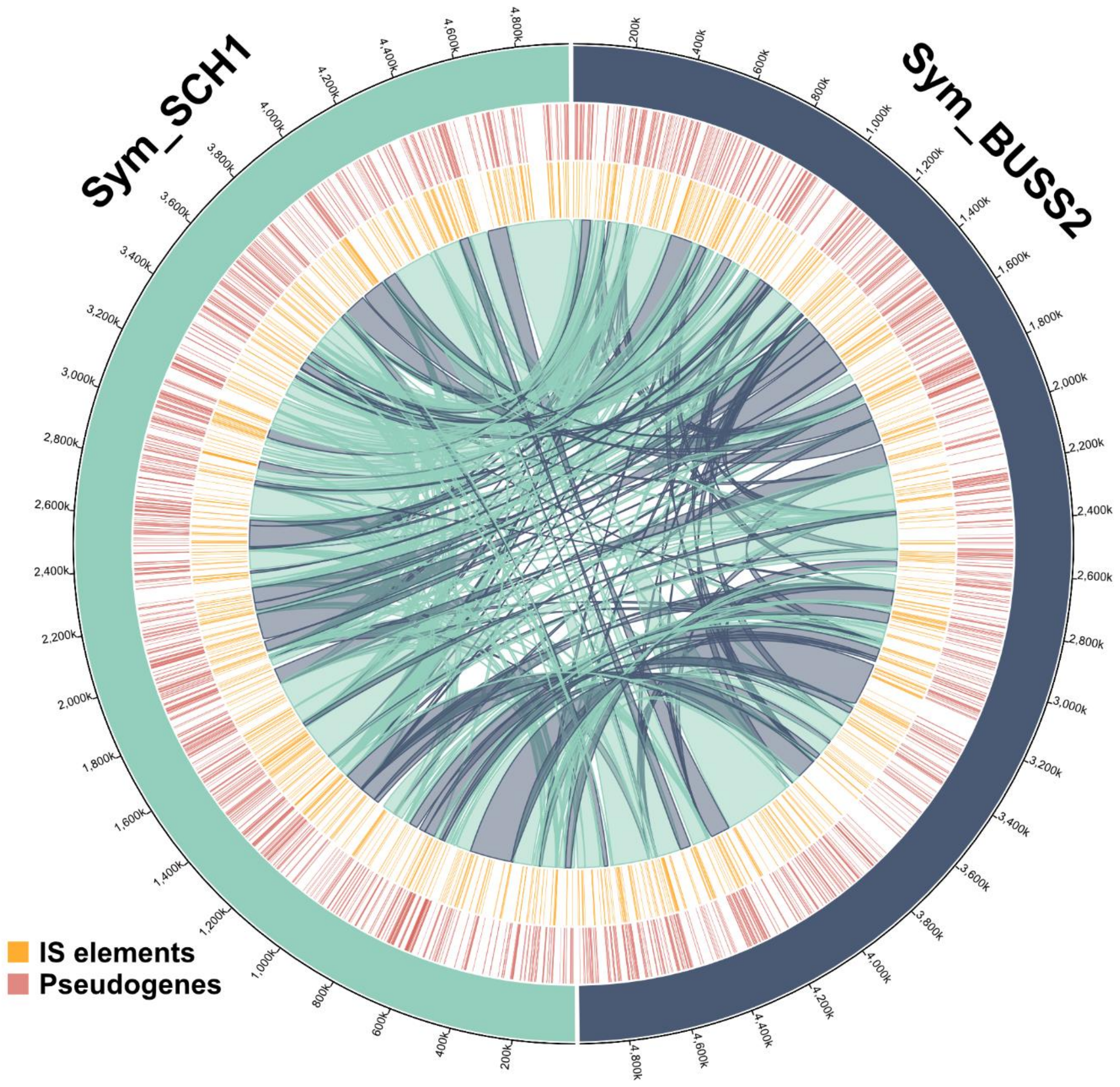


**Figure S5 Extensive synteny differences between Sym_BUSS2 and Sym_SCH1.** The synteny of the Sym_BUSS2 genome (4,958,664 bp) and the largest contig of the Sym_SCH1 genome (4,969,409 bp) were determined using Sibelia (Minkin et al. 2013) and visualized as a ribbon-plot (dark blue – sense segment, cyan – inverted segment). The inner track (yellow lines) shows IS elements predicted using ISsaga 2.0 (http://issaga.biotoul.fr/). The middle track (red lines) shows the positions of pseudogenes. The outer track displays the respective genomes (cyan – Sym_SCH1, dark blue – Sym_BUSS2). Genome coordinates are indicated by outer ticks every 200 kbp. The figure was prepared using Circa (OMGenomics).

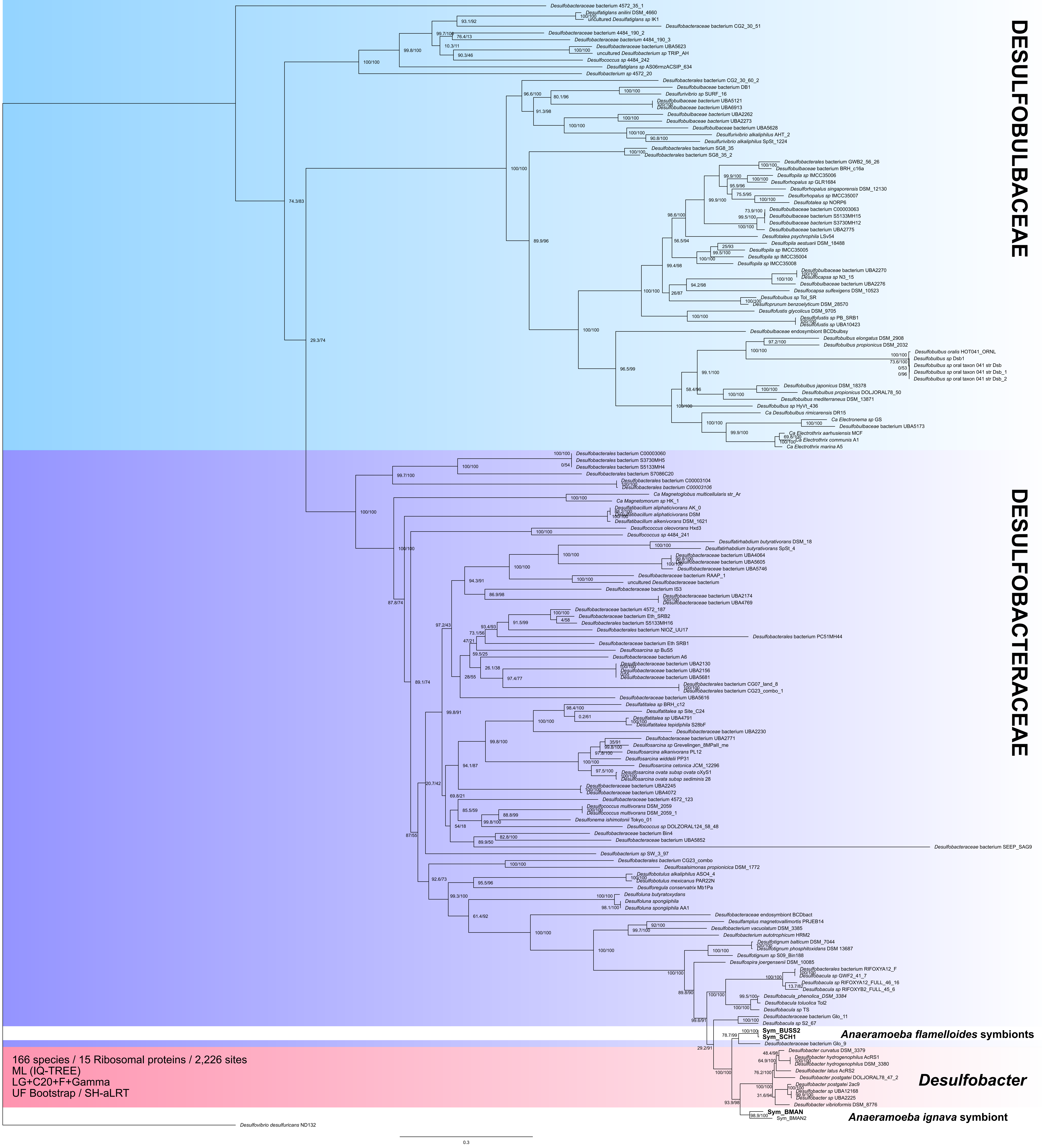
DESULFOBULBACEAE
DESULFOBACTERACEAE
Anaeramoeba flamelloides symbionts
Sym_BUSS2
Sym_SCH1
Desulfobacter
Sym_BMAN
Anaeramoeba ignava symbiont
166 species / 15 Ribosomal proteins / 2,226 sites
ML (IQ-TREE)
LG+C20+F+Gamma
UF Bootstrap / SH-aLRT
Desulfovibrio desulfuricans ND132
0.3

**Figure S6: The *Anaeramoeba* symbionts belong to the *Desulfobacteraceae* family.** Phylogenomic analysis is inferred from 168 taxa, 15 ribosomal proteins, and 2,226 sites. The ML tree was estimated with IQTree under the LG+C20+F+Gamma model of evolution. Bipartition support values are derived from 1,000 ultrafast bootstrap / SH-aLRT bootstraps. Scale bar indicates inferred number of substitutions per site. Tree files and alignments are available at FigShare: https://doi.org/10.6084/m9.figshare.20375601

**Figure S7A:** The distribution of IS elements in positional bins of 25 kb along the chromosome of Sym_BUSS2.

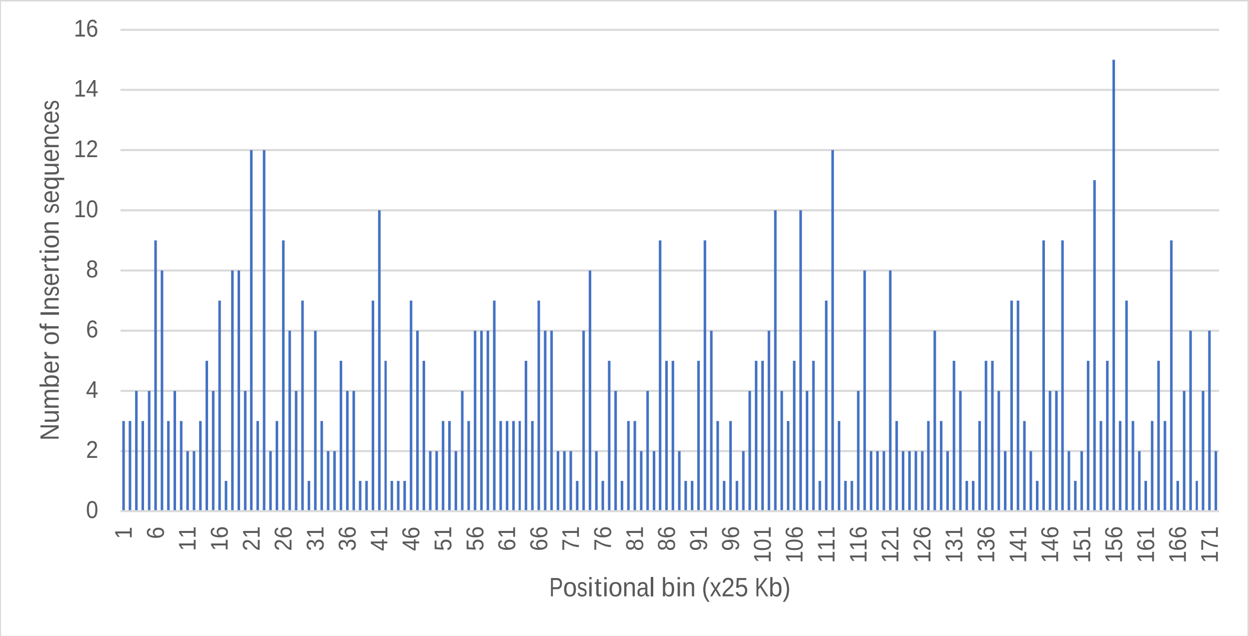


**Figure S7B:** The distribution of IS elements in positional bins of 25 kb along the chromosome of Sym_SCH1.

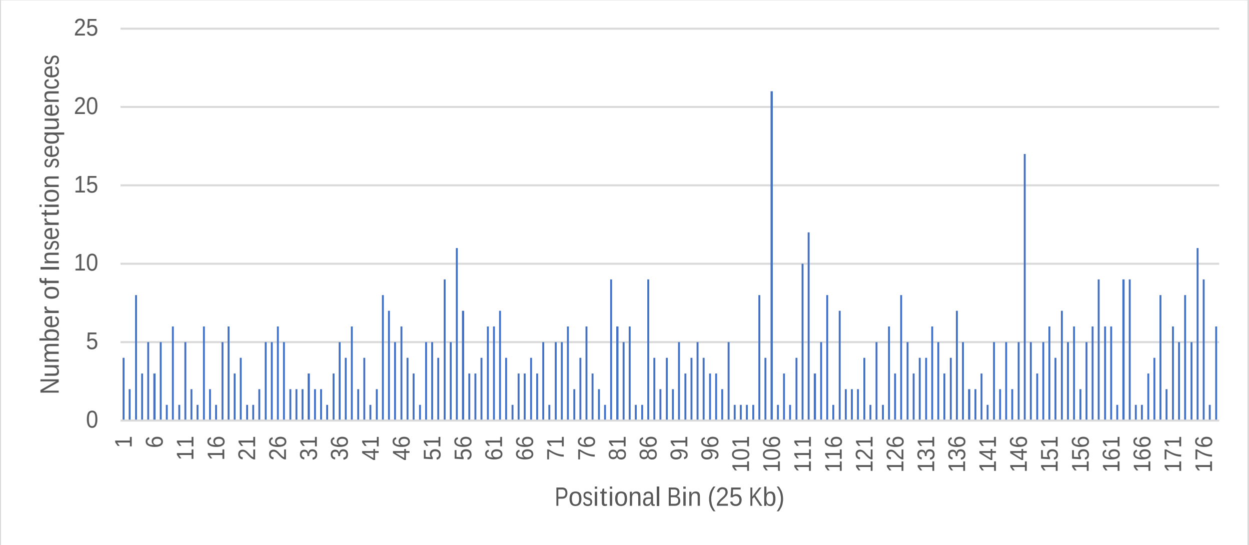

**Figure S7C:** Distance bins (500 bp) of IS elements positions of Sym_BUSS2 to a shift in synteny between Sym_BUSS2 and Sym_SCH1. The synteny blocks were inferred using Sibelia. The five most common IS elements categories are shown.

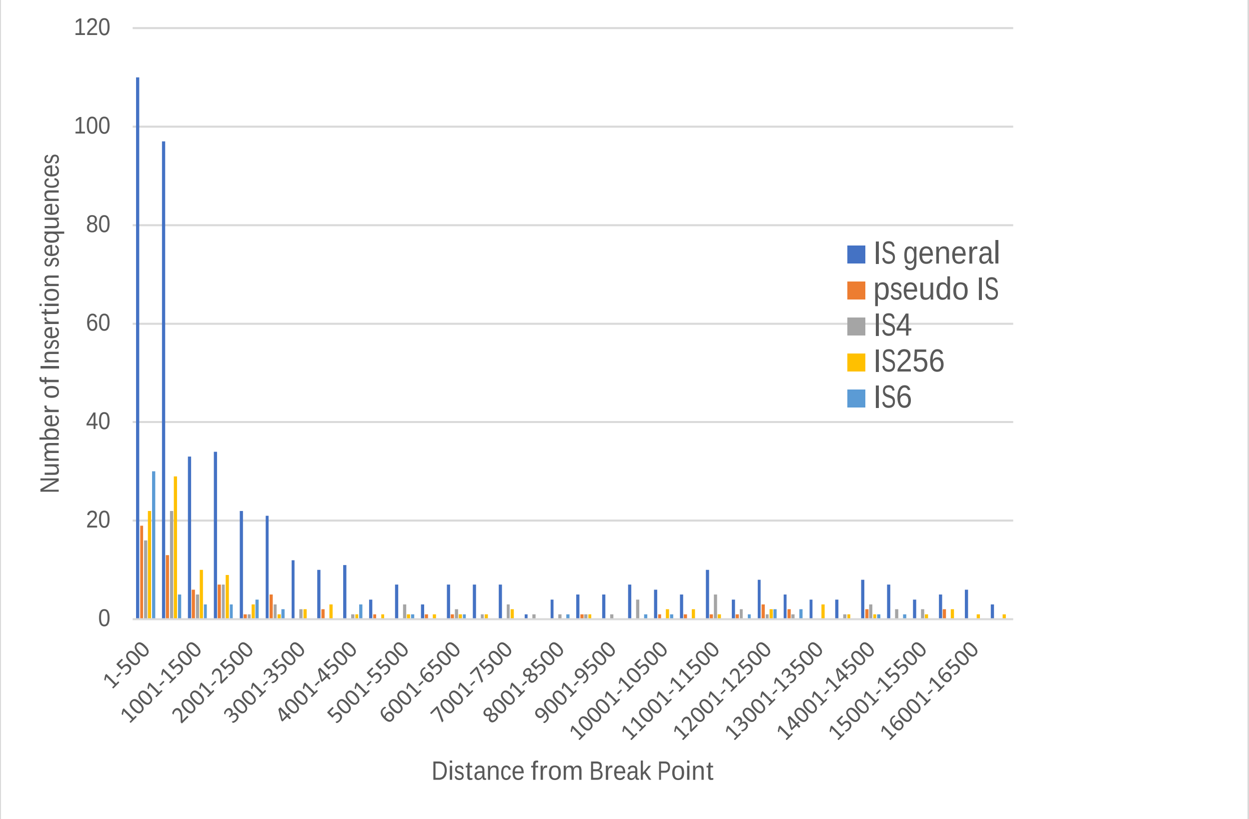


**Figure S7D:** Distance bins (500 bp) of IS elements positions of Sym_SCH1 to a shift in synteny between Sym_BUSS2 and Sym_SCH1. The synteny blocks were inferred using Sibelia. The five most common IS elements categories are shown.

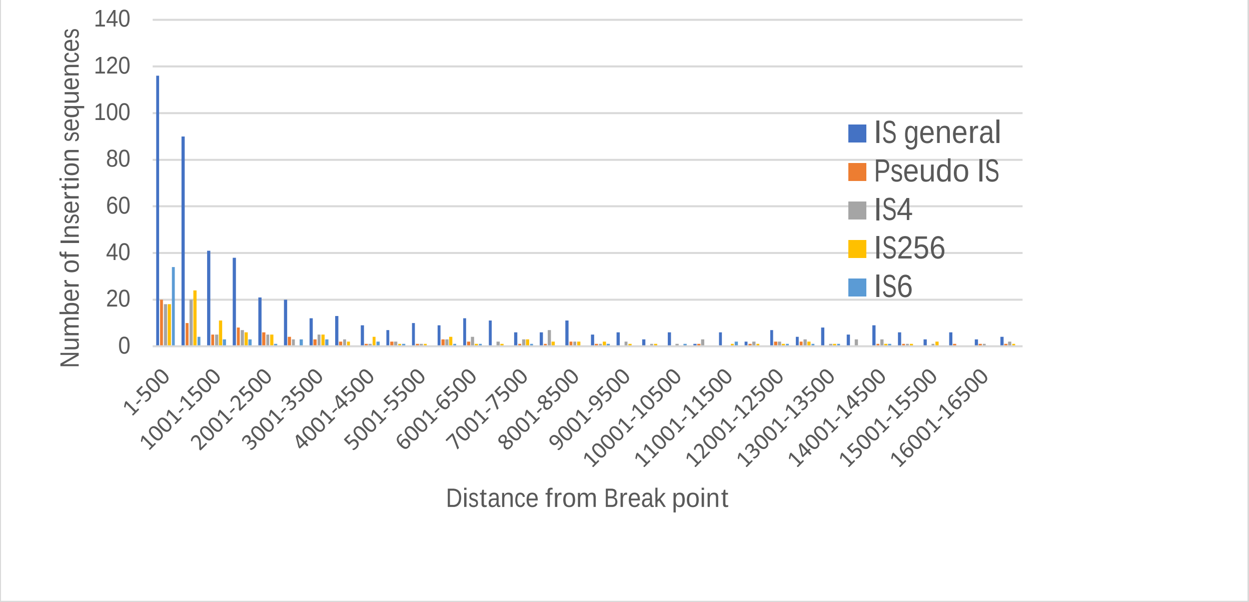

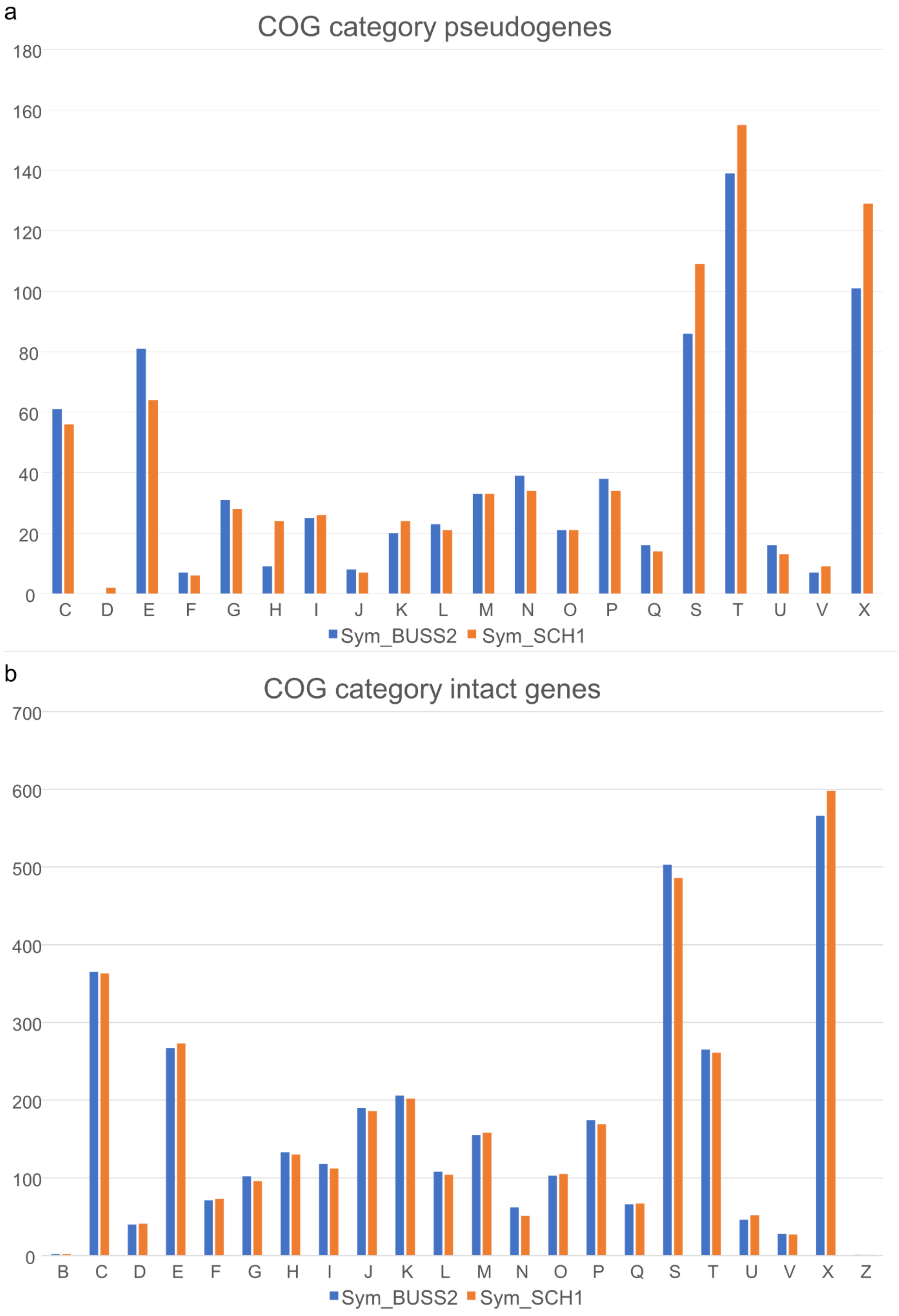
a
COG category pseudogenes
180
160
140
120
100
80
60
40
20
0
C D E F G H I J K L M N O P Q S T U V X
Sym_BUSS2 Sym_SCH1
b
COG category intact genes
700
600
500
400
300
200
100
0
B C D E F G H I J K L M N O P Q S T U V X Z
Sym_BUSS2 Sym_SCH1

**Figure S8: Genes in signal transduction are commonly pseudogenized *A. flamelloides* symbionts.** COG categories of *A. flamelloides* symbiont pseudogenes **a**, and intact genes **b,.** The COG categories of pseudogenes and intact genes for Sym_BUSS2 (blue) and Sym_SCH1 (orange) were determined by eggnog-mapper v2.1.7. Proteins classified in multiple COG categories were counted once in each category. The y-axis shows the number of classifications. Abbreviations: B - Chromatin structure and dynamics, C - Energy production and conversion, D - Cell cycle control and mitosis, E - Amino acid metabolism and transport, F - Nucleotide metabolism and transport, G - Carbohydrate metabolism and transport, H - Coenzyme metabolism, I - Lipid metabolism, J - Translation, K - Transcription, L - Replication and repair, M - Cell wall/membrane/envelope biogenesis, N - Cell motility, O - Post-translational modification, protein turnover, chaperone functions, P - Inorganic ion transport and metabolism, Q - Secondary Structure, T - Signal transduction, U - Intracellular trafficking and secretion, Y - Nuclear structure, Z - Cytoskeleton, R - General functional prediction only, S - Function unknown, V - Defense mechanisms, X - Mobilome: prophages, transposons.

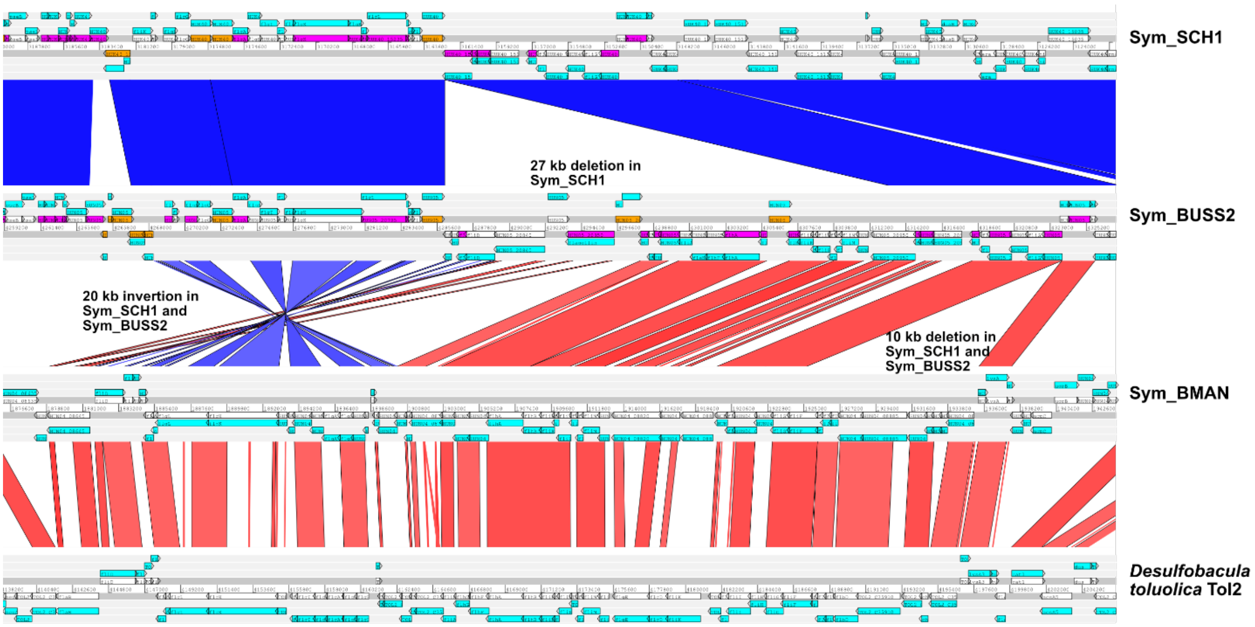


**Figure S9: Pseudogenization of the flagellar operon in Sym_SCH1 and Sym_BUSS2.** A synteny plot of the flagellar operons of Sym_SCH1, Sym_BUSS2, Sym_BMAN and the free-living *Desulfobacula toluolica* Tol2. Sym_BMAN and *Desulfobacula toluolica* Tol2 share conserved gene order and lack pseudogenes (purple) and IS elements (orange). Sym_SCH1 and Sym_BUSS2 have experienced substantial genome decay in this region. This is due to deletions, inversions, frame-shift pseudogenization as well as IS element induced pseudogenization. The part of the genome upstream on the flagellar operon in Sym_SCH1 shows no evidence of decay. Syntenic regions as determined by blastn similarity. Select IS element connections were pruned to IS to increase viewability.

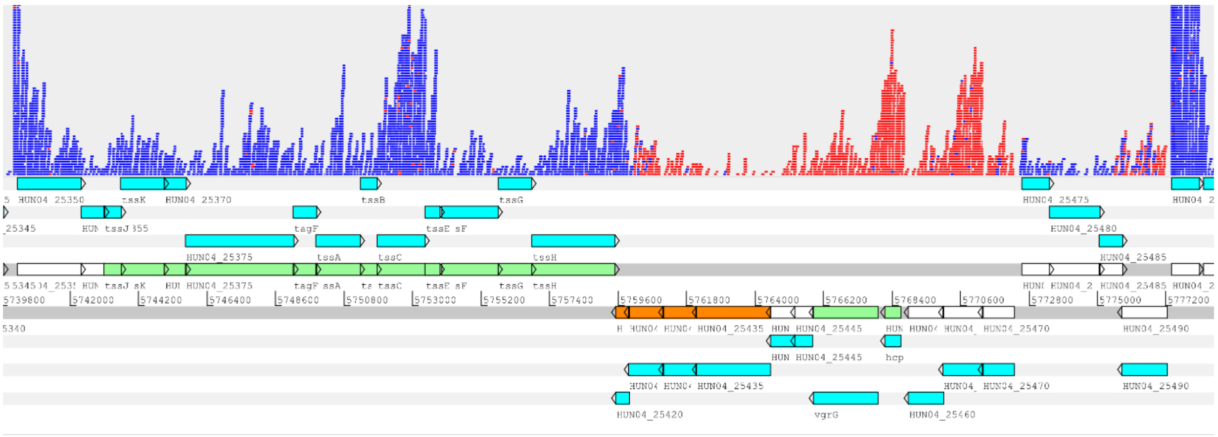


**Figure S10: Type VI secretion system in Sym_BMAN.** Sym_BMAN encodes a type VI secretion system (T6SS) that consists of two convergent transcription units of 25 genes, with most genes showing evidence of expression. The first transcription unit includes 12 genes (*tssJKLM*, *tagF*, *tssA*, *tssBC*, *tssEFGH*) and two hypothetical proteins. The second transcription unit encodes, *vgrG*, *hcp*, and upstream of these; a forkhead associated domain (FHA) protein, a TPR repeat protein, a carbohydrate esterase 4 (CE4) superfamily protein. The downstream genes include a DUF4280 family protein previously suggested as potential spike (PAAR) proteins (Lays, Tannier, and Henry 2016). Directly downstream of *vgrG*, we identified a DUF4123 protein (HUN04_25440), which is generally encoded upstream of putative T6SS effectors (Liang et al. 2015). The four effector candidates (HUN04_25435, HUN04_25430, HUN04_25425, HUN04_25420) encoded between the DUF4123 domain protein and DUF4280 domain protein have no known function, but putative homologs are present in both free-living and bacterial symbiont genomes. Light green – conserved T6SS genes. Orange – effector candidate genes.

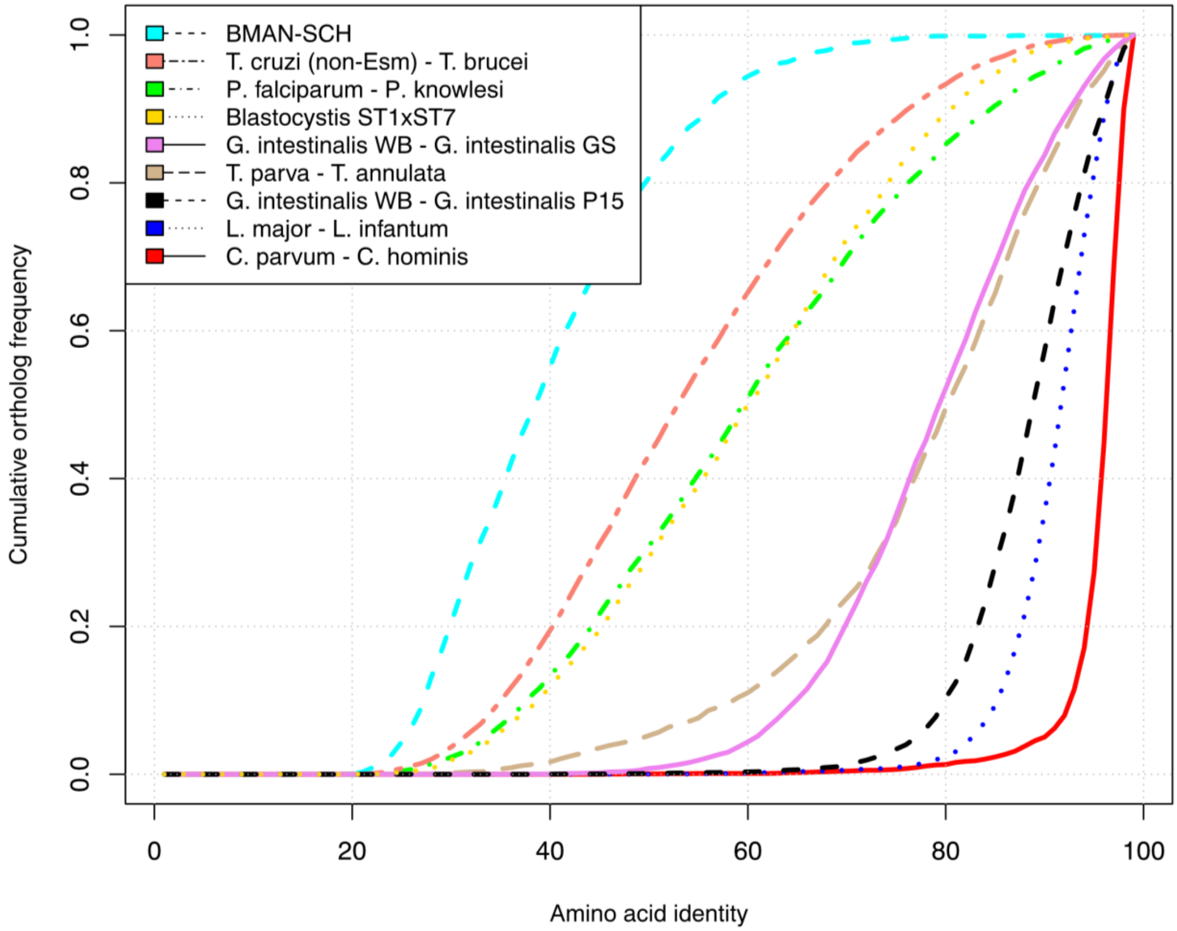


**Figure S11: A comparison of amino acid identities between *Anaeramoeba* species to other eukaryotes.** Cumulative ortholog frequency is plotted on the y-axis while the x-axis represents the amino acid identity between orthologs. The more similar species pairs are to each other, the steeper the curve and further to the right. Note that the order of the pairs in the legend matches the order in the figure. Comparisons for *Anaeramoeba* species were based on aligned regions of reciprocal best Basic Local Alignment Search Tool (BLAST) hits, while non-*Anaeramoeba* data were obtained from the authors of a comparative study of *Giardia intestinalis* (Jerlstrom-Hultqvist et al. 2010) and *Blastocystis* ST1 (Gentekaki et al. 2017).

a

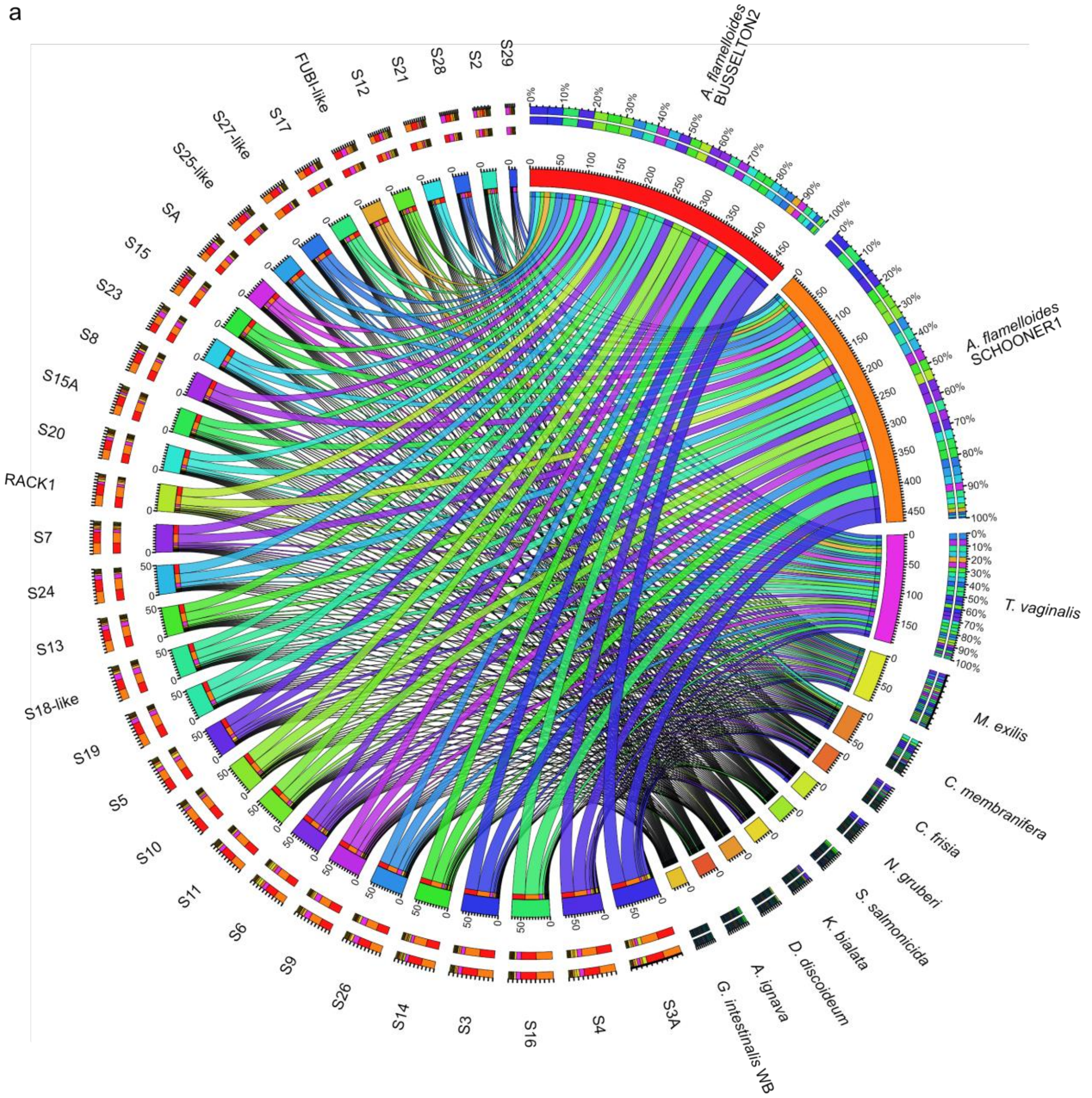
A. flamelloides BUSSELTON2
A. flamelloides SCHOONER1
T. vaginalis
M. exilis
C. membranifera
C. frisia
N. gruberi
S. salmonicida
K. bialata
D. discoideum
A. ignava
G. intestinalis WB
S3A
S4
S16
S3
S14
S26
S9
S6
S11
S10
S5
S19
S18-like
S13
S24
S7
RACK1
S20
S15A
S8
S23
S15
SA
S25-like
S27-like
S17
FUBI-like
S12
S21
S28
S2
S29

b

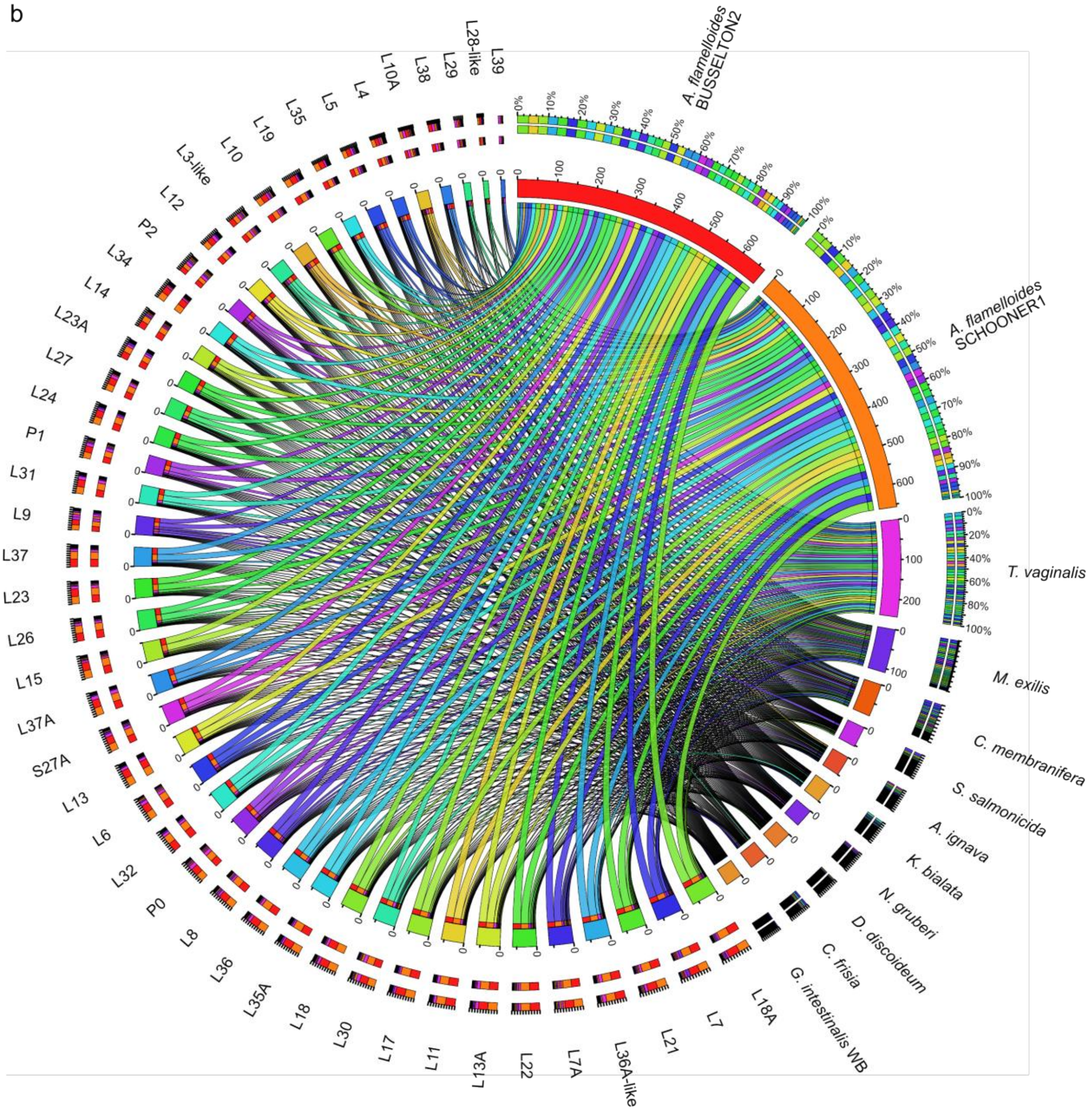


**Figure S12: Expanded copy number of ribosomal proteins (RP) in *A. flamelloides*.** The *A. flamelloides* genome is highly amplified in **a,** 40S ribosomal subunits, and **b,** 60S ribosomal subunits. Color-coded taxa [BUSSELTON2 – *G. intestinalis* WB] are depicted clockwise and sorted from the largest to the smallest number of RP orthologs detected in each taxon [S3A-S29 or L18A-L39]. Each color-coded ribbon from each taxon represents the RP ortholog specified in the opposite end of the ribbon and the width of the ribbon indicates the size of the expansion. The percentage of the expansion within each taxon matches the color-coded ribbon and it is depicted in the outer circle next to each taxon label. The outer circle next to the RP orthologs is color-coded by taxon. For example, in figure **a,** BUSSELTON2 largest expansions are for the orthologs S3A and S4 with 26 and 23 proteins, respectively (an approximation of these counts is shown in the inner red circle under the taxon name that is marked from 0 to 450 with small divisions each indicating 5 proteins). The expansions of these two orthologs amount to 10% of all the considered RP orthologs in BUSSELTON2. In terms of S3A and S4, the outer circle indicates that most proteins for such orthologues belong to BUSSELTON2 (red) and SCHOONER1 (orange).

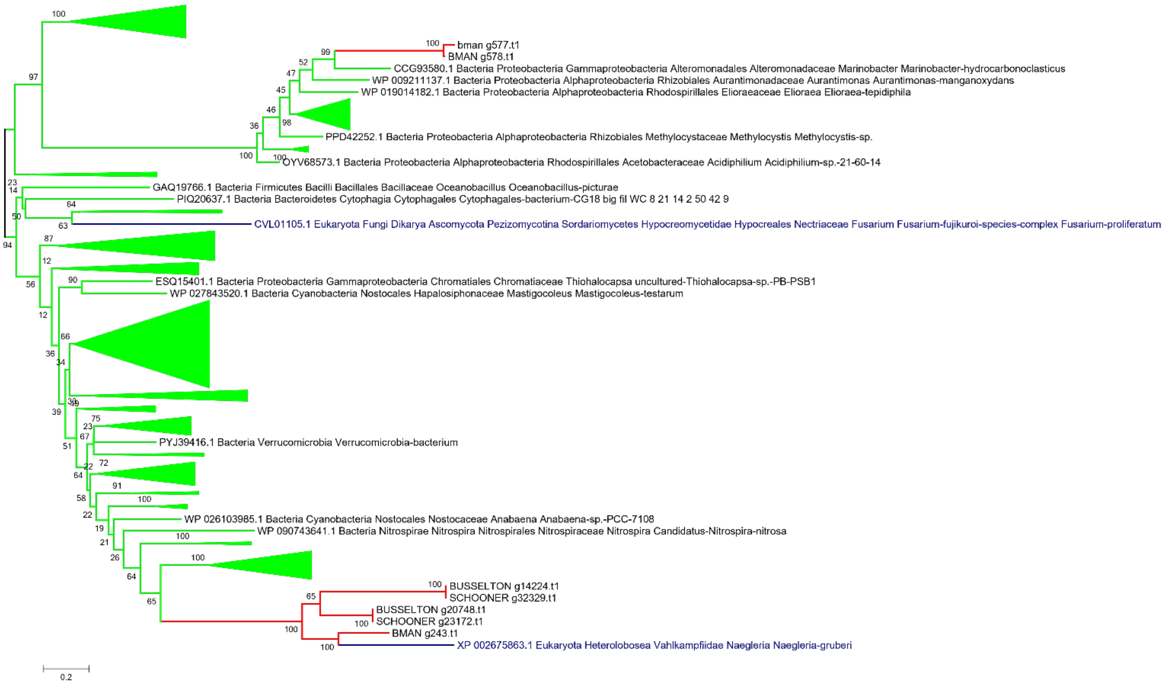


**Figure S13A** Phylogenetic reconstruction of 3-hydroxybutyrate dehydrogenase. Alignment with MAFFT, site selection BMGE and phylogeny IQtree model C20+G4 1000 ultrafast bootstraps. *Anaeramoeba* sequences are in red, eukaryotic sequences in blue, and prokaryotic sequences in bright green.

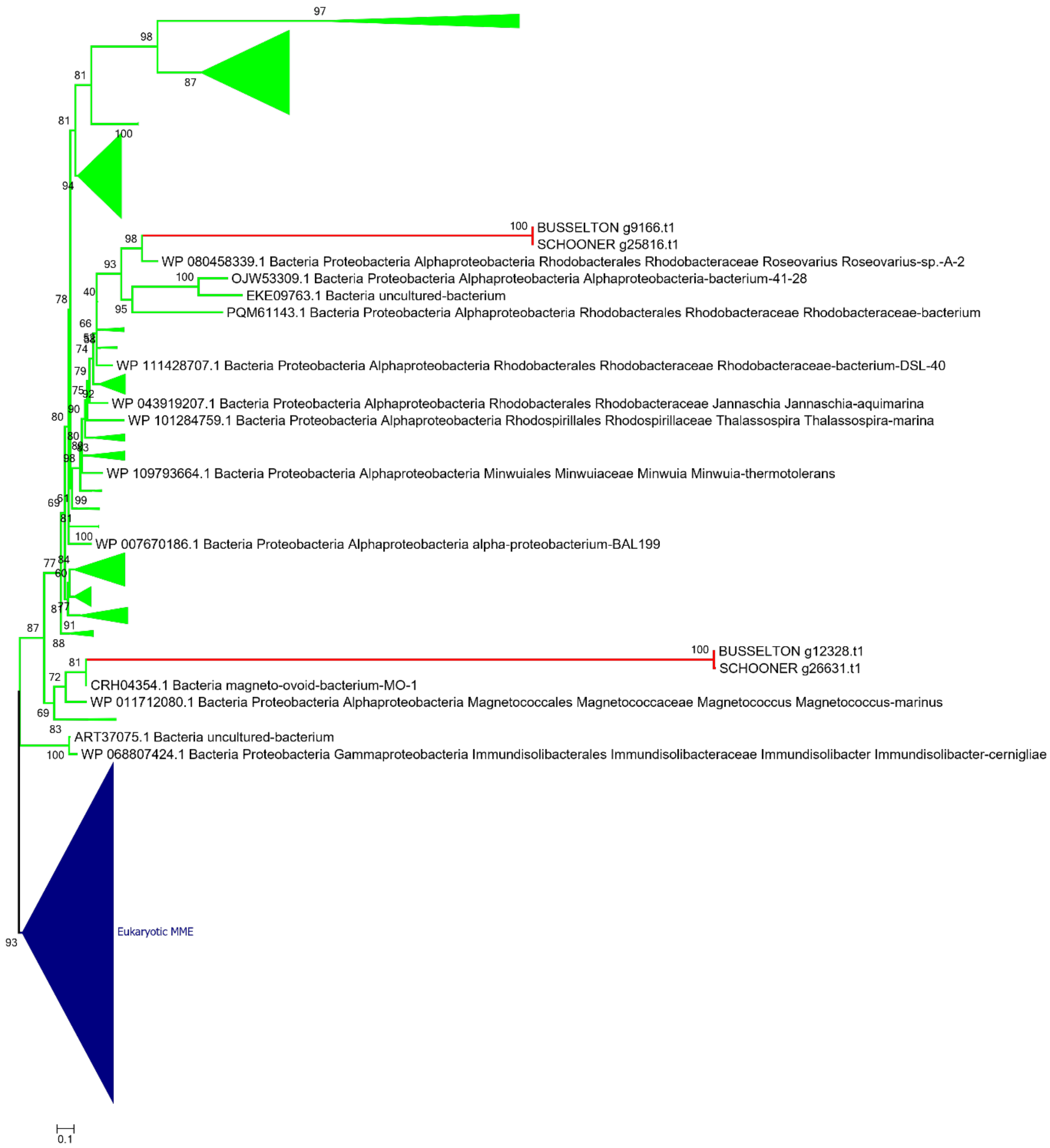


**Figure S13B** Phylogenetic reconstruction of Methylmalonyl CoA epimerase (MME). Alignment with MAFFT, site selection BMGE and phylogeny IQtree model C20 1000 ultrafast bootstrap. *Anaeramoeba* sequences are in red, eukaryotic sequences in blue, and prokaryotic sequences in bright green.

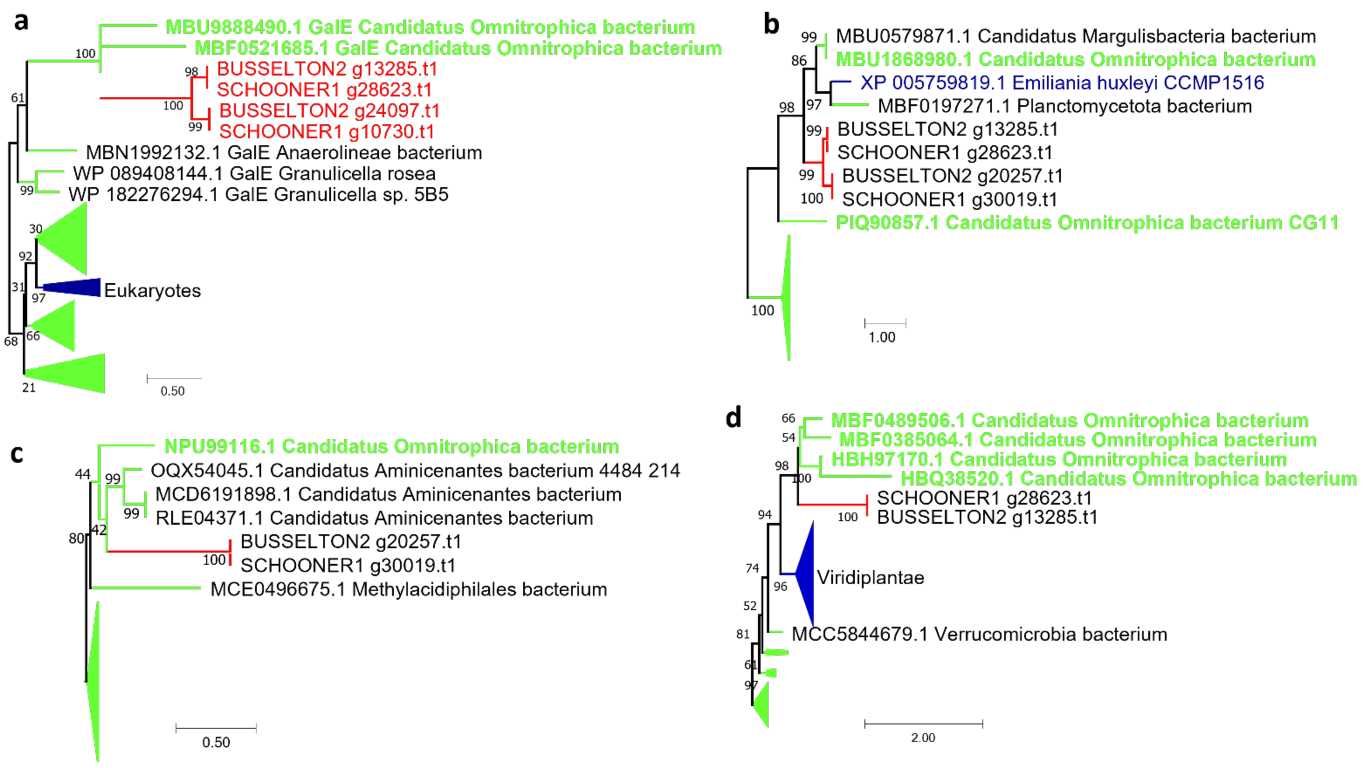


**Figure S13C** Phylogenetic reconstruction of the 4 enzymes constituting the Leloir pathway. Alignment with MAFFT, site selection BMGE, and phylogeny IQtree model C60 with 1000 ultrafast bootstrap. *Anaeramoeba* sequences are in red, eukaryotic sequences in blue, and prokaryotic sequences in bright green. Candidatus omnitrophica bacterium labels are in green bold characters. **a**, UDP-galactose 4 epimerase (GalE). **b**, galactokinase (GalK). **c**, galactose mutarotase (GalM). **d**, galactose 1-P uridyltransferase (GalT).

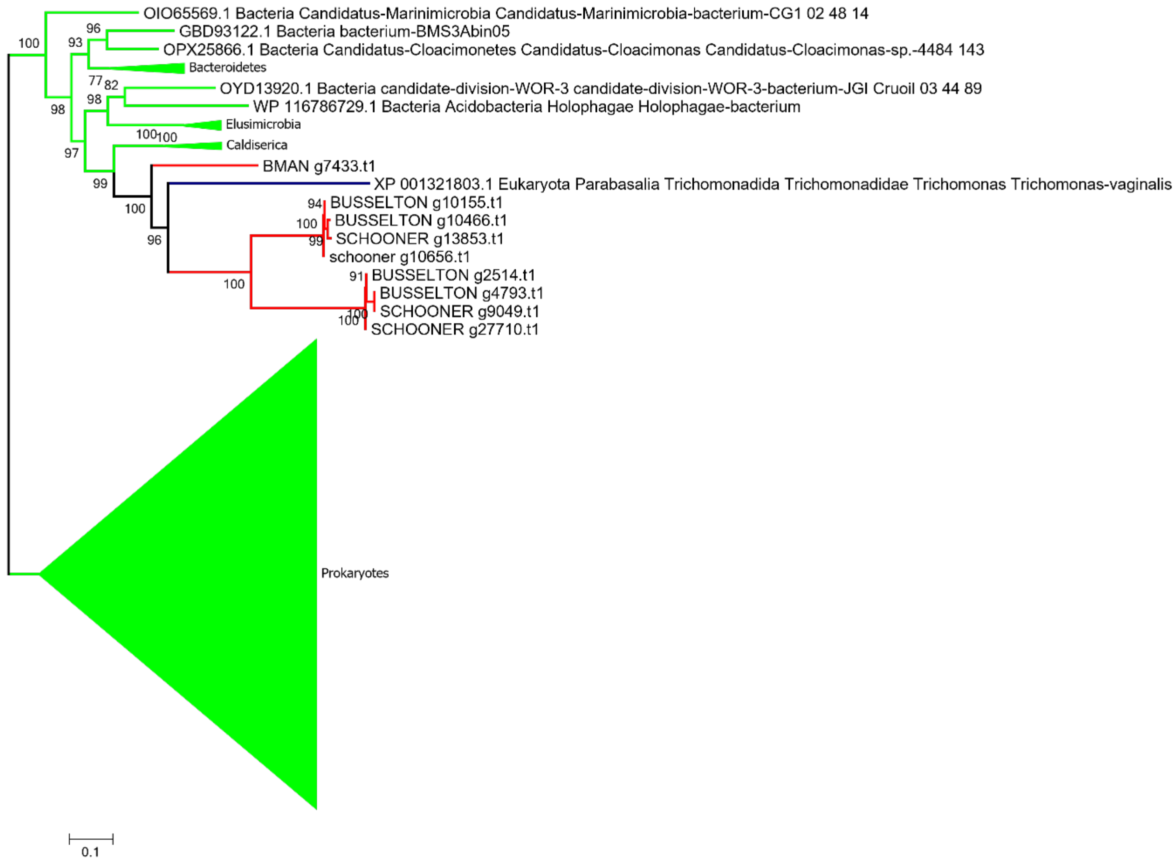


**Figure S13D** Phylogenetic reconstruction of CysK. Alignment with MAFFT, site selection BMGE and phylogeny IQtree model LG4X 1000 ultrafast bootstrap. *Anaeramoeba* sequences are in red, eukaryotic sequences in blue, and prokaryotic sequences in bright green.

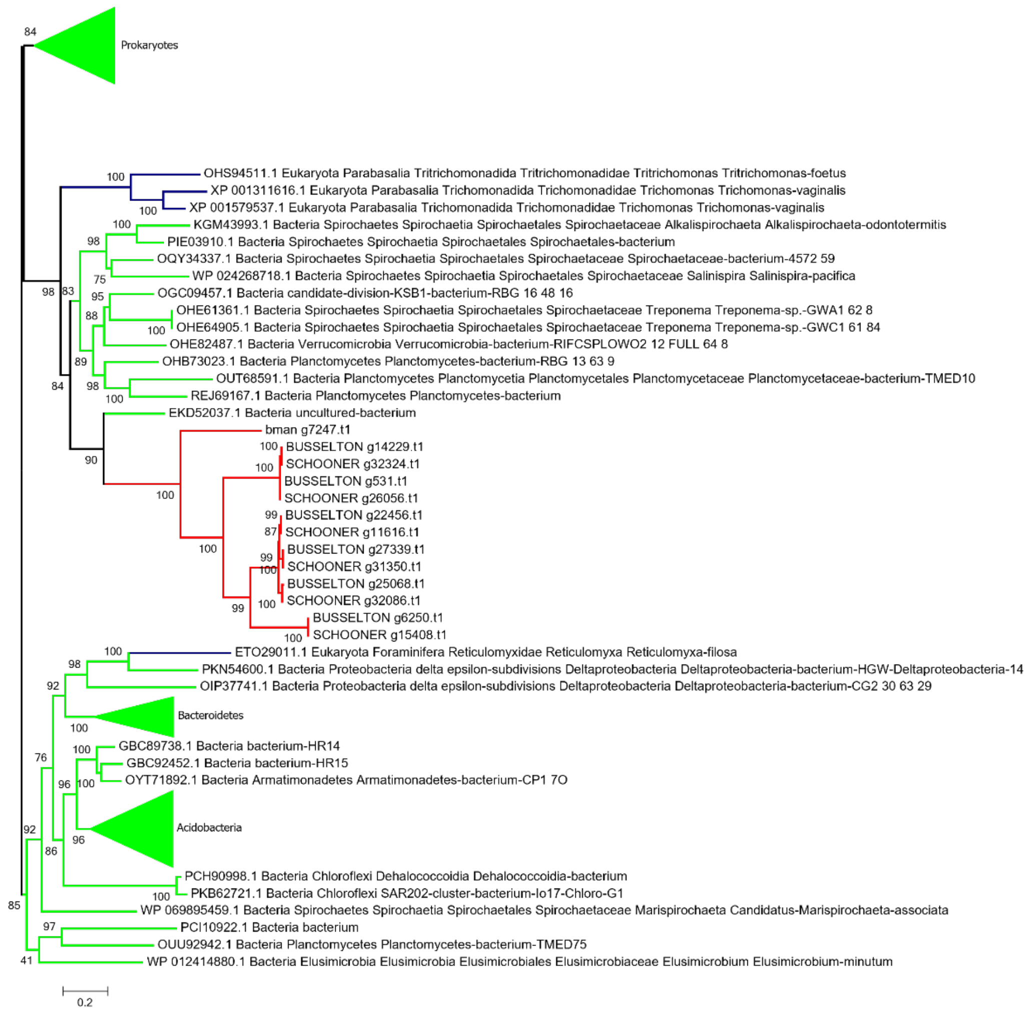


**Figure S13E** Phylogenetic reconstruction of D-phosphoglycerate dehydrogenase. Alignment with MAFFT, site selection BMGE and phylogeny IQtree model LG4X 1000 ultrafast bootstrap. *Anaeramoeba* sequences are in red, eukaryotic sequences in blue, and prokaryotic sequences in bright green.

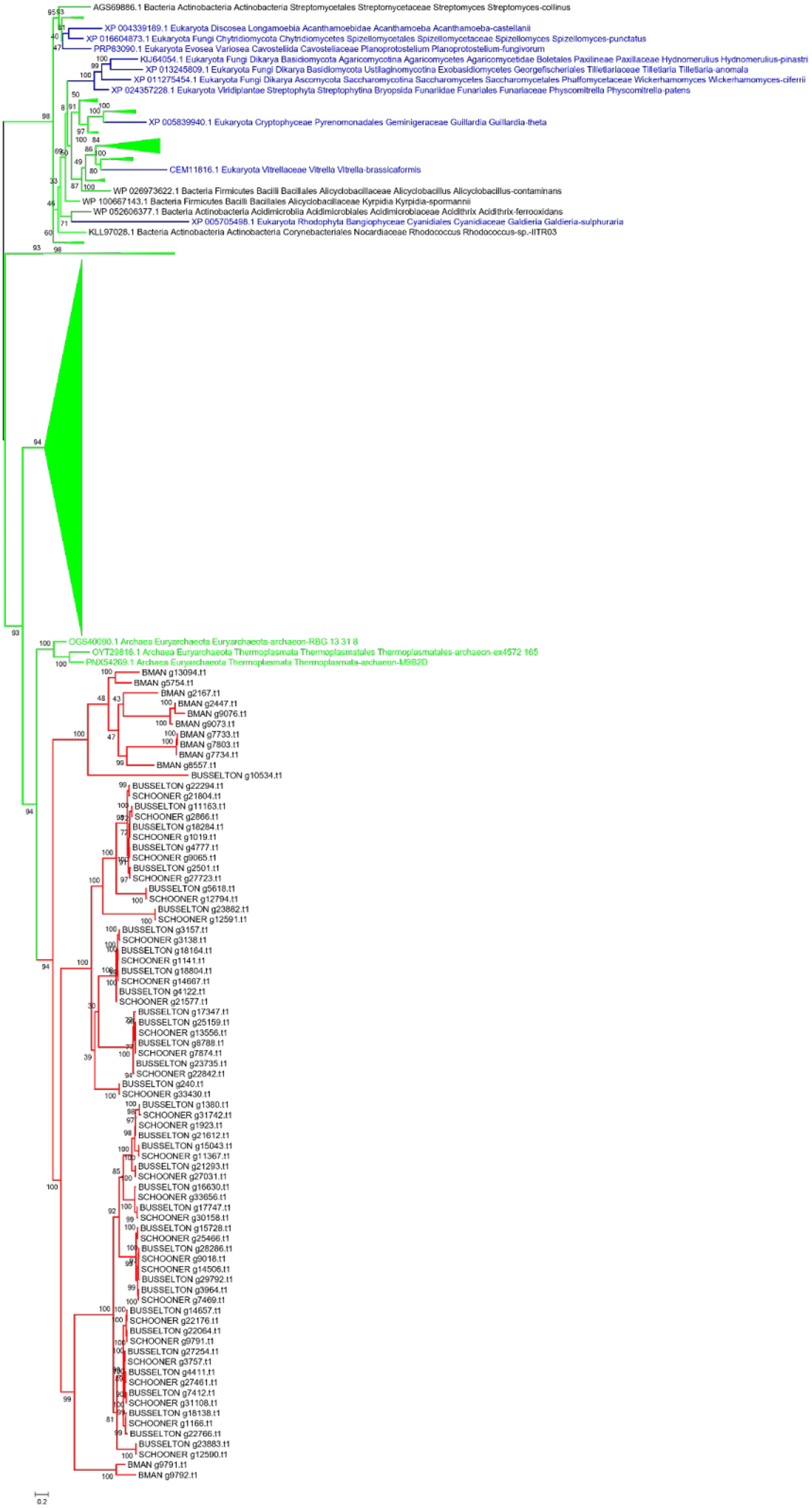
AGS69886.1 Bacteria Actinobacteria Actinobacteria Streptomycetales Streptomycetaceae Streptomyces Streptomyces-collinus
XP 004339189.1 Eukaryota Discosea Longamoebia Acanthamoebidae Acanthamoeba Acanthamoeba-castellanii
XP 016604873.1 Eukaryota Fungi Chytridiomycota Chytridiomycetes Spizellomycetales Spizellomycetaceae Spizellomyces Spizellomyces-punctatus
PRP83090.1 Eukaryota Evosea Variosea Cavosteliida Cavosteliaceae Planoprotostelium Planoprotostelium-fungivorum
KIJ64054.1 Eukaryota Fungi Dikarya Basidiomycota Agaricomycotina Agaricomycetes Agaricomycetidae Boletales Paxilineae Paxillaceae Hydnomerulius Hydnomerulius-pinastri
XP 013245809.1 Eukaryota Fungi Dikarya Basidiomycota Ustilaginomycotina Exobasidiomycetes Georgefischeriales Tilletiariaceae Tilletiaria Tilletiaria-anomala
XP 011275454.1 Eukaryota Fungi Dikarya Ascomycota Saccharomycotina Saccharomycetes Saccharomycetales Phaffomycetaceae Wickerhamomyces Wickerhamomyces-ciferrii
XP 024357228.1 Eukaryota Viridiplantae Streptophyta Streptophytina Bryopsida Funariidae Funariales Funariaceae Physcomitrella Physcomitrella-patens
XP 005839940.1 Eukaryota Cryptophyceae Pyrenomonadales Geminigeraceae Guillardia Guillardia-theta
CEM11816.1 Eukaryota Vitrellaceae Vitrella Vitrella-brassicaformis
WP 026973622.1 Bacteria Firmicutes Bacilli Bacillales Alicyclobacillaceae Alicyclobacillus Alicyclobacillus-contaminans
WP 100667143.1 Bacteria Firmicutes Bacilli Bacillales Alicyclobacillaceae Kyrpidia Kyrpidia-spormannii
WP 052606377.1 Bacteria Actinobacteria Acidimicrobiia Acidimicrobiales Acidimicrobiaceae Acidithrix Acidithrix-ferrooxidans
XP 005705498.1 Eukaryota Rhodophyta Bangiophyceae Cyanidiales Cyanidiaceae Galdieria Galdieria-sulphuraria
KLL97028.1 Bacteria Actinobacteria Actinobacteria Corynebacteriales Nocardiaceae Rhodococcus Rhodococcus-sp.-IITR03
OGS40090.1 Archaea Euryarchaeota Euryarchaeota-archaeon-RBG 13 31 8
OYT29816.1 Archaea Euryarchaeota Thermoplasmata Thermoplasmatales Thermoplasmatales-archaeon-ex4572 165
PNX54269.1 Archaea Euryarchaeota Thermoplasmata Thermoplasmata-archaeon-M9B2D
BMAN g13094.t1
BMAN g5754.t1
BMAN g2167.t1
BMAN g2447.t1
BMAN g9076.t1
BMAN g9073.t1
BMAN g7733.t1
BMAN g7803.t1
BMAN g7734.t1
BMAN g8557.t1
BUSSELTON g10534.t1
BUSSELTON g22294.t1
SCHOONER g21804.t1
BUSSELTON g11163.t1
SCHOONER g2866.t1
BUSSELTON g18284.t1
SCHOONER g1019.t1
BUSSELTON g4777.t1
SCHOONER g9065.t1
BUSSELTON g2501.t1
SCHOONER g27723.t1
BUSSELTON g5618.t1
SCHOONER g12794.t1
BUSSELTON g23882.t1
SCHOONER g12591.t1
BUSSELTON g3157.t1
SCHOONER g3138.t1
BUSSELTON g18164.t1
SCHOONER g1141.t1
BUSSELTON g18804.t1
SCHOONER g14667.t1
BUSSELTON g4122.t1
SCHOONER g21577.t1
BUSSELTON g17347.t1
BUSSELTON g25159.t1
SCHOONER g13556.t1
BUSSELTON g8788.t1
SCHOONER g7874.t1
BUSSELTON g23735.t1
SCHOONER g22842.t1
BUSSELTON g240.t1
SCHOONER g33430.t1
BUSSELTON g1380.t1
SCHOONER g31742.t1
SCHOONER g1923.t1
BUSSELTON g21612.t1
BUSSELTON g15043.t1
SCHOONER g11367.t1
BUSSELTON g21293.t1
SCHOONER g27031.t1
BUSSELTON g16630.t1
SCHOONER g33656.t1
BUSSELTON g17747.t1
SCHOONER g30158.t1
BUSSELTON g15728.t1
SCHOONER g25466.t1
BUSSELTON g28286.t1
SCHOONER g9018.t1
SCHOONER g14506.t1
BUSSELTON g29792.t1
BUSSELTON g3964.t1
SCHOONER g7469.t1
BUSSELTON g14657.t1
SCHOONER g22176.t1
BUSSELTON g22064.t1
SCHOONER g9791.t1
BUSSELTON g27254.t1
SCHOONER g3757.t1
BUSSELTON g4411.t1
SCHOONER g27461.t1
BUSSELTON g7412.t1
SCHOONER g31108.t1
BUSSELTON g18138.t1
SCHOONER g1166.t1
BUSSELTON g22766.t1
BUSSELTON g23883.t1
SCHOONER g12590.t1
BMAN g9791.t1
BMAN g9792.t1
0.2

**Figure S13F** Phylogenetic reconstruction of Acetate transporters. Alignment with MAFFT, site selection BMGE and phylogeny IQtree model C20+G4 1000 ultrafast bootstrap. *Anaeramoeba* sequences are in red, eukaryotic sequences in blue, and prokaryotic sequences in bright green.

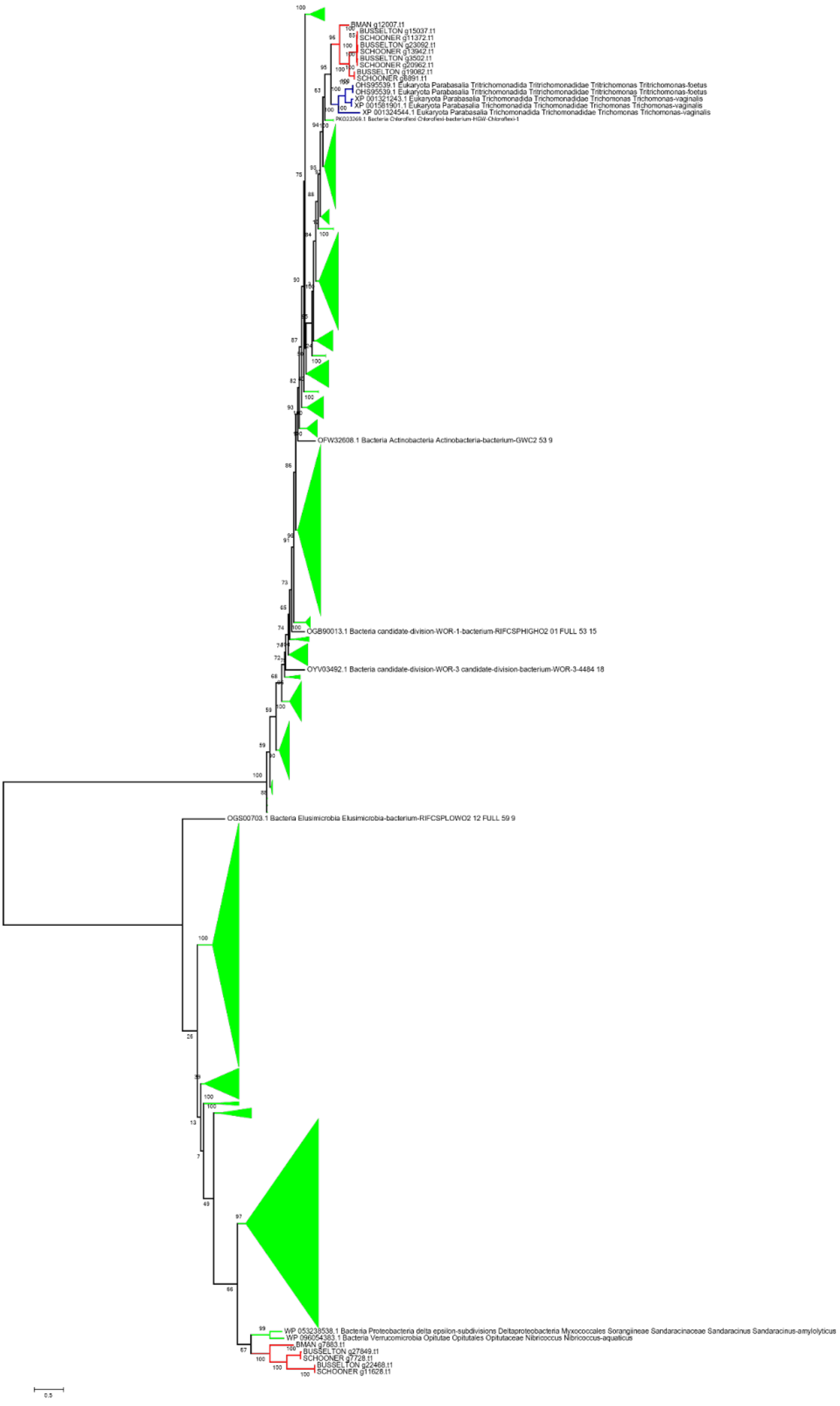
BMAN g12007.t1
BUSSELTON g15037.t1
SCHOONER g11372.t1
BUSSELTON g23092.t1
SCHOONER g13942.t1
BUSSELTON g3502.t1
SCHOONER g20962.t1
BUSSELTON g19082.t1
SCHOONER g6891.t1
OHS95539.1 Eukaryota Parabasalia Tritrichomonadida Tritrichomonadidae Tritrichomonas Tritrichomonas-foetus
OHS95539.1 Eukaryota Parabasalia Tritrichomonadida Tritrichomonadidae Tritrichomonas Tritrichomonas-foetus
XP 001321243.1 Eukaryota Parabasalia Trichomonadida Trichomonadidae Trichomonas Trichomonas-vaginalis
XP 001581901.1 Eukaryota Parabasalia Trichomonadida Trichomonadidae Trichomonas Trichomonas-vaginalis
XP 001324544.1 Eukaryota Parabasalia Trichomonadida Trichomonadidae Trichomonas Trichomonas-vaginalis
PKO23269.1 Bacteria Chloroflexi Chloroflexi-bacterium-HGW-Chloroflexi-1
OFW32608.1 Bacteria Actinobacteria Actinobacteria-bacterium-GWC2 53 9
OGB90013.1 Bacteria candidate-division-WOR-1-bacterium-RIFCSPHIGHO2 01 FULL 53 15
OYV03492.1 Bacteria candidate-division-WOR-3 candidate-division-bacterium-WOR-3-4484 18
OGS00703.1 Bacteria Elusimicrobia Elusimicrobia-bacterium-RIFCSPLOWO2 12 FULL 59 9
WP 053238538.1 Bacteria Proteobacteria delta epsilon-subdivisions Deltaproteobacteria Myxococcales Sorangiineae Sandaracinaceae Sandaracinus Sandaracinus-amylolyticus
WP 096054383.1 Bacteria Verrucomicrobia Opitutae Opitutales Opitutaceae Nibricoccus Nibricoccus-aquaticus
BMAN g7883.t1
BUSSELTON g27849.t1
SCHOONER g7728.t1
BUSSELTON g22468.t1
SCHOONER g11628.t1
0.5

**Figure S13G** Phylogenetic reconstruction of Pyruvate phosphate dikinase. Alignment with MAFFT, site selection BMGE and phylogeny IQtree model LG4X 1000 ultrafast bootstrap. *Anaeramoeba* sequences are in red, eukaryotic sequences in blue, and prokaryotic sequences in bright green.

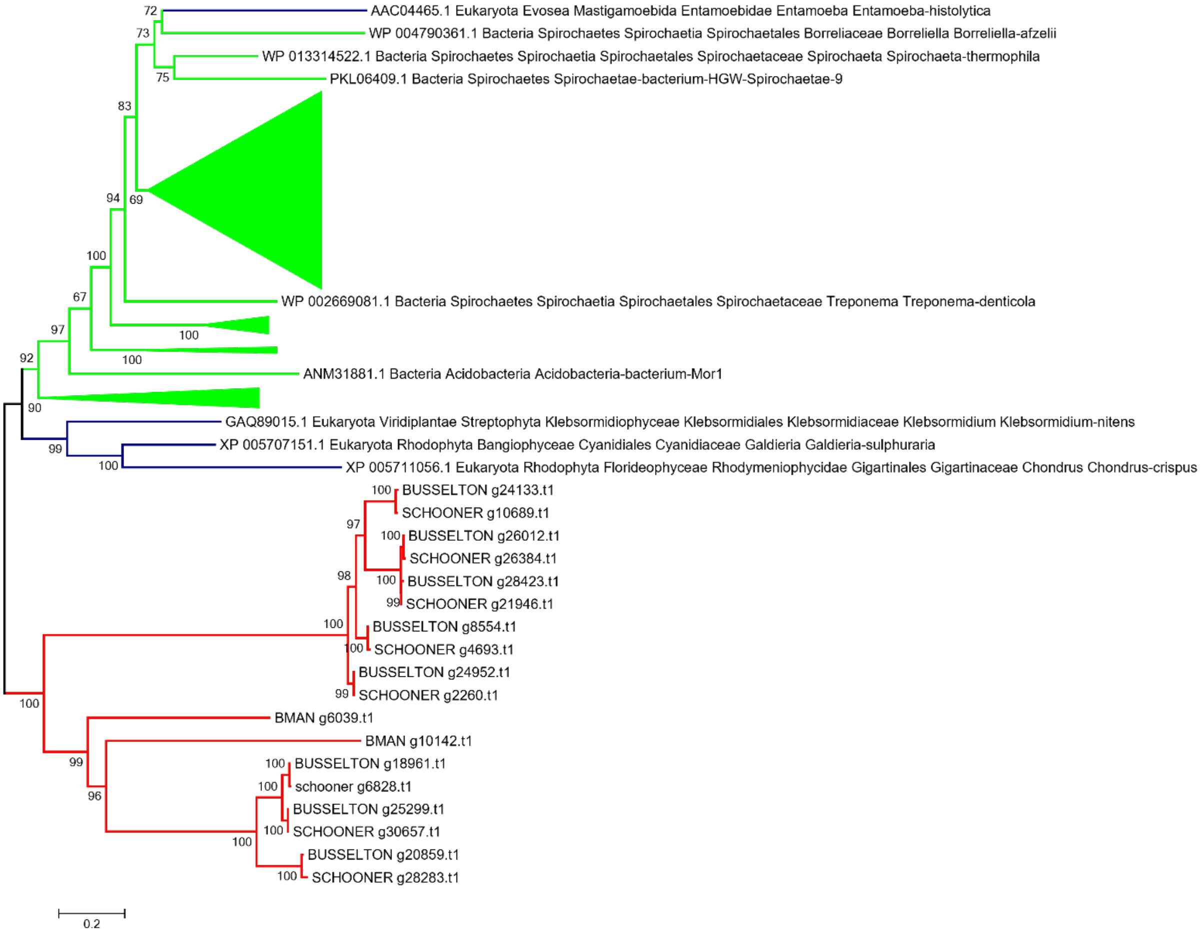


**Figure S13H** Phylogenetic reconstruction of ATP-independent phosphofructokinase. Alignment with MAFFT, site selection BMGE and phylogeny IQtree model LG4X 1000 ultrafast bootstrap. *Anaeramoeba* sequences are in red, eukaryotic sequences in blue, and prokaryotic sequences in bright green.

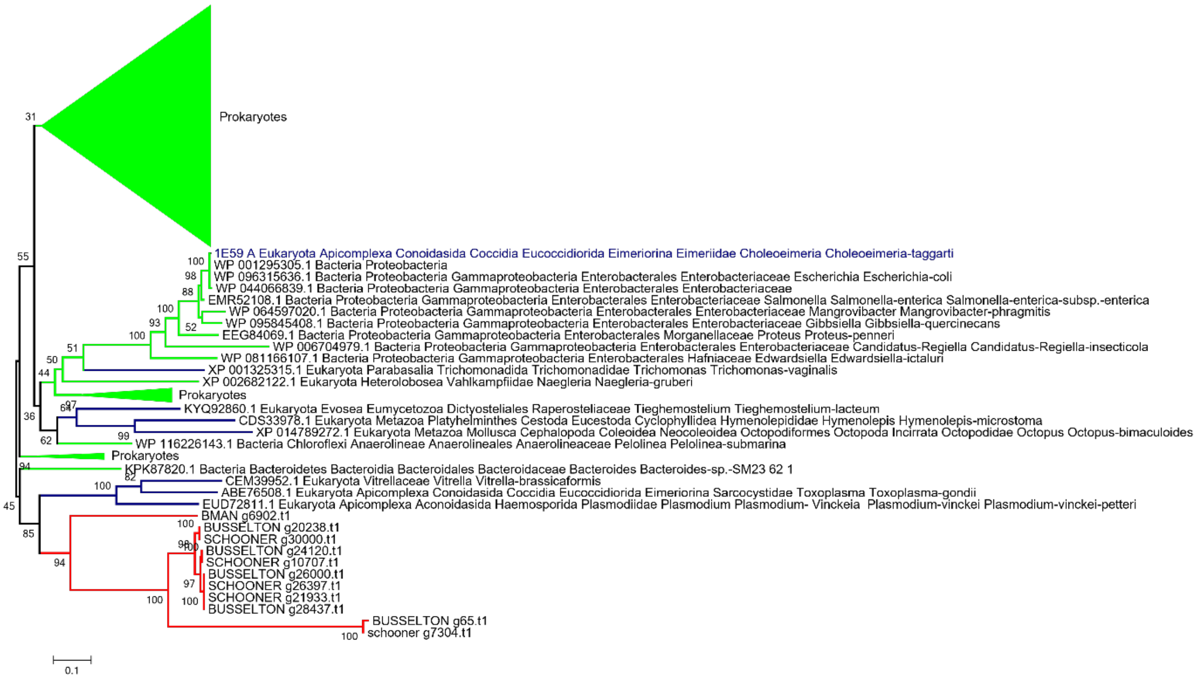


**Figure S13I** Phylogenetic reconstruction of Phosphoglycerate mutase (OG 363). Alignment with MAFFT, site selection BMGE and phylogeny IQtree model LG4X 1000 ultrafast bootstrap. *Anaeramoeba* sequences are in red, eukaryotic sequences in blue, and prokaryotic sequences in bright green.

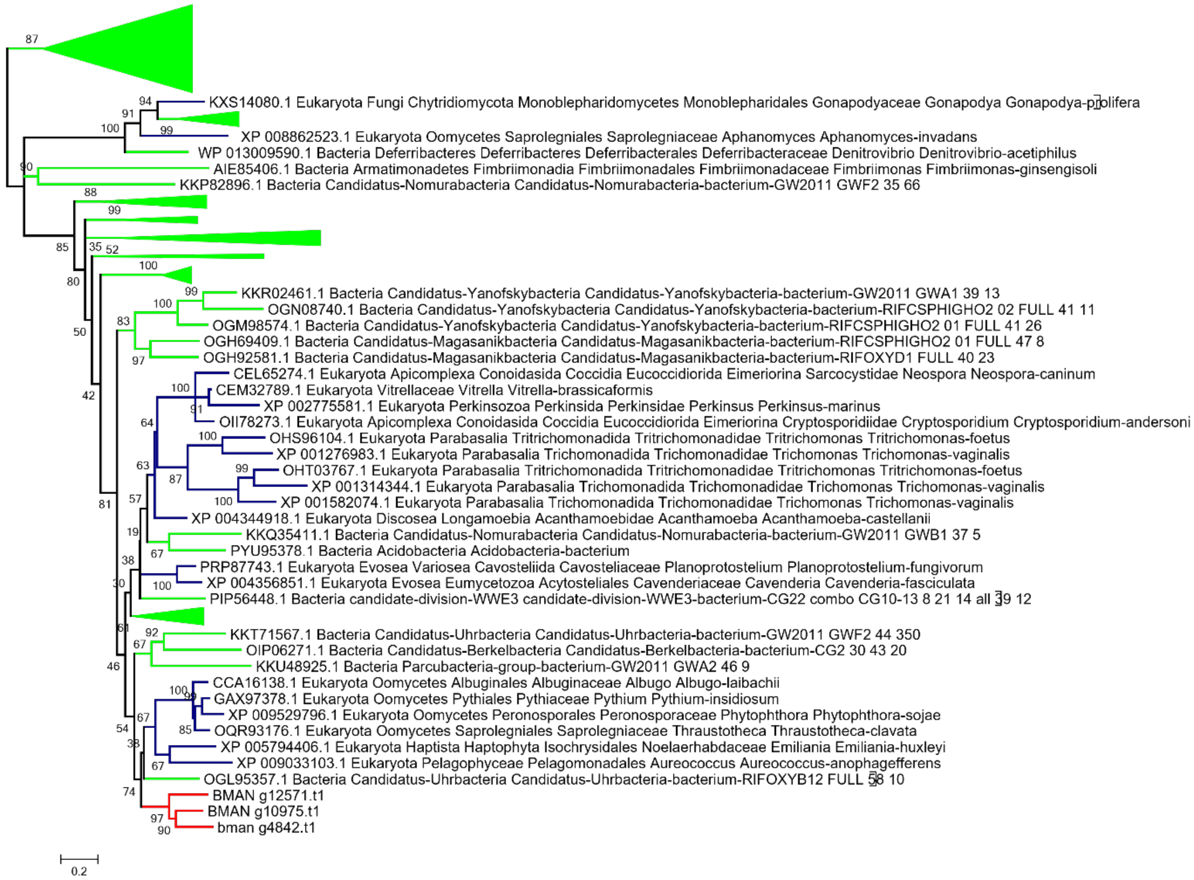


**Figure S13J** Phylogenetic reconstruction of Phosphoglycerate mutase (OG 2532). Alignment with MAFFT, site selection BMGE and phylogeny IQtree model LG4X,1,000 ultrafast bootstrap. *Anaeramoeba* sequences are in red, eukaryotic sequences in blue, and prokaryotic sequences in bright green.

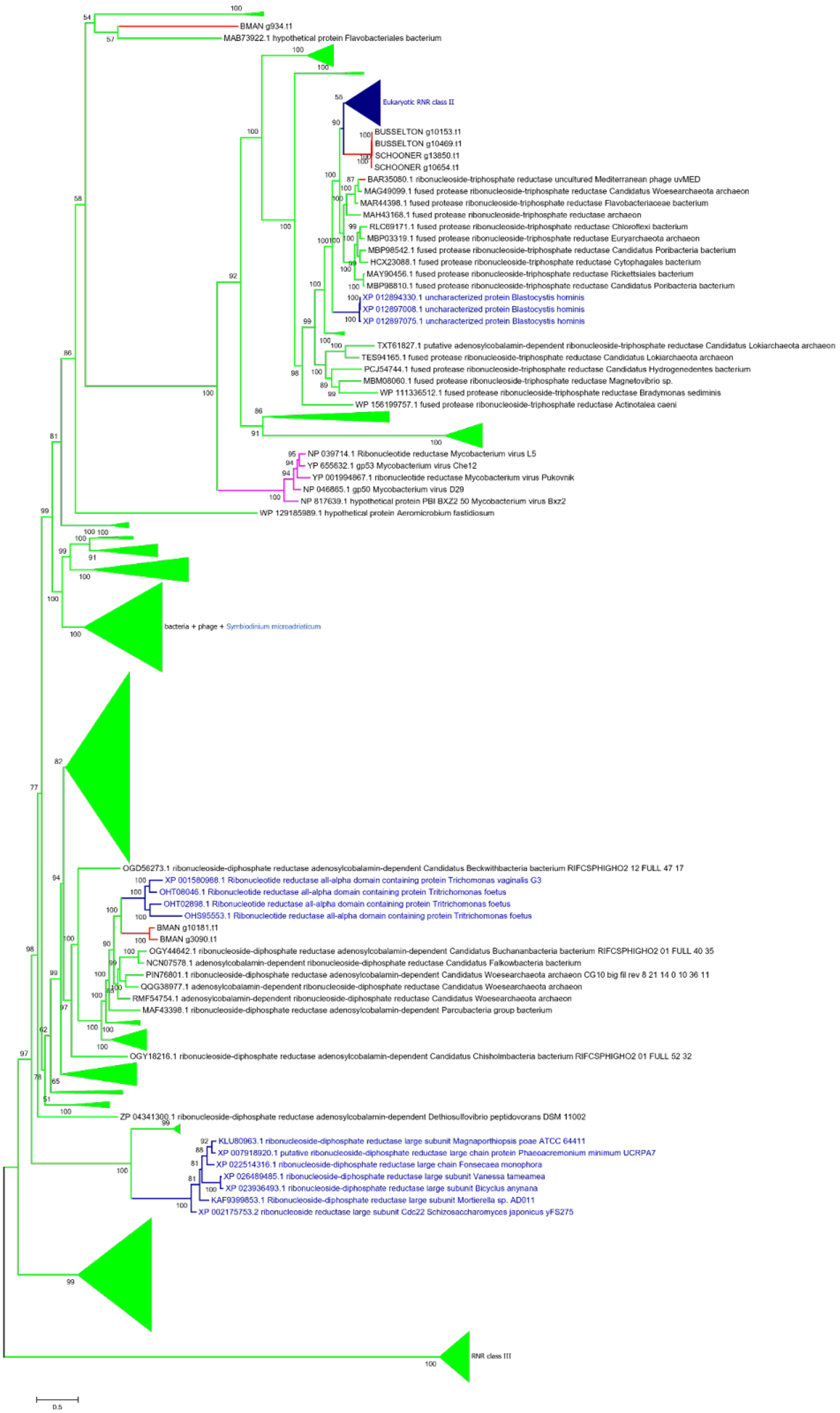
BMAN g934.t1
MAB73922.1 hypothetical protein Flavobacteriales bacterium
Eukaryotic RNR class II
BUSSELTON g10153.t1
BUSSELTON g10469.t1
SCHOONER g13850.t1
SCHOONER g10654.t1
BAR35080.1 ribonucleoside-triphosphate reductase uncultured Mediterranean phage uvMED
MAG49099.1 fused protease ribonucleoside-triphosphate reductase Candidatus Woesearchaeota archaeon
MAR44398.1 fused protease ribonucleoside-triphosphate reductase Flavobacteriaceae bacterium
MAH43168.1 fused protease ribonucleoside-triphosphate reductase archaeon
RLC69171.1 fused protease ribonucleoside-triphosphate reductase Chloroflexi bacterium
MBP03319.1 fused protease ribonucleoside-triphosphate reductase Euryarchaeota archaeon
MBP98542.1 fused protease ribonucleoside-triphosphate reductase Candidatus Poribacteria bacterium
HCX23088.1 fused protease ribonucleoside-triphosphate reductase Cytophagales bacterium
MAY90456.1 fused protease ribonucleoside-triphosphate reductase Rickettsiales bacterium
MBP98810.1 fused protease ribonucleoside-triphosphate reductase Candidatus Poribacteria bacterium
XP 012894330.1 uncharacterized protein Blastocystis hominis
XP 012897008.1 uncharacterized protein Blastocystis hominis
XP 012897075.1 uncharacterized protein Blastocystis hominis
TXT61827.1 putative adenosylcobalamin-dependent ribonucleoside-triphosphate reductase Candidatus Lokiarchaeota archaeon
TES94165.1 fused protease ribonucleoside-triphosphate reductase Candidatus Lokiarchaeota archaeon
PCJ54744.1 fused protease ribonucleoside-triphosphate reductase Candidatus Hydrogenedentes bacterium
MBM08060.1 fused protease ribonucleoside-triphosphate reductase Magnetovibrio sp.
WP 111336512.1 fused protease ribonucleoside-triphosphate reductase Bradymonas sediminis
WP 156199757.1 fused protease ribonucleoside-triphosphate reductase Actinotalea caeni
NP 039714.1 Ribonucleotide reductase Mycobacterium virus L5
YP 655632.1 gp53 Mycobacterium virus Che12
YP 001994867.1 ribonucleotide reductase Mycobacterium virus Pukovnik
NP 046865.1 gp50 Mycobacterium virus D29
NP 817639.1 hypothetical protein PBI BXZ2 50 Mycobacterium virus Bxz2
WP 129185989.1 hypothetical protein Aeromicrobium fastidiosum
bacteria + phage + Symbiodinium microadriaticum
OGD56273.1 ribonucleoside-diphosphate reductase adenosylcobalamin-dependent Candidatus Beckwithbacteria bacterium RIFCSPHIGHO2 12 FULL 47 17
XP 001580988.1 Ribonucleotide reductase all-alpha domain containing protein Trichomonas vaginalis G3
OHT08046.1 Ribonucleotide reductase all-alpha domain containing protein Tritrichomonas foetus
OHT02898.1 Ribonucleotide reductase all-alpha domain containing protein Tritrichomonas foetus
OHS95553.1 Ribonucleotide reductase all-alpha domain containing protein Tritrichomonas foetus
BMAN g10181.t1
BMAN g3090.t1
OGY44642.1 ribonucleoside-diphosphate reductase adenosylcobalamin-dependent Candidatus Buchananbacteria bacterium RIFCSPHIGHO2 01 FULL 40 35
NCN07578.1 adenosylcobalamin-dependent ribonucleoside-diphosphate reductase Candidatus Falkowbacteria bacterium
PIN76801.1 ribonucleoside-diphosphate reductase adenosylcobalamin-dependent Candidatus Woesearchaeota archaeon CG10 big fil rev 8 21 14 0 10 36 11
QQG38977.1 adenosylcobalamin-dependent ribonucleoside-diphosphate reductase Candidatus Woesearchaeota archaeon
RMF54754.1 adenosylcobalamin-dependent ribonucleoside-diphosphate reductase Candidatus Woesearchaeota archaeon
MAF43398.1 ribonucleoside-diphosphate reductase adenosylcobalamin-dependent Parcubacteria group bacterium
OGY18216.1 ribonucleoside-diphosphate reductase adenosylcobalamin-dependent Candidatus Chisholmbacteria bacterium RIFCSPHIGHO2 01 FULL 52 32
ZP 04341300.1 ribonucleoside-diphosphate reductase adenosylcobalamin-dependent Dethiosulfovibrio peptidovorans DSM 11002
KLU80963.1 ribonucleoside-diphosphate reductase large subunit Magnaporthiopsis poae ATCC 64411
XP 007918920.1 putative ribonucleoside-diphosphate reductase large chain protein Phaeoacremonium minimum UCRPA7
XP 022514316.1 ribonucleoside-diphosphate reductase large chain Fonsecaea monophora
XP 026489485.1 ribonucleoside-diphosphate reductase large subunit Vanessa tameamea
XP 023936493.1 ribonucleoside-diphosphate reductase large subunit Bicyclus anynana
KAF9399853.1 Ribonucleoside-diphosphate reductase large subunit Mortierella sp. AD011
XP 002175753.2 ribonucleoside reductase large subunit Cdc22 Schizosaccharomyces japonicus yFS275
RNR class III
0.5

**Figure 13K** Phylogenetic reconstruction of Ribonucleotide reductases. Alignment with MAFFT, site selection BMGE and phylogeny IQtree model C20+G4 1000 ultrafast bootstrap. *Anaeramoeba* sequences are in red, eukaryotic sequences in blue, viral sequences in light purple and prokaryotic sequences in bright green.

.

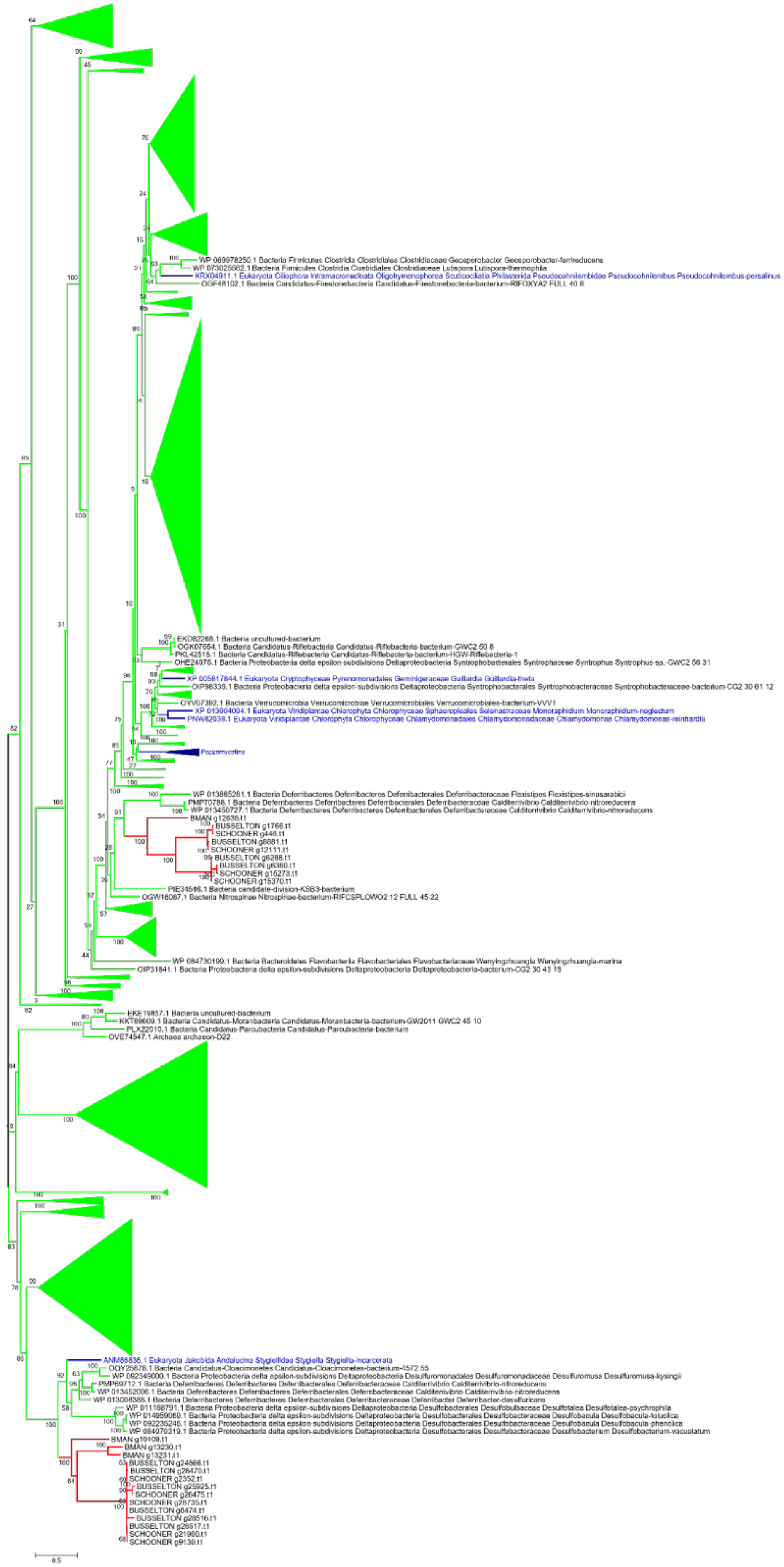

WP 069978250.1 Bacteria Firmicutes Clostridia Clostridiales Clostridiaceae Geosporobacter Geosporobacter-ferrireducens
WP 073025562.1 Bacteria Firmicutes Clostridia Clostridiales Clostridiaceae Lutispora Lutispora-thermophila
KRX04911.1 Eukaryota Ciliophora Intramacronucleata Oligohymenophorea Scuticociliatia Philasterida Pseudocohnilembidae Pseudocohnilembus Pseudocohnilembus-persalinus
OGF48102.1 Bacteria Candidatus-Firestonebacteria Candidatus-Firestonebacteria-bacterium-RIFOXYA2 FULL 40 8
EKD82268.1 Bacteria uncultured-bacterium
OGK07654.1 Bacteria Candidatus-Riflebacteria Candidatus-Riflebacteria-bacterium-GWC2 50 8
PKL42515.1 Bacteria Candidatus-Riflebacteria Candidatus-Riflebacteria-bacterium-HGW-Riflebacteria-1
OHE24075.1 Bacteria Proteobacteria delta epsilon-subdivisions Deltaproteobacteria Syntrophobacterales Syntrophaceae Syntrophus Syntrophus-sp.-GWC2 56 31
XP 005817844.1 Eukaryota Cryptophyceae Pyrenomonadales Geminigeraceae Guillardia Guillardia-theta
OIP98335.1 Bacteria Proteobacteria delta epsilon-subdivisions Deltaproteobacteria Syntrophobacterales Syntrophobacteraceae Syntrophobacteraceae-bacterium-CG2 30 61 12
OYV07392.1 Bacteria Verrucomicrobia Verrucomicrobiae Verrucomicrobiales Verrucomicrobiales-bacterium-VVV1
XP 013904094.1 Eukaryota Viridiplantae Chlorophyta Chlorophyceae Sphaeropleales Selenastraceae Monoraphidium Monoraphidium-neglectum
PNW82038.1 Eukaryota Viridiplantae Chlorophyta Chlorophyceae Chlamydomonadales Chlamydomonadaceae Chlamydomonas Chlamydomonas-reinhardtii
Pezizomycotina
WP 013885281.1 Bacteria Deferribacteres Deferribacteres Deferribacterales Deferribacteraceae Flexistipes Flexistipes-sinusarabici
PMP70788.1 Bacteria Deferribacteres Deferribacteres Deferribacterales Deferribacteraceae Calditerrivibrio Calditerrivibrio-nitroreducens
WP 013450727.1 Bacteria Deferribacteres Deferribacteres Deferribacterales Deferribacteraceae Calditerrivibrio Calditerrivibrio-nitroreducens
BMAN g12635.t1
BUSSELTON g1766.t1
SCHOONER g448.t1
BUSSELTON g8881.t1
SCHOONER g12111.t1
BUSSELTON g6288.t1
BUSSELTON g6380.t1
SCHOONER g15273.t1
SCHOONER g15370.t1
PIE34546.1 Bacteria candidate-division-KSB3-bacterium
OGW16067.1 Bacteria Nitrospinae Nitrospinae-bacterium-RIFCSPLOWO2 12 FULL 45 22
WP 084730199.1 Bacteria Bacteroidetes Flavobacteriia Flavobacteriales Flavobacteriaceae Wenyingzhuangia Wenyingzhuangia-marina
OIP31841.1 Bacteria Proteobacteria delta epsilon-subdivisions Deltaproteobacteria Deltaproteobacteria-bacterium-CG2 30 43 15
EKE19857.1 Bacteria uncultured-bacterium
KKT89609.1 Bacteria Candidatus-Moranbacteria Candidatus-Moranbacteria-bacterium-GW2011 GWC2 45 10
PLX22010.1 Bacteria Candidatus-Parcubacteria Candidatus-Parcubacteria-bacterium
OVE74547.1 Archaea archaeon-D22
ANM86836.1 Eukaryota Jakobida Andalucina Stygiellidae Stygiella Stygiella-incarcerata
OQY25878.1 Bacteria Candidatus-Cloacimonetes Candidatus-Cloacimonetes-bacterium-4572 55
WP 092349000.1 Bacteria Proteobacteria delta epsilon-subdivisions Deltaproteobacteria Desulfuromonadales Desulfuromonadaceae Desulfuromusa Desulfuromusa-kysingii
PMP69712.1 Bacteria Deferribacteres Deferribacteres Deferribacterales Deferribacteraceae Calditerrivibrio Calditerrivibrio-nitroreducens
WP 013452006.1 Bacteria Deferribacteres Deferribacteres Deferribacterales Deferribacteraceae Calditerrivibrio Calditerrivibrio-nitroreducens
WP 013008388.1 Bacteria Deferribacteres Deferribacteres Deferribacterales Deferribacteraceae Deferribacter Deferribacter-desulfuricans
WP 011188791.1 Bacteria Proteobacteria delta epsilon-subdivisions Deltaproteobacteria Desulfobacterales Desulfobulbaceae Desulfotalea Desulfotalea-psychrophila
WP 014959069.1 Bacteria Proteobacteria delta epsilon-subdivisions Deltaproteobacteria Desulfobacterales Desulfobacteraceae Desulfobacula Desulfobacula-toluolica
WP 092235246.1 Bacteria Proteobacteria delta epsilon-subdivisions Deltaproteobacteria Desulfobacterales Desulfobacteraceae Desulfobacula Desulfobacula-phenolica
WP 084070319.1 Bacteria Proteobacteria delta epsilon-subdivisions Deltaproteobacteria Desulfobacterales Desulfobacteraceae Desulfobacterium Desulfobacterium-vacuolatum
BMAN g10409.t1
BMAN g13230.t1
BMAN g13231.t1
BUSSELTON g24866.t1
BUSSELTON g28470.t1
SCHOONER g2352.t1
BUSSELTON g25925.t1
SCHOONER g26475.t1
SCHOONER g28735.t1
BUSSELTON g8474.t1
BUSSELTON g28516.t1
BUSSELTON g28517.t1
SCHOONER g21900.t1
SCHOONER g9130.t1
0.5

**Figure S13L** Phylogenetic reconstruction of NADH oxidase. Alignment with MAFFT, site selection BMGE, and phylogeny IQtree model C20+G4,1,000 ultrafast bootstrap. *Anaeramoeba* sequences are in red, eukaryotic sequences in blue, and prokaryotic sequences in bright green.

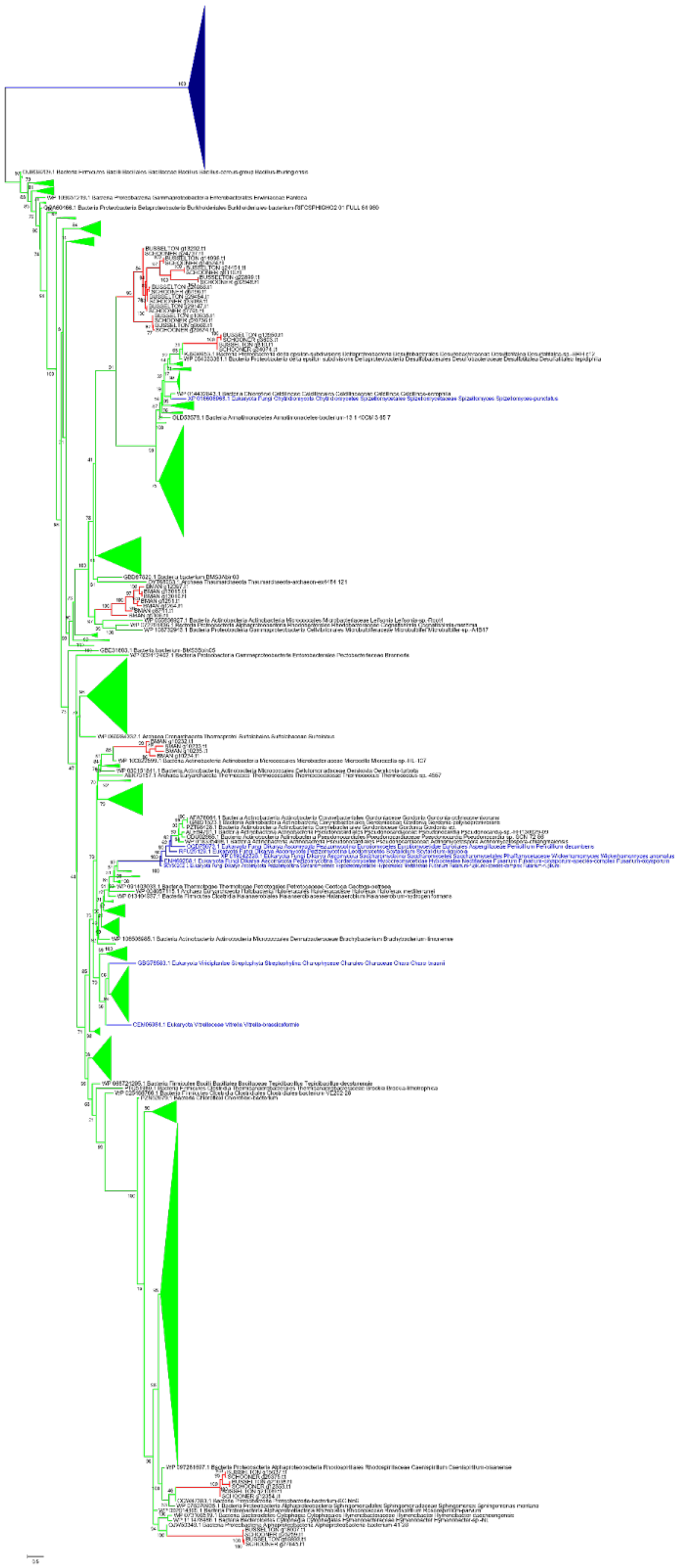

**Figure S13M** Phylogenetic reconstruction of OsmC. Alignment with MAFFT, site selection BMGE and phylogeny IQtree model LG4X 1000 ultrafast bootstrap. *Anaeramoeba* sequences are in red, Eukaryotic sequences in blue and prokaryotic sequences in bright green.

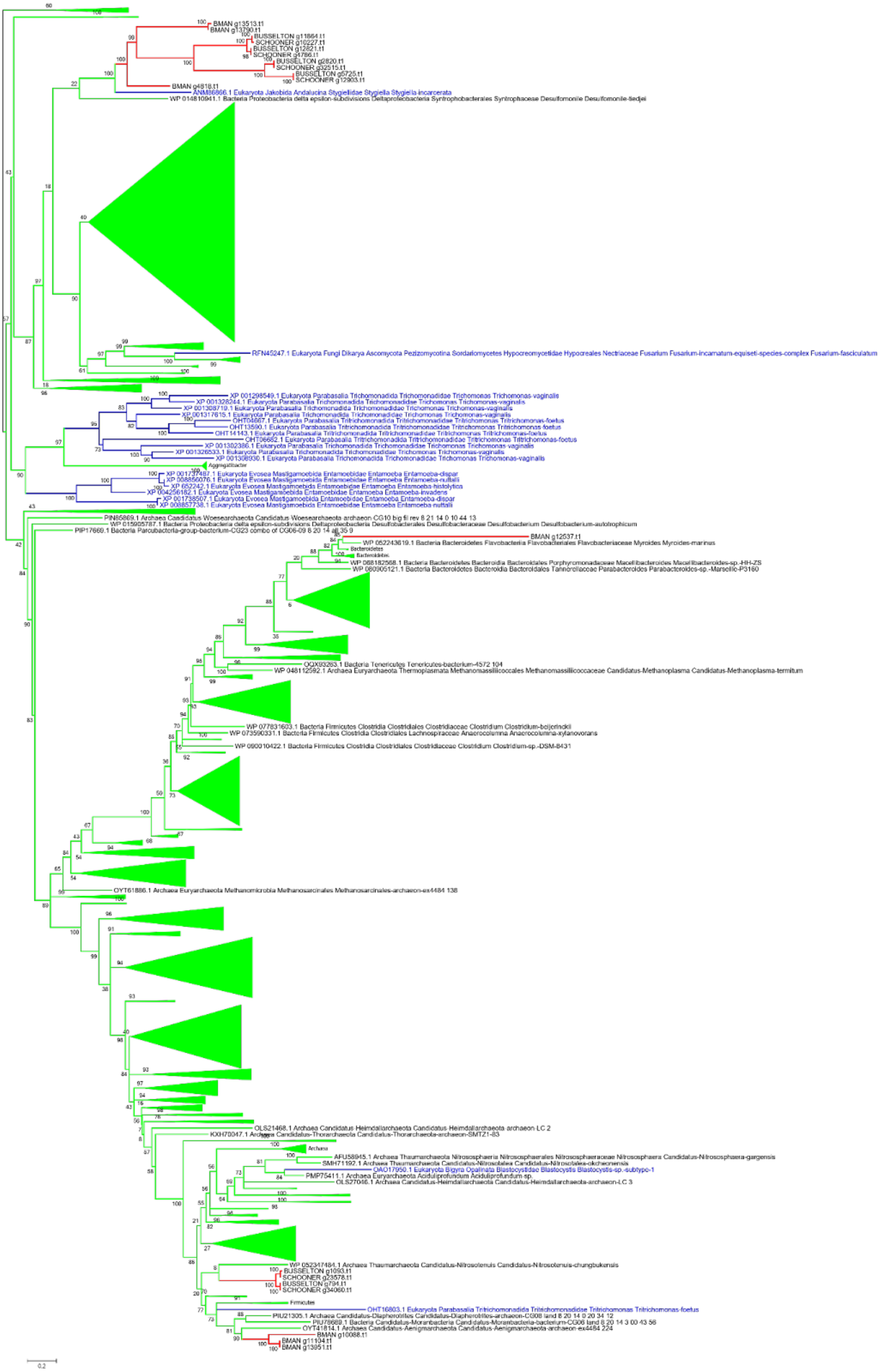

BMAN g13513.t1
BMAN g13790.t1
BUSSELTON g11864.t1
SCHOONER g10227.t1
BUSSELTON g12821.t1
SCHOONER g4786.t1
BUSSELTON g2820.t1
SCHOONER g32515.t1
BUSSELTON g5725.t1
SCHOONER g12903.t1
BMAN g4818.t1
ANM86866.1 Eukaryota Jakobida Andalucina Stygiellidae Stygiella Stygiella-incarcerata
WP 014810941.1 Bacteria Proteobacteria delta epsilon-subdivisions Deltaproteobacteria Syntrophobacterales Syntrophaceae Desulfomonile Desulfomonile-tiedjei
RFN45247.1 Eukaryota Fungi Dikarya Ascomycota Pezizomycotina Sordariomycetes Hypocreomycetidae Hypocreales Nectriaceae Fusarium Fusarium-incarnatum-equiseti-species-complex Fusarium-fasciculatum
XP 001298549.1 Eukaryota Parabasalia Trichomonadida Trichomonadidae Trichomonas Trichomonas-vaginalis
XP 001328244.1 Eukaryota Parabasalia Trichomonadida Trichomonadidae Trichomonas Trichomonas-vaginalis
XP 001308719.1 Eukaryota Parabasalia Trichomonadida Trichomonadidae Trichomonas Trichomonas-vaginalis
XP 001317615.1 Eukaryota Parabasalia Trichomonadida Trichomonadidae Trichomonas Trichomonas-vaginalis
OHT04667.1 Eukaryota Parabasalia Tritrichomonadida Tritrichomonadidae Tritrichomonas Tritrichomonas-foetus
OHT13590.1 Eukaryota Parabasalia Tritrichomonadida Tritrichomonadidae Tritrichomonas Tritrichomonas-foetus
OHT14143.1 Eukaryota Parabasalia Tritrichomonadida Tritrichomonadidae Tritrichomonas Tritrichomonas-foetus
OHT06682.1 Eukaryota Parabasalia Tritrichomonadida Tritrichomonadidae Tritrichomonas Tritrichomonas-foetus
XP 001302386.1 Eukaryota Parabasalia Trichomonadida Trichomonadidae Trichomonas Trichomonas-vaginalis
XP 001326533.1 Eukaryota Parabasalia Trichomonadida Trichomonadidae Trichomonas Trichomonas-vaginalis
XP 001308930.1 Eukaryota Parabasalia Trichomonadida Trichomonadidae Trichomonas Trichomonas-vaginalis
Aggregatibacter
XP 001737487.1 Eukaryota Evosea Mastigamoebida Entamoebidae Entamoeba Entamoeba-dispar
XP 008856076.1 Eukaryota Evosea Mastigamoebida Entamoebidae Entamoeba Entamoeba-nuttalli
XP 652242.1 Eukaryota Evosea Mastigamoebida Entamoebidae Entamoeba Entamoeba-histolytica
XP 004256182.1 Eukaryota Evosea Mastigamoebida Entamoebidae Entamoeba Entamoeba-invadens
XP 001738507.1 Eukaryota Evosea Mastigamoebida Entamoebidae Entamoeba Entamoeba-dispar
XP 008857738.1 Eukaryota Evosea Mastigamoebida Entamoebidae Entamoeba Entamoeba-nuttalli
PIN85869.1 Archaea Candidatus Woesearchaeota Candidatus Woesearchaeota archaeon CG10 big fil rev 8 21 14 0 10 44 13
WP 015905787.1 Bacteria Proteobacteria delta epsilon-subdivisions Deltaproteobacteria Desulfobacterales Desulfobacteraceae Desulfobacterium Desulfobacterium-autotrophicum
PIP17669.1 Bacteria Parcubacteria-group-bacterium-CG23 combo of CG06-09 8 20 14 all 35 9
BMAN g12537.t1
WP 052243619.1 Bacteria Bacteroidetes Flavobacteriia Flavobacteriales Flavobacteriaceae Myroides Myroides-marinus
Bacteroidetes
Bacteroidetes
WP 068182568.1 Bacteria Bacteroidetes Bacteroidia Bacteroidales Porphyromonadaceae Macellibacteroides Macellibacteroides-sp.-HH-ZS
WP 080905121.1 Bacteria Bacteroidetes Bacteroidia Bacteroidales Tannerellaceae Parabacteroides Parabacteroides-sp.-Marseille-P3160
OQX93263.1 Bacteria Tenericutes Tenericutes-bacterium-4572 104
WP 048112592.1 Archaea Euryarchaeota Thermoplasmata Methanomassiliicoccales Methanomassiliicoccaceae Candidatus-Methanoplasma Candidatus-Methanoplasma-termitum
WP 077831603.1 Bacteria Firmicutes Clostridia Clostridiales Clostridiaceae Clostridium Clostridium-beijerinckii
WP 073590331.1 Bacteria Firmicutes Clostridia Clostridiales Lachnospiraceae Anaerocolumna Anaerocolumna-xylanovorans
WP 090010422.1 Bacteria Firmicutes Clostridia Clostridiales Clostridiaceae Clostridium Clostridium-sp.-DSM-8431
OYT61886.1 Archaea Euryarchaeota Methanomicrobia Methanosarcinales Methanosarcinales-archaeon-ex4484 138
OLS21468.1 Archaea Candidatus Heimdallarchaeota Candidatus Heimdallarchaeota archaeon LC 2
KXH70047.1 Archaea Candidatus-Thorarchaeota Candidatus-Thorarchaeota-archaeon-SMTZ1-83
Archaea
AFU58945.1 Archaea Thaumarchaeota Nitrososphaeria Nitrososphaerales Nitrososphaeraceae Nitrososphaera Candidatus-Nitrososphaera-gargensis
SMH71192.1 Archaea Thaumarchaeota Candidatus-Nitrosotalea Candidatus-Nitrosotalea-okcheonensis
OAO17950.1 Eukaryota Bigyra Opalinata Blastocystidae Blastocystis Blastocystis-sp.-subtype-1
PMP75411.1 Archaea Euryarchaeota Aciduliprofundum Aciduliprofundum-sp.
OLS27046.1 Archaea Candidatus-Heimdallarchaeota Candidatus-Heimdallarchaeota-archaeon-LC 3
WP 052347484.1 Archaea Thaumarchaeota Candidatus-Nitrosotenuis Candidatus-Nitrosotenuis-chungbukensis
BUSSELTON g1093.t1
SCHOONER g23578.t1
BUSSELTON g794.t1
SCHOONER g34060.t1
Firmicutes
OHT16803.1 Eukaryota Parabasalia Tritrichomonadida Tritrichomonadidae Tritrichomonas Tritrichomonas-foetus
PIU21305.1 Archaea Candidatus-Diapherotrites Candidatus-Diapherotrites-archaeon-CG08 land 8 20 14 0 20 34 12
PIU78689.1 Bacteria Candidatus-Moranbacteria Candidatus-Moranbacteria-bacterium-CG06 land 8 20 14 3 00 43 56
OYT41814.1 Archaea Candidatus-Aenigmarchaeota Candidatus-Aenigmarchaeota-archaeon-ex4484 224
BMAN g10088.t1
BMAN g11104.t1
BMAN g13951.t1
0.2

**Figure S13N** Phylogenetic reconstruction of Nitroreductases. Alignment with MAFFT, site selection BMGE and phylogeny IQtree model LG4X 1000 ultrafast bootstrap. *Anaeramoeba* sequences are in red, Eukaryotic sequences in blue and prokaryotic sequences in bright green.

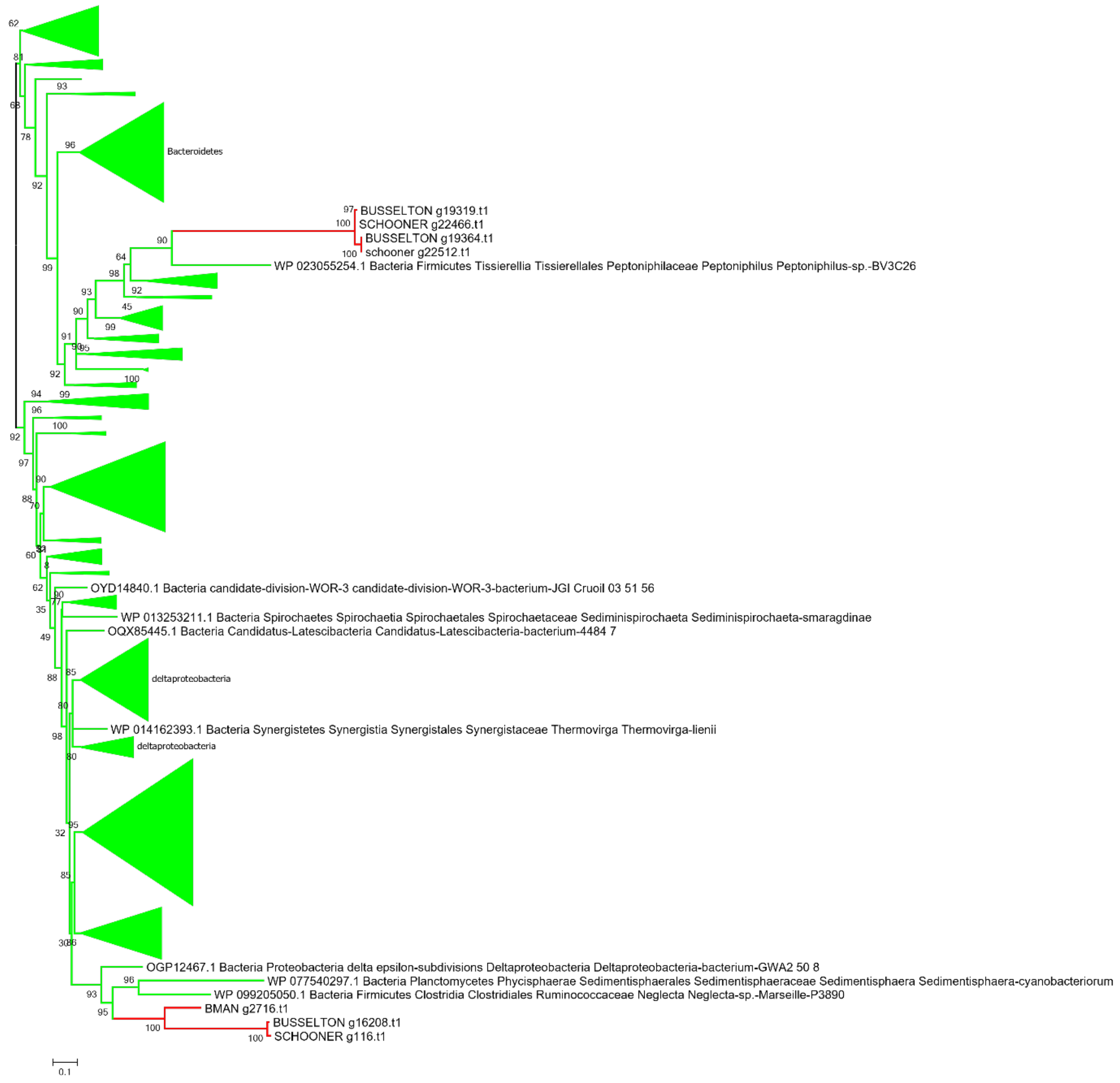


**Figure S13O** Phylogenetic reconstruction of Rubrerythrin. Alignment with MAFFT, site selection BMGE and phylogeny IQtree model LG4X 1000 ultrafast bootstrap. *Anaeramoeba* sequences are in red and prokaryotic sequences in bright green.

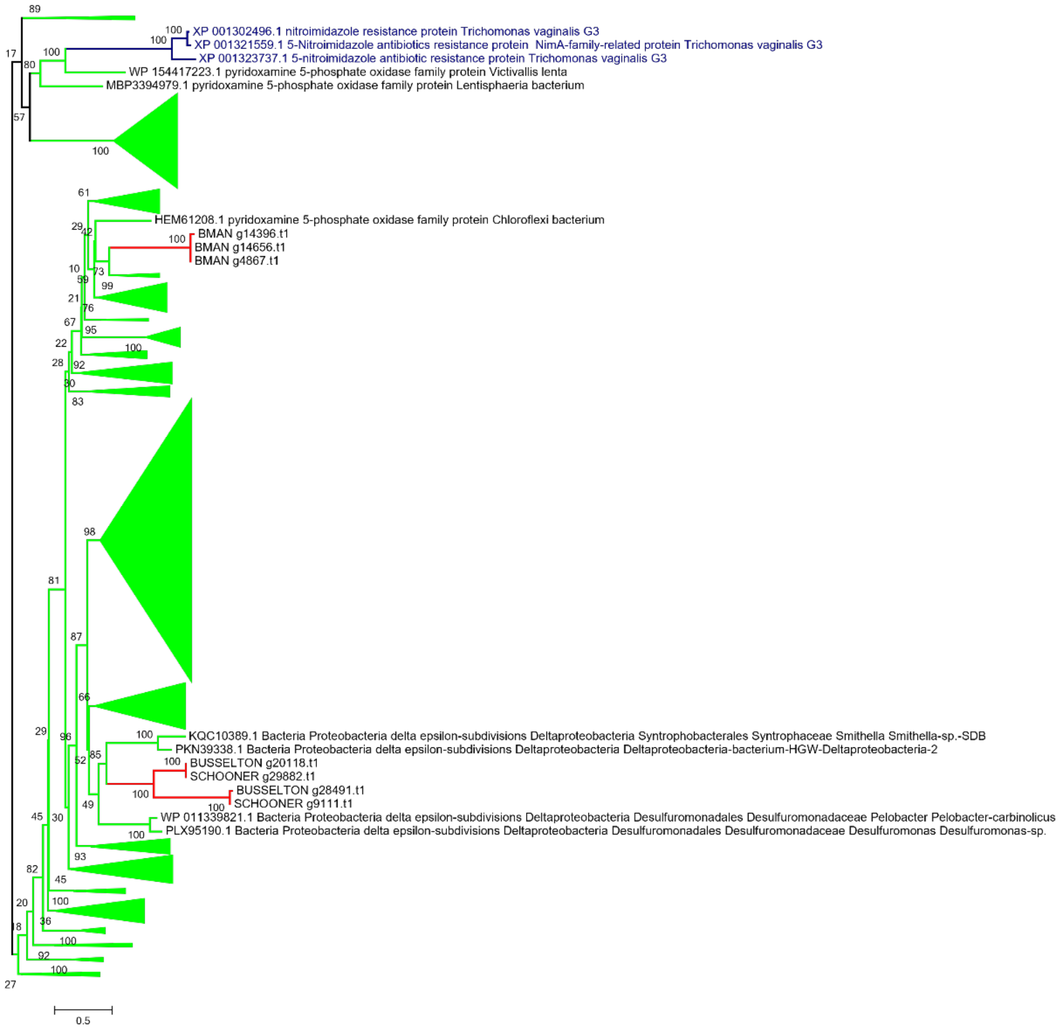


**Figure S13P** Phylogenetic reconstruction of 5-Nitroimidazole reductases. Alignment with MAFFT, site selection BMGE and phylogeny IQtree model LG4X 1000 ultrafast bootstrap. Anaeramoeba sequences are in red, eukaryotic sequences in blue, and prokaryotic sequences in bright green.

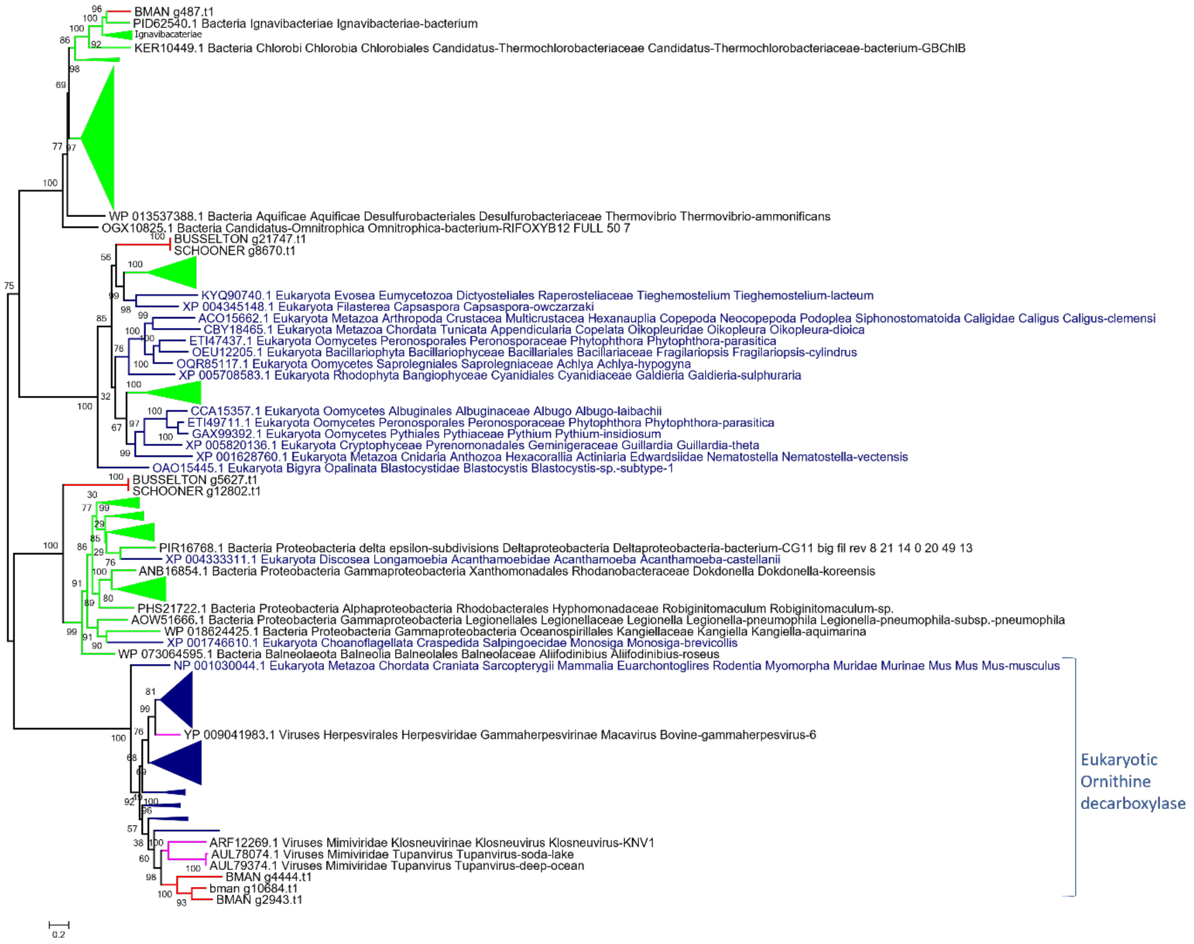


**Figure S13Q** Phylogenetic reconstruction of Diaminopimelate decarboxylase. Alignment with MAFFT, site selection BMGE and phylogeny IQtree model LG4X 1000 ultrafast bootstrap. *Anaeramoeba* sequences are in red, eukaryotic sequences in blue, viral sequences in fushia, and prokaryotic sequences in bright green.

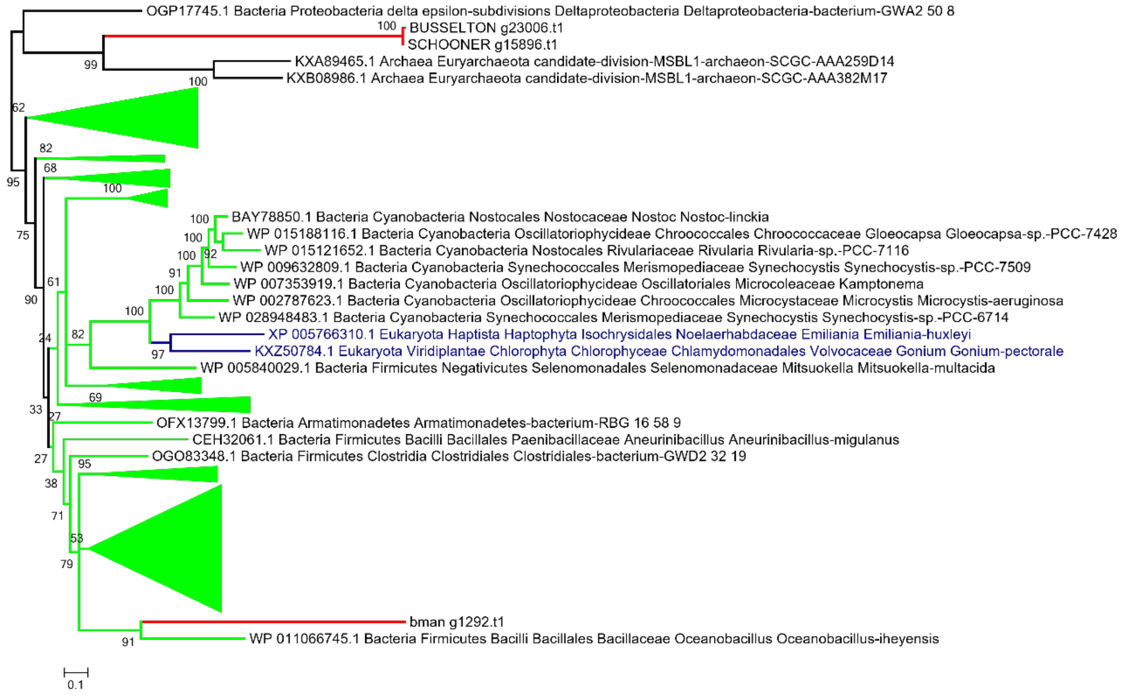


**Figure S13R** Phylogenetic reconstruction of Diaminopimelate epimerase. Alignment with MAFFT, site selection BMGE and phylogeny IQtree model LG4X 1000 ultrafast bootstrap. *Anaeramoeba* sequences are in red, eukaryotic sequences in blue, and prokaryotic sequences in bright green.

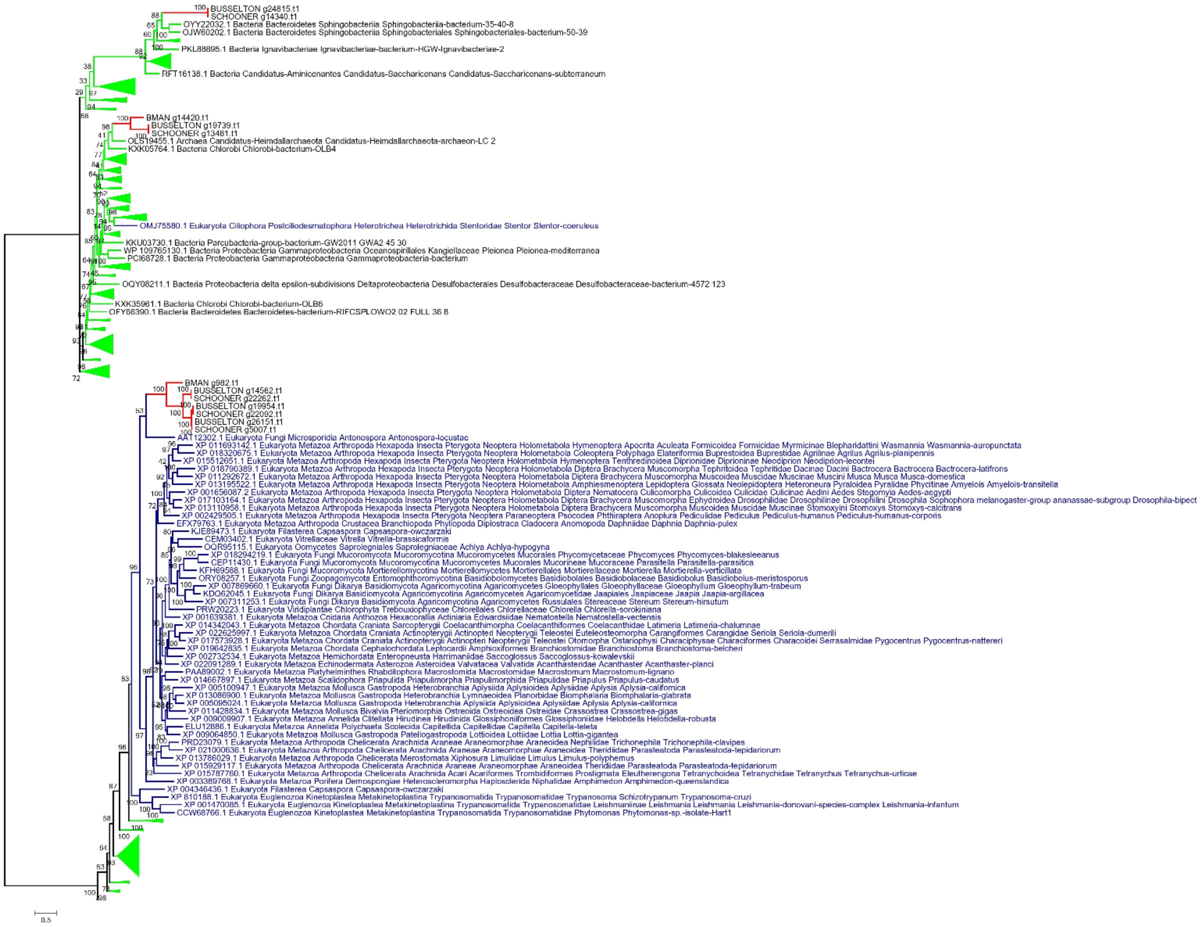

**Figure S13S** Phylogenetic reconstruction of Saccharopine dehydrogenase. Alignment with MAFFT, site selection BMGE and phylogeny IQtree model LG4X 1000 ultrafast bootstrap. *Anaeramoeba* sequences are in red, eukaryotic sequences in blue, and prokaryotic sequences in bright green.

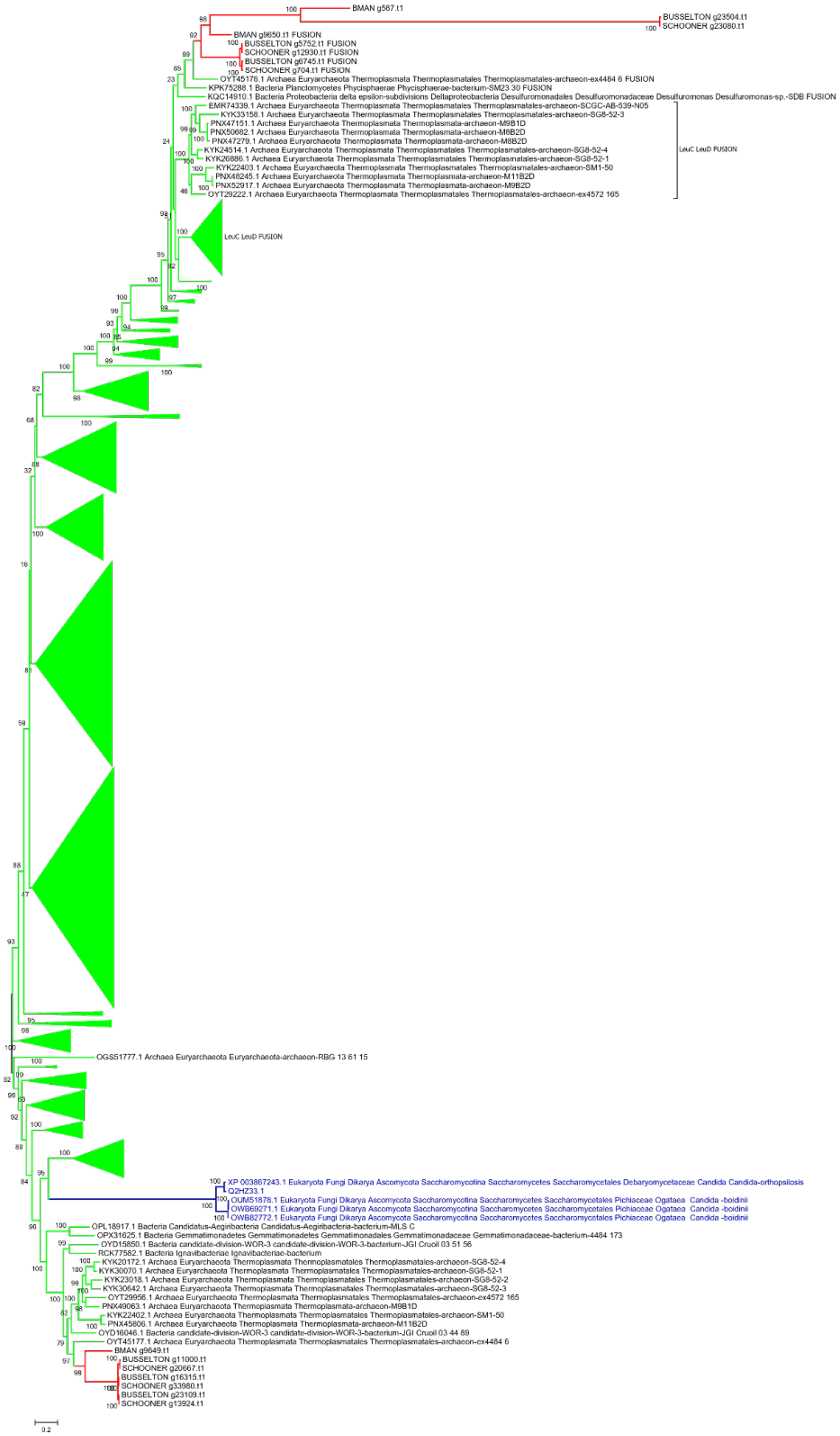

BMAN g587.t1
BUSSELTON g23504.t1
SCHOONER g23080.t1
BMAN g9650.t1 FUSION
BUSSELTON g5752.t1 FUSION
SCHOONER g12930.t1 FUSION
BUSSELTON g6745.t1 FUSION
SCHOONER g704.t1 FUSION
OYT45176.1 Archaea Euryarchaeota Thermoplasmata Thermoplasmatales Thermoplasmatales-archaeon-ex4484 6 FUSION
KPK75288.1 Bacteria Planctomycetes Phycisphaerae Phycisphaerae-bacterium-SM23 30 FUSION
KQC14910.1 Bacteria Proteobacteria delta epsilon-subdivisions Deltaproteobacteria Desulfuromonadales Desulfuromonadaceae Desulfuromonas Desulfuromonas-sp.-SDB FUSION
EMR74339.1 Archaea Euryarchaeota Thermoplasmata Thermoplasmatales Thermoplasmatales-archaeon-SCGC-AB-539-N05
KYK33158.1 Archaea Euryarchaeota Thermoplasmata Thermoplasmatales Thermoplasmatales-archaeon-SG8-52-3
PNX47151.1 Archaea Euryarchaeota Thermoplasmata Thermoplasmata-archaeon-M9B1D
PNX50682.1 Archaea Euryarchaeota Thermoplasmata Thermoplasmata-archaeon-M8B2D
PNX47279.1 Archaea Euryarchaeota Thermoplasmata Thermoplasmata-archaeon-M8B2D
KYK24514.1 Archaea Euryarchaeota Thermoplasmata Thermoplasmatales Thermoplasmatales-archaeon-SG8-52-4
KYK26886.1 Archaea Euryarchaeota Thermoplasmata Thermoplasmatales Thermoplasmatales-archaeon-SG8-52-1
KYK22403.1 Archaea Euryarchaeota Thermoplasmata Thermoplasmatales Thermoplasmatales-archaeon-SM1-50
PNX48245.1 Archaea Euryarchaeota Thermoplasmata Thermoplasmata-archaeon-M11B2D
PNX52917.1 Archaea Euryarchaeota Thermoplasmata Thermoplasmata-archaeon-M9B2D
OYT29222.1 Archaea Euryarchaeota Thermoplasmata Thermoplasmatales Thermoplasmatales-archaeon-ex4572 165
LeuC LeuD FUSION
LeuC LeuD FUSION
OGS51777.1 Archaea Euryarchaeota Euryarchaeota-archaeon-RBG 13 61 15
XP 003867243.1 Eukaryota Fungi Dikarya Ascomycota Saccharomycotina Saccharomycetes Saccharomycetales Debaryomycetaceae Candida Candida-orthopsilosis
Q2HZ33.1
OUM51878.1 Eukaryota Fungi Dikarya Ascomycota Saccharomycotina Saccharomycetes Saccharomycetales Pichiaceae Ogataea Candida -boidinii
OWB69271.1 Eukaryota Fungi Dikarya Ascomycota Saccharomycotina Saccharomycetes Saccharomycetales Pichiaceae Ogataea Candida -boidinii
OWB82772.1 Eukaryota Fungi Dikarya Ascomycota Saccharomycotina Saccharomycetes Saccharomycetales Pichiaceae Ogataea Candida -boidinii
OPL18917.1 Bacteria Candidatus-Aegiribacteria Candidatus-Aegiribacteria-bacterium-MLS C
OPX31625.1 Bacteria Gemmatimonadetes Gemmatimonadetes Gemmatimonadales Gemmatimonadaceae Gemmatimonadaceae-bacterium-4484 173
OYD15850.1 Bacteria candidate-division-WOR-3 candidate-division-WOR-3-bacterium-JGI Cruoil 03 51 56
RCK77582.1 Bacteria Ignavibacteriae Ignavibacteriae-bacterium
KYK20172.1 Archaea Euryarchaeota Thermoplasmata Thermoplasmatales Thermoplasmatales-archaeon-SG8-52-4
KYK30070.1 Archaea Euryarchaeota Thermoplasmata Thermoplasmatales Thermoplasmatales-archaeon-SG8-52-1
KYK23018.1 Archaea Euryarchaeota Thermoplasmata Thermoplasmatales Thermoplasmatales-archaeon-SG8-52-2
KYK30642.1 Archaea Euryarchaeota Thermoplasmata Thermoplasmatales Thermoplasmatales-archaeon-SG8-52-3
OYT29956.1 Archaea Euryarchaeota Thermoplasmata Thermoplasmatales Thermoplasmatales-archaeon-ex4572 165
PNX49063.1 Archaea Euryarchaeota Thermoplasmata Thermoplasmata-archaeon-M9B1D
KYK22402.1 Archaea Euryarchaeota Thermoplasmata Thermoplasmatales Thermoplasmatales-archaeon-SM1-50
PNX45806.1 Archaea Euryarchaeota Thermoplasmata Thermoplasmata-archaeon-M11B2D
OYD16046.1 Bacteria candidate-division-WOR-3 candidate-division-WOR-3-bacterium-JGI Cruoil 03 44 89
OYT45177.1 Archaea Euryarchaeota Thermoplasmata Thermoplasmatales Thermoplasmatales-archaeon-ex4484 6
BMAN g9649.t1
BUSSELTON g11000.t1
SCHOONER g20667.t1
BUSSELTON g16315.t1
SCHOONER g33980.t1
BUSSELTON g23109.t1
SCHOONER g13924.t1
0.2

**Figure S13T** Phylogenetic reconstruction of LeuC. Alignment with MAFFT, site selection BMGE and phylogeny IQtree model LG4X 1000 ultrafast bootstrap. *Anaeramoeba* sequences are in red, eukaryotic sequences in blue, and prokaryotic sequences in bright green.

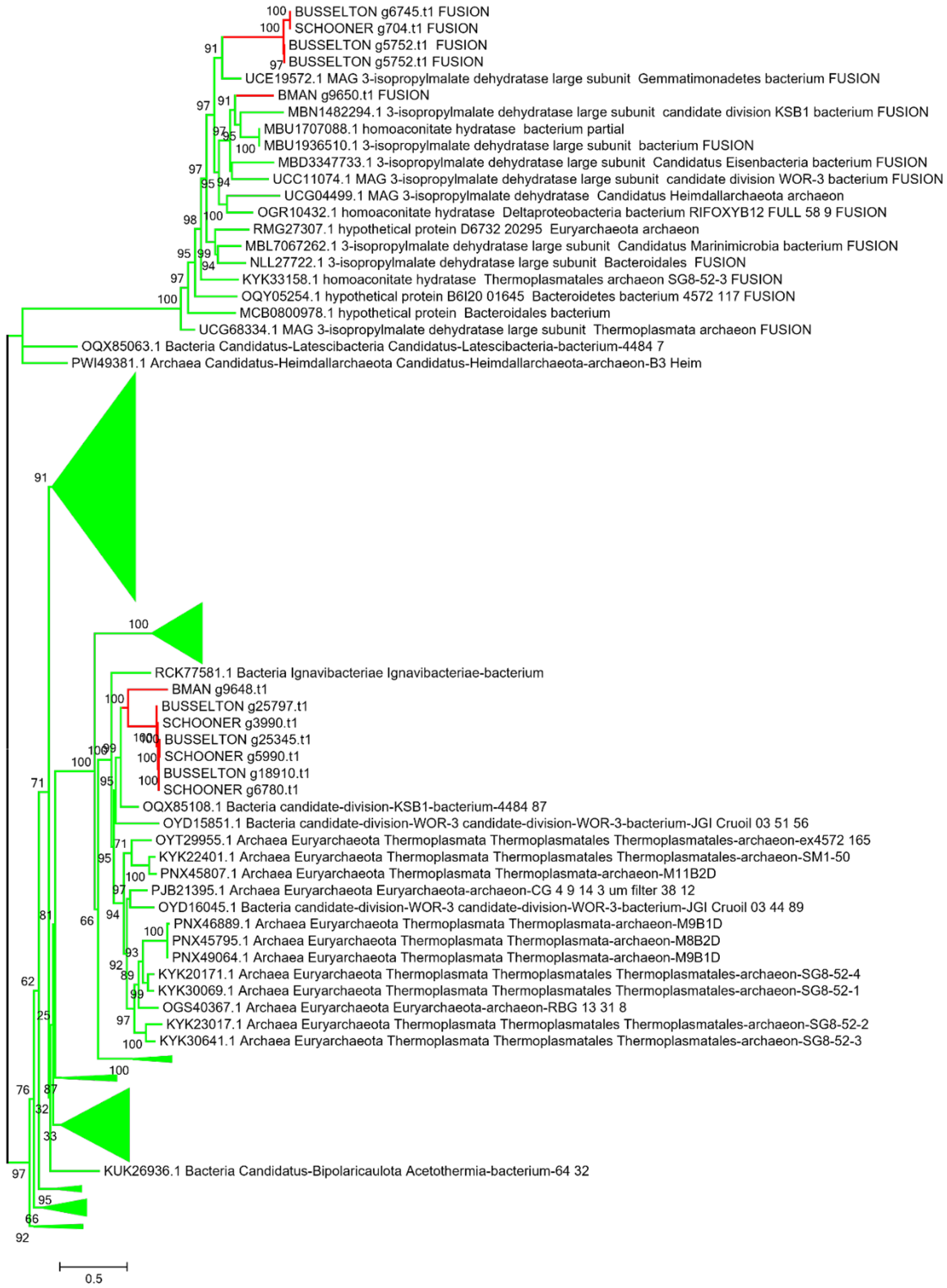


**Figure S13U** Phylogenetic reconstruction of LeuD. Alignment with MAFFT, site selection BMGE and phylogeny IQtree model LG4X 1000 ultrafast bootstrap. *Anaeramoeba* sequences are in red, eukaryotic sequences in blue, and prokaryotic sequences in bright green.

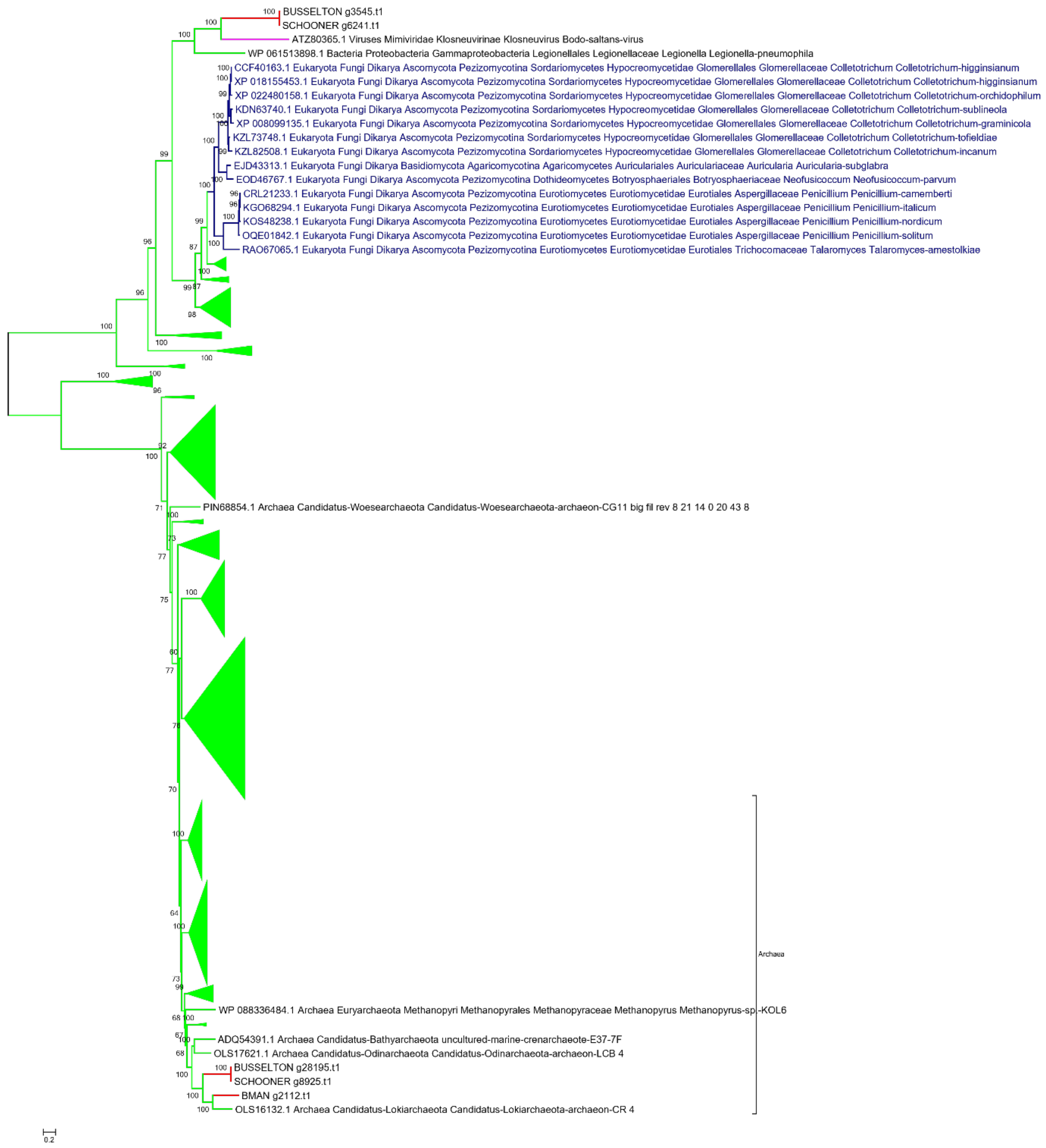


**Figure S13V** Phylogenetic reconstruction of FAD-dependent dehydrogenase. Alignment with MAFFT, site selection with BMGE and phylogeny with IQtree model C20+G4 with 1000 ultrafast bootstrap. Anaeramoeba sequences are in red, eukaryotic sequences in blue, viral sequences in light purple and prokaryotic sequences in bright green.

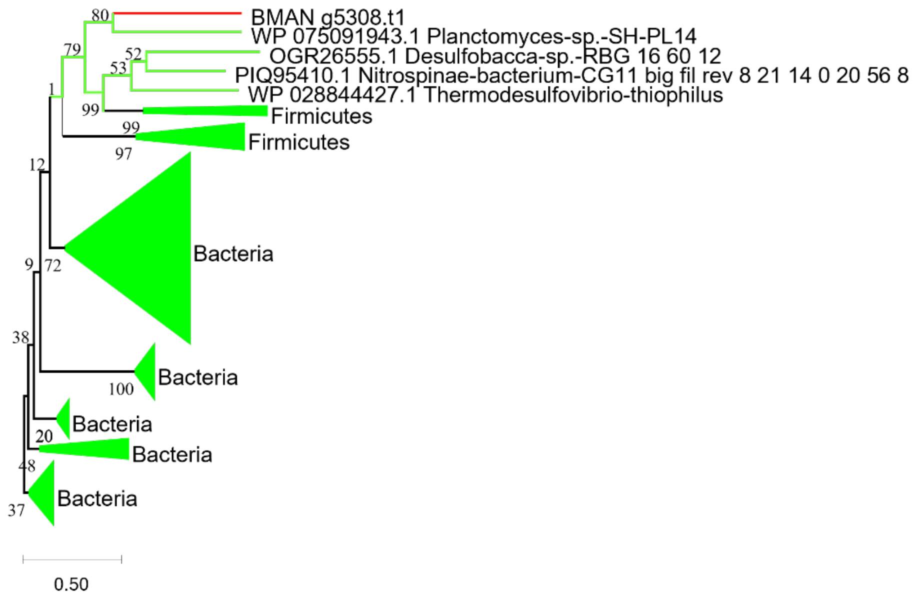


**Figure S13W** Phylogenetic reconstruction of CobC. Alignment with MAFFT, site selection BMGE and phylogeny IQtree model LG4X 1000 ultrafast bootstrap. *Anaeramoeba* sequences are in red, and prokaryotic sequences in bright green.

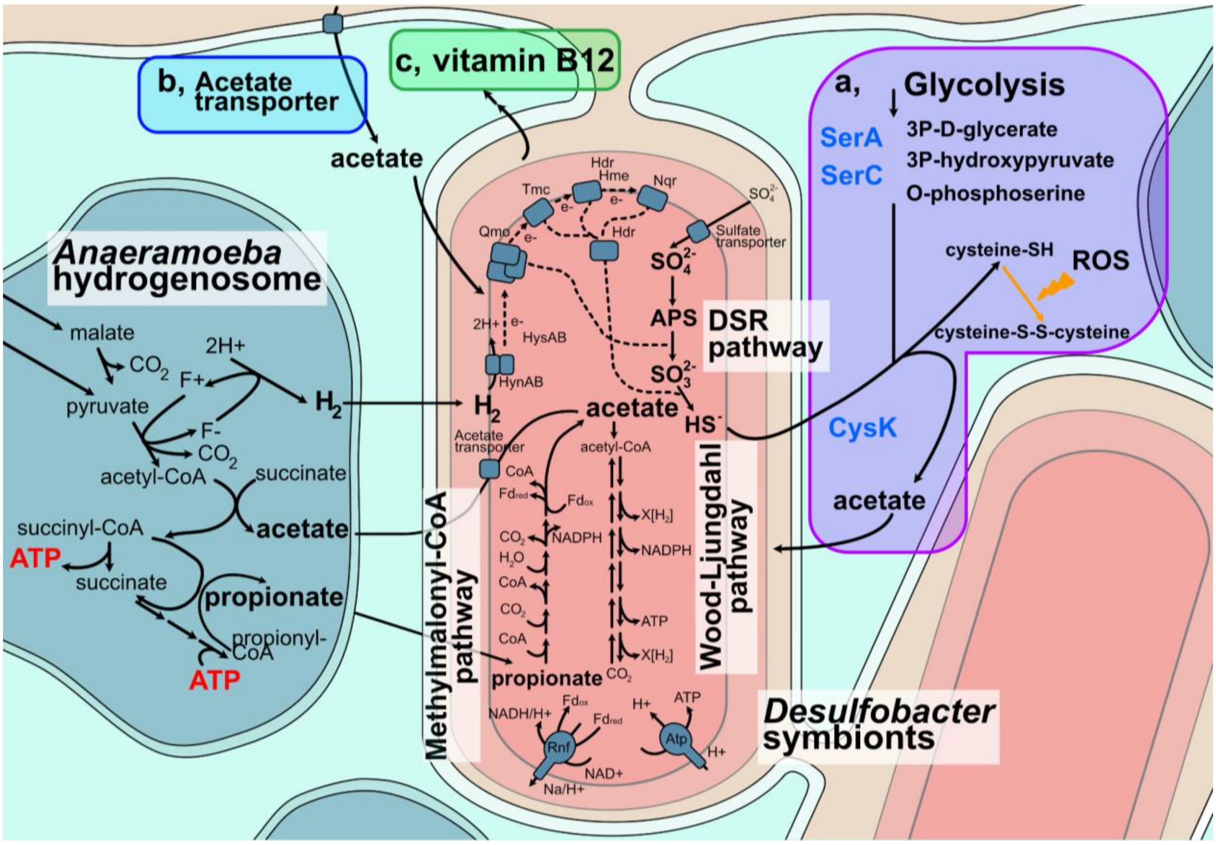


**Figure S14: *Anaeramoeba* uses ancestral and LGT-derived enzymes to interact with its symbionts.** Suggested syntrophic interactions between *Anaeramoeba* hydrogenosomes (blue) and symbionts (pink/salmon) based on metabolic reconstruction from transcriptomic and genomic evidence. The host's ATP-producing hydrogenosomes generate $H_2$, acetate, and propionate as end-products. Based on metatranscriptomic data, the symbionts use the products of the hydrogenosomes by prominently expressing the dissimilatory sulfate reduction (DSR), methylmalonyl-CoA, and the Wood-Ljungdahl pathways. The symbionts are in deep membrane-pits with a connection to the cell surrounding that gives ready access to sulfate (gold). **a,** A hybrid pathway to reuse $SH^-$ is formed by the ancestral SerC and the LGTs CysK and SerA (purple). The pathway would recycle acetate for the host or symbiont and provide reduced cysteine for mitigating ROS. **b,** The host has laterally transferred acetate transporters that might help to provide acetate to the symbionts (light blue). The actual localization and orientation of the acetate transporters are unknown, and we only show a suggested localization. **c,** Vitamin B12 is produced by the symbionts and might be transferred to the host that requires it as a cofactor for several enzymes (green). *A. ignava* BMAN has CobC, the final enzyme of the vitamin B12 synthesis pathway as an LGT.

A)

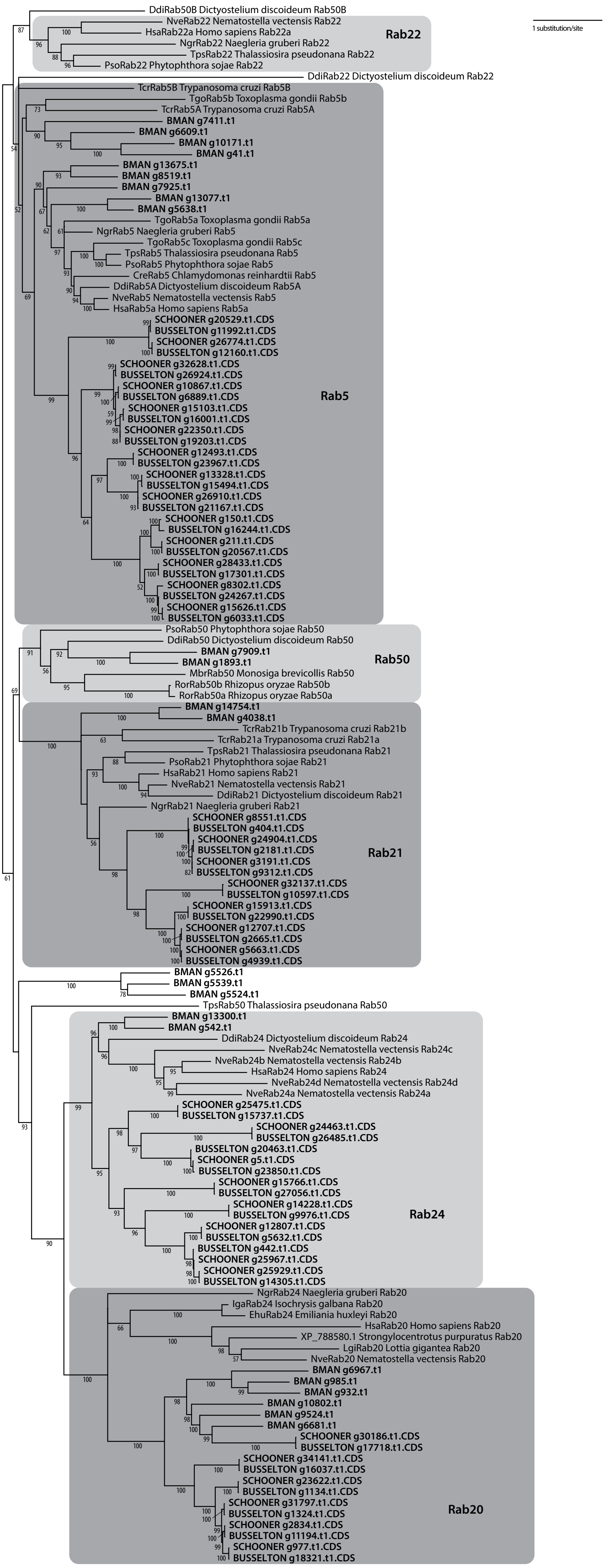
1 substitution/site
DdiRab50B Dictyostelium discoideum Rab50B
NveRab22 Nematostella vectensis Rab22
HsaRab22a Homo sapiens Rab22a
NgrRab22 Naegleria gruberi Rab22
TpsRab22 Thalassiosira pseudonana Rab22
PsoRab22 Phytophthora sojae Rab22
Rab22
DdiRab22 Dictyostelium discoideum Rab22
TcrRab5B Trypanosoma cruzi Rab5B
TgoRab5b Toxoplasma gondii Rab5b
TcrRab5A Trypanosoma cruzi Rab5A
BMAN g7411.t1
BMAN g6609.t1
BMAN g10171.t1
BMAN g41.t1
BMAN g13675.t1
BMAN g8519.t1
BMAN g7925.t1
BMAN g13077.t1
BMAN g5638.t1
TgoRab5a Toxoplasma gondii Rab5a
NgrRab5 Naegleria gruberi Rab5
TgoRab5c Toxoplasma gondii Rab5c
TpsRab5 Thalassiosira pseudonana Rab5
PsoRab5 Phytophthora sojae Rab5
CreRab5 Chlamydomonas reinhardtii Rab5
DdiRab5A Dictyostelium discoideum Rab5A
NveRab5 Nematostella vectensis Rab5
HsaRab5a Homo sapiens Rab5a
SCHOONER g20529.t1.CDS
BUSSELTON g11992.t1.CDS
SCHOONER g26774.t1.CDS
BUSSELTON g12160.t1.CDS
SCHOONER g32628.t1.CDS
BUSSELTON g26924.t1.CDS
SCHOONER g10867.t1.CDS
BUSSELTON g6889.t1.CDS
SCHOONER g15103.t1.CDS
BUSSELTON g16001.t1.CDS
SCHOONER g22350.t1.CDS
BUSSELTON g19203.t1.CDS
SCHOONER g12493.t1.CDS
BUSSELTON g23967.t1.CDS
SCHOONER g13328.t1.CDS
BUSSELTON g15494.t1.CDS
SCHOONER g26910.t1.CDS
BUSSELTON g21167.t1.CDS
SCHOONER g150.t1.CDS
BUSSELTON g16244.t1.CDS
SCHOONER g211.t1.CDS
BUSSELTON g20567.t1.CDS
SCHOONER g28433.t1.CDS
BUSSELTON g17301.t1.CDS
SCHOONER g8302.t1.CDS
BUSSELTON g24267.t1.CDS
SCHOONER g15626.t1.CDS
BUSSELTON g6033.t1.CDS
Rab5
PsoRab50 Phytophthora sojae Rab50
DdiRab50 Dictyostelium discoideum Rab50
BMAN g7909.t1
BMAN g1893.t1
MbrRab50 Monosiga brevicollis Rab50
RorRab50b Rhizopus oryzae Rab50b
RorRab50a Rhizopus oryzae Rab50a
Rab50
BMAN g14754.t1
BMAN g4038.t1
TcrRab21b Trypanosoma cruzi Rab21b
TcrRab21a Trypanosoma cruzi Rab21a
TpsRab21 Thalassiosira pseudonana Rab21
PsoRab21 Phytophthora sojae Rab21
HsaRab21 Homo sapiens Rab21
NveRab21 Nematostella vectensis Rab21
DdiRab21 Dictyostelium discoideum Rab21
NgrRab21 Naegleria gruberi Rab21
SCHOONER g8551.t1.CDS
BUSSELTON g404.t1.CDS
SCHOONER g24904.t1.CDS
BUSSELTON g2181.t1.CDS
SCHOONER g3191.t1.CDS
BUSSELTON g9312.t1.CDS
SCHOONER g32137.t1.CDS
BUSSELTON g10597.t1.CDS
SCHOONER g15913.t1.CDS
BUSSELTON g22990.t1.CDS
SCHOONER g12707.t1.CDS
BUSSELTON g2665.t1.CDS
SCHOONER g5663.t1.CDS
BUSSELTON g4939.t1.CDS
Rab21
BMAN g5526.t1
BMAN g5539.t1
BMAN g5524.t1
TpsRab50 Thalassiosira pseudonana Rab50
BMAN g13300.t1
BMAN g542.t1
DdiRab24 Dictyostelium discoideum Rab24
NveRab24c Nematostella vectensis Rab24c
NveRab24b Nematostella vectensis Rab24b
HsaRab24 Homo sapiens Rab24
NveRab24d Nematostella vectensis Rab24d
NveRab24a Nematostella vectensis Rab24a
SCHOONER g25475.t1.CDS
BUSSELTON g15737.t1.CDS
SCHOONER g24463.t1.CDS
BUSSELTON g26485.t1.CDS
BUSSELTON g20463.t1.CDS
SCHOONER g5.t1.CDS
BUSSELTON g23850.t1.CDS
SCHOONER g15766.t1.CDS
BUSSELTON g27056.t1.CDS
SCHOONER g14228.t1.CDS
BUSSELTON g9976.t1.CDS
SCHOONER g12807.t1.CDS
BUSSELTON g5632.t1.CDS
BUSSELTON g442.t1.CDS
SCHOONER g25967.t1.CDS
SCHOONER g25929.t1.CDS
BUSSELTON g14305.t1.CDS
Rab24
NgrRab24 Naegleria gruberi Rab20
IgaRab24 Isochrysis galbana Rab20
EhuRab24 Emiliania huxleyi Rab20
HsaRab20 Homo sapiens Rab20
XP_788580.1 Strongylocentrotus purpuratus Rab20
LgiRab20 Lottia gigantea Rab20
NveRab20 Nematostella vectensis Rab20
BMAN g6967.t1
BMAN g985.t1
BMAN g932.t1
BMAN g10802.t1
BMAN g9524.t1
BMAN g6681.t1
SCHOONER g30186.t1.CDS
BUSSELTON g17718.t1.CDS
SCHOONER g34141.t1.CDS
BUSSELTON g16037.t1.CDS
SCHOONER g23622.t1.CDS
BUSSELTON g1134.t1.CDS
SCHOONER g31797.t1.CDS
BUSSELTON g1324.t1.CDS
SCHOONER g2834.t1.CDS
BUSSELTON g11194.t1.CDS
SCHOONER g977.t1.CDS
BUSSELTON g18321.t1.CDS
Rab20

B)

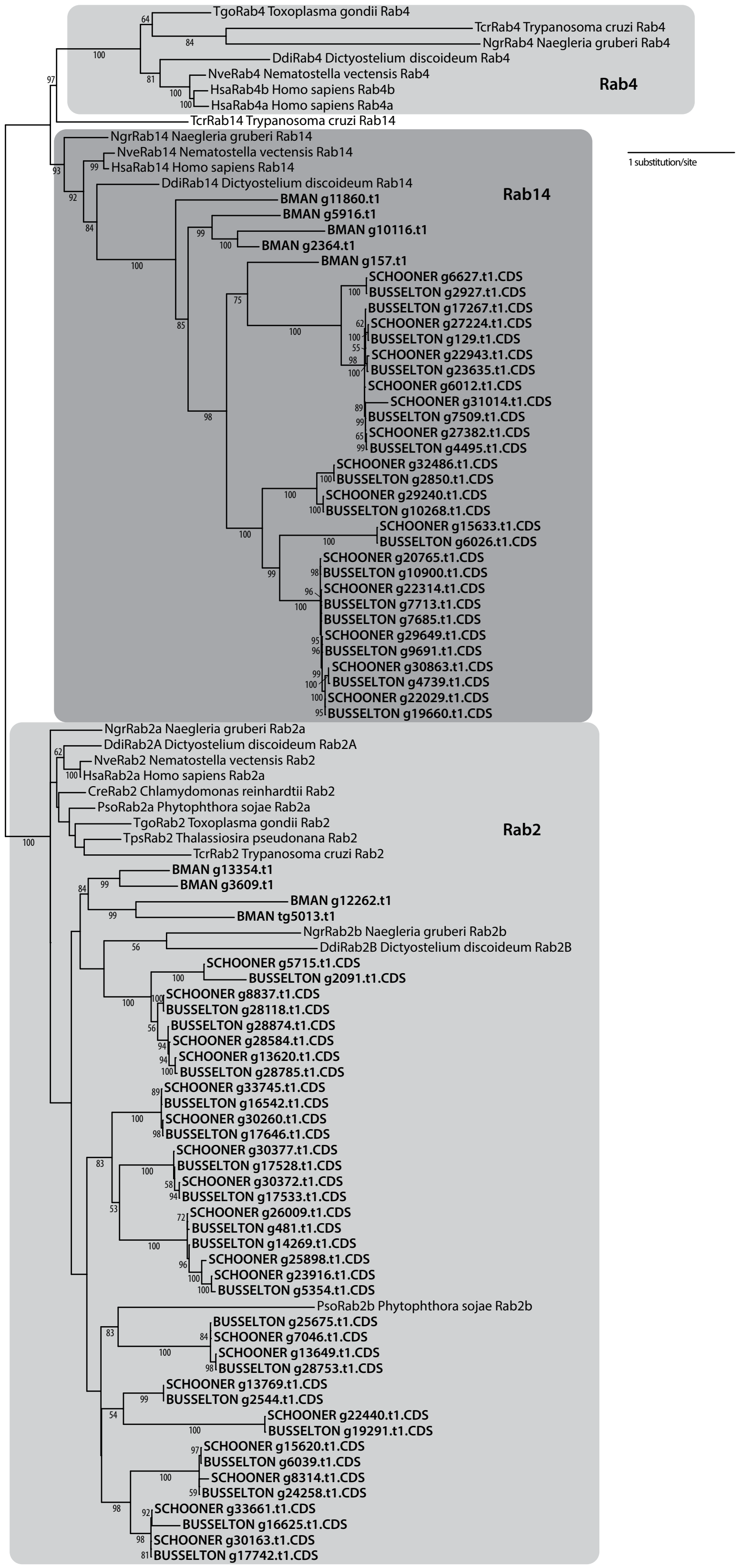
TgoRab4 Toxoplasma gondii Rab4
TcrRab4 Trypanosoma cruzi Rab4
NgrRab4 Naegleria gruberi Rab4
DdiRab4 Dictyostelium discoideum Rab4
NveRab4 Nematostella vectensis Rab4
HsaRab4b Homo sapiens Rab4b
HsaRab4a Homo sapiens Rab4a
Rab4
TcrRab14 Trypanosoma cruzi Rab14
1 substitution/site
NgrRab14 Naegleria gruberi Rab14
NveRab14 Nematostella vectensis Rab14
HsaRab14 Homo sapiens Rab14
DdiRab14 Dictyostelium discoideum Rab14
Rab14
BMAN g11860.t1
BMAN g5916.t1
BMAN g10116.t1
BMAN g2364.t1
BMAN g157.t1
SCHOONER g6627.t1.CDS
BUSSELTON g2927.t1.CDS
BUSSELTON g17267.t1.CDS
SCHOONER g27224.t1.CDS
BUSSELTON g129.t1.CDS
SCHOONER g22943.t1.CDS
BUSSELTON g23635.t1.CDS
SCHOONER g6012.t1.CDS
SCHOONER g31014.t1.CDS
BUSSELTON g7509.t1.CDS
SCHOONER g27382.t1.CDS
BUSSELTON g4495.t1.CDS
SCHOONER g32486.t1.CDS
BUSSELTON g2850.t1.CDS
SCHOONER g29240.t1.CDS
BUSSELTON g10268.t1.CDS
SCHOONER g15633.t1.CDS
BUSSELTON g6026.t1.CDS
SCHOONER g20765.t1.CDS
BUSSELTON g10900.t1.CDS
SCHOONER g22314.t1.CDS
BUSSELTON g7713.t1.CDS
BUSSELTON g7685.t1.CDS
SCHOONER g29649.t1.CDS
BUSSELTON g9691.t1.CDS
SCHOONER g30863.t1.CDS
BUSSELTON g4739.t1.CDS
SCHOONER g22029.t1.CDS
BUSSELTON g19660.t1.CDS
NgrRab2a Naegleria gruberi Rab2a
DdiRab2A Dictyostelium discoideum Rab2A
NveRab2 Nematostella vectensis Rab2
HsaRab2a Homo sapiens Rab2a
CreRab2 Chlamydomonas reinhardtii Rab2
PsoRab2a Phytophthora sojae Rab2a
TgoRab2 Toxoplasma gondii Rab2
TpsRab2 Thalassiosira pseudonana Rab2
TcrRab2 Trypanosoma cruzi Rab2
Rab2
BMAN g13354.t1
BMAN g3609.t1
BMAN g12262.t1
BMAN tg5013.t1
NgrRab2b Naegleria gruberi Rab2b
DdiRab2B Dictyostelium discoideum Rab2B
SCHOONER g5715.t1.CDS
BUSSELTON g2091.t1.CDS
SCHOONER g8837.t1.CDS
BUSSELTON g28118.t1.CDS
BUSSELTON g28874.t1.CDS
SCHOONER g28584.t1.CDS
SCHOONER g13620.t1.CDS
BUSSELTON g28785.t1.CDS
SCHOONER g33745.t1.CDS
BUSSELTON g16542.t1.CDS
SCHOONER g30260.t1.CDS
BUSSELTON g17646.t1.CDS
SCHOONER g30377.t1.CDS
BUSSELTON g17528.t1.CDS
SCHOONER g30372.t1.CDS
BUSSELTON g17533.t1.CDS
SCHOONER g26009.t1.CDS
BUSSELTON g481.t1.CDS
BUSSELTON g14269.t1.CDS
SCHOONER g25898.t1.CDS
SCHOONER g23916.t1.CDS
BUSSELTON g5354.t1.CDS
PsoRab2b Phytophthora sojae Rab2b
BUSSELTON g25675.t1.CDS
SCHOONER g7046.t1.CDS
SCHOONER g13649.t1.CDS
BUSSELTON g28753.t1.CDS
SCHOONER g13769.t1.CDS
BUSSELTON g2544.t1.CDS
SCHOONER g22440.t1.CDS
BUSSELTON g19291.t1.CDS
SCHOONER g15620.t1.CDS
BUSSELTON g6039.t1.CDS
SCHOONER g8314.t1.CDS
BUSSELTON g24258.t1.CDS
SCHOONER g33661.t1.CDS
BUSSELTON g16625.t1.CDS
SCHOONER g30163.t1.CDS
BUSSELTON g17742.t1.CDS

C)

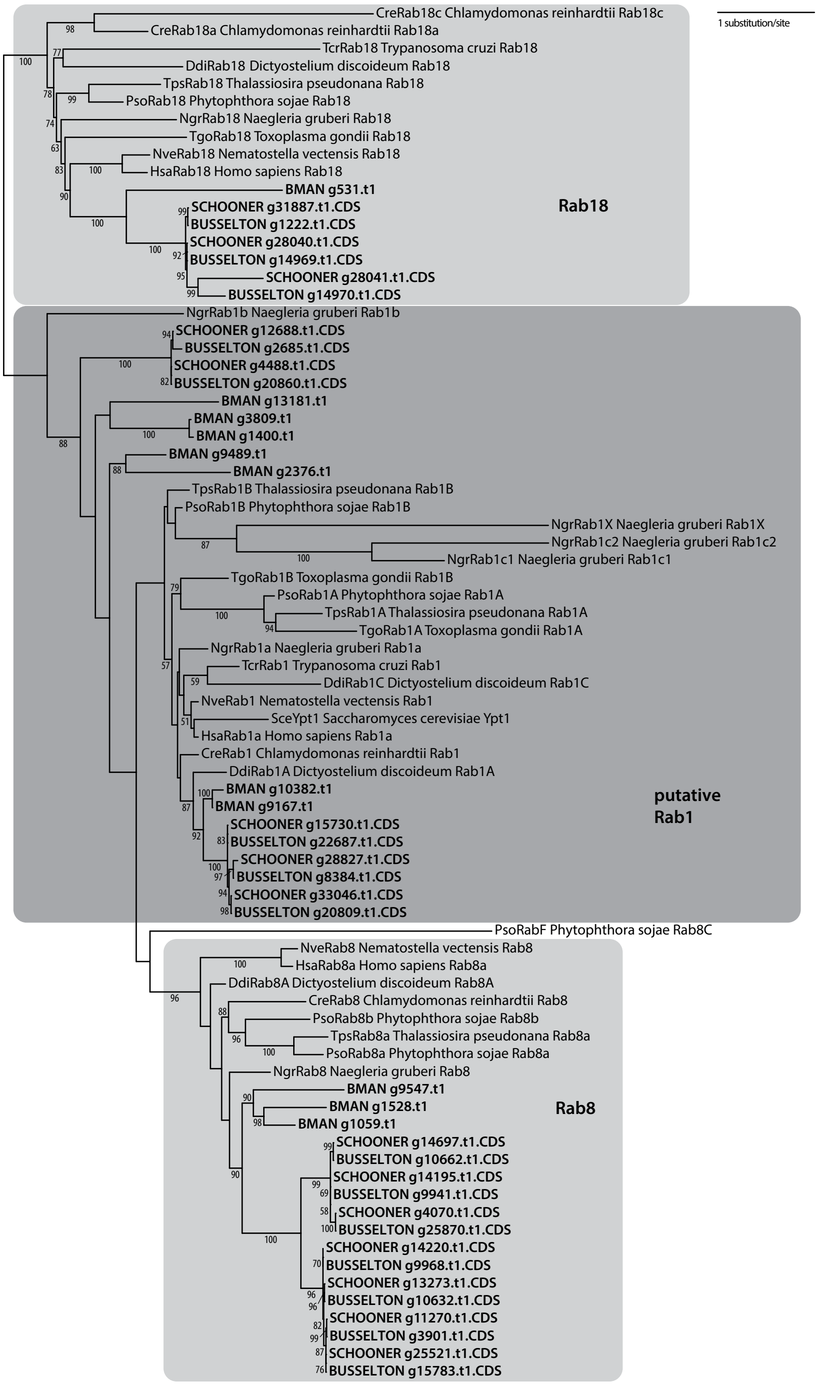

1 substitution/site
CreRab18c Chlamydomonas reinhardtii Rab18c
CreRab18a Chlamydomonas reinhardtii Rab18a
TcrRab18 Trypanosoma cruzi Rab18
DdiRab18 Dictyostelium discoideum Rab18
TpsRab18 Thalassiosira pseudonana Rab18
PsoRab18 Phytophthora sojae Rab18
NgrRab18 Naegleria gruberi Rab18
TgoRab18 Toxoplasma gondii Rab18
NveRab18 Nematostella vectensis Rab18
HsaRab18 Homo sapiens Rab18
BMAN g531.t1
SCHOONER g31887.t1.CDS
BUSSELTON g1222.t1.CDS
SCHOONER g28040.t1.CDS
BUSSELTON g14969.t1.CDS
SCHOONER g28041.t1.CDS
BUSSELTON g14970.t1.CDS
Rab18
NgrRab1b Naegleria gruberi Rab1b
SCHOONER g12688.t1.CDS
BUSSELTON g2685.t1.CDS
SCHOONER g4488.t1.CDS
BUSSELTON g20860.t1.CDS
BMAN g13181.t1
BMAN g3809.t1
BMAN g1400.t1
BMAN g9489.t1
BMAN g2376.t1
TpsRab1B Thalassiosira pseudonana Rab1B
PsoRab1B Phytophthora sojae Rab1B
NgrRab1X Naegleria gruberi Rab1X
NgrRab1c2 Naegleria gruberi Rab1c2
NgrRab1c1 Naegleria gruberi Rab1c1
TgoRab1B Toxoplasma gondii Rab1B
PsoRab1A Phytophthora sojae Rab1A
TpsRab1A Thalassiosira pseudonana Rab1A
TgoRab1A Toxoplasma gondii Rab1A
NgrRab1a Naegleria gruberi Rab1a
TcrRab1 Trypanosoma cruzi Rab1
DdiRab1C Dictyostelium discoideum Rab1C
NveRab1 Nematostella vectensis Rab1
SceYpt1 Saccharomyces cerevisiae Ypt1
HsaRab1a Homo sapiens Rab1a
CreRab1 Chlamydomonas reinhardtii Rab1
DdiRab1A Dictyostelium discoideum Rab1A
BMAN g10382.t1
BMAN g9167.t1
SCHOONER g15730.t1.CDS
BUSSELTON g22687.t1.CDS
SCHOONER g28827.t1.CDS
BUSSELTON g8384.t1.CDS
SCHOONER g33046.t1.CDS
BUSSELTON g20809.t1.CDS
putative Rab1
PsoRabF Phytophthora sojae Rab8C
NveRab8 Nematostella vectensis Rab8
HsaRab8a Homo sapiens Rab8a
DdiRab8A Dictyostelium discoideum Rab8A
CreRab8 Chlamydomonas reinhardtii Rab8
PsoRab8b Phytophthora sojae Rab8b
TpsRab8a Thalassiosira pseudonana Rab8a
PsoRab8a Phytophthora sojae Rab8a
NgrRab8 Naegleria gruberi Rab8
BMAN g9547.t1
BMAN g1528.t1
BMAN g1059.t1
SCHOONER g14697.t1.CDS
BUSSELTON g10662.t1.CDS
SCHOONER g14195.t1.CDS
BUSSELTON g9941.t1.CDS
SCHOONER g4070.t1.CDS
BUSSELTON g25870.t1.CDS
SCHOONER g14220.t1.CDS
BUSSELTON g9968.t1.CDS
SCHOONER g13273.t1.CDS
BUSSELTON g10632.t1.CDS
SCHOONER g11270.t1.CDS
BUSSELTON g3901.t1.CDS
SCHOONER g25521.t1.CDS
BUSSELTON g15783.t1.CDS
Rab8

D)

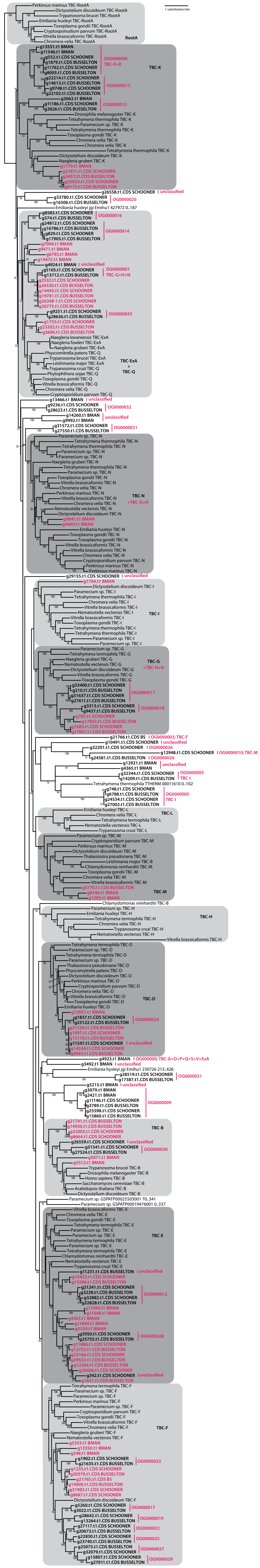
1 substitution/site
RootA
TBC-K
TBC-ExA
+
TBC-Q
TBC-N
+TBC-G+H
TBC-I
TBC-G
+TBC-H+N
TBC-L
TBC-M
TBC-H
TBC-D
TBC-B
TBC-E
TBC-F

**Figure S15: Phylogenetic analyses of Rab and TBC proteins in *Anaeramoeba*. a-c,** Phylogenetic analyses of Rab proteins of subfamilies containing endocytic Rabs **a,** Rabs 2, 4, and 14 **b,** and Rabs 1, 8, and 18 **c,**. **d,** Phylogenetic analysis of TBC proteins. *Anaeramoeba* sequences are in bold. Sequences having the same identity in the phylogenetic analysis and orthogroup (OG) clustering (Table S10D) are in pink. Sequences in black have OG identity displayed next to them. Tree files and alignments are available at FigShare: https://doi.org/10.6084/m9.figshare.22193497.v1

a.

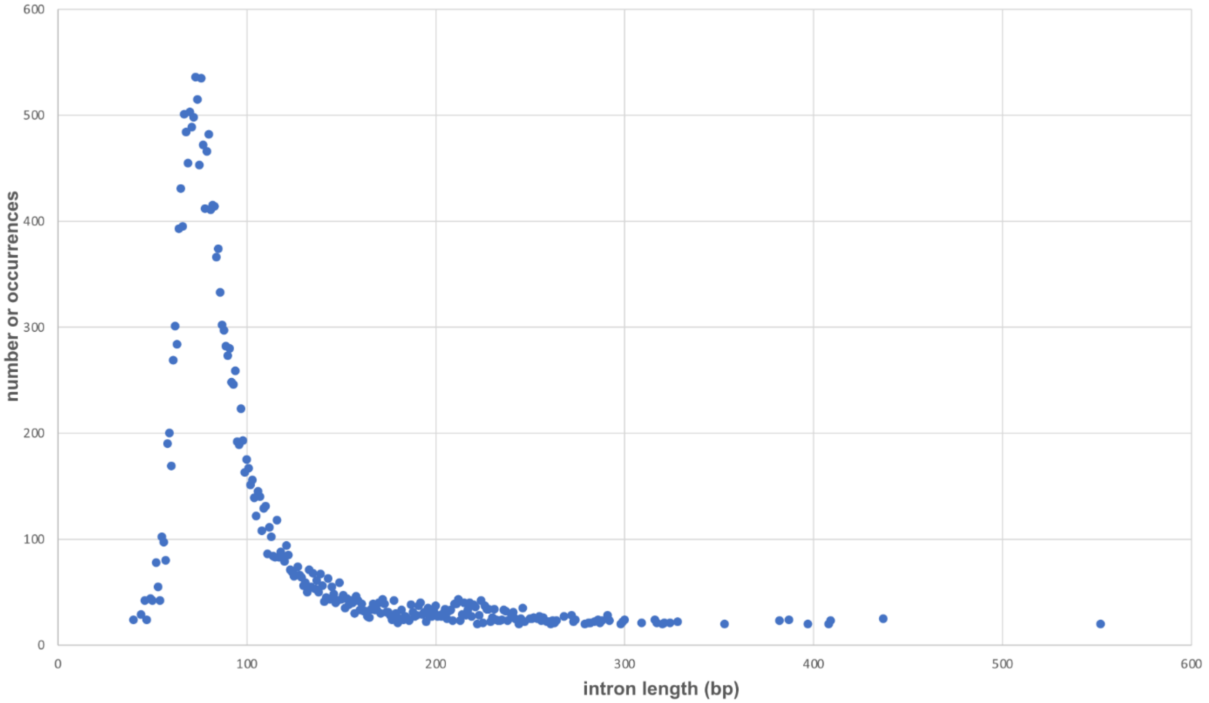


**Figure S16A** Intron lengths in *A. flamelloides* BMAN.

b.

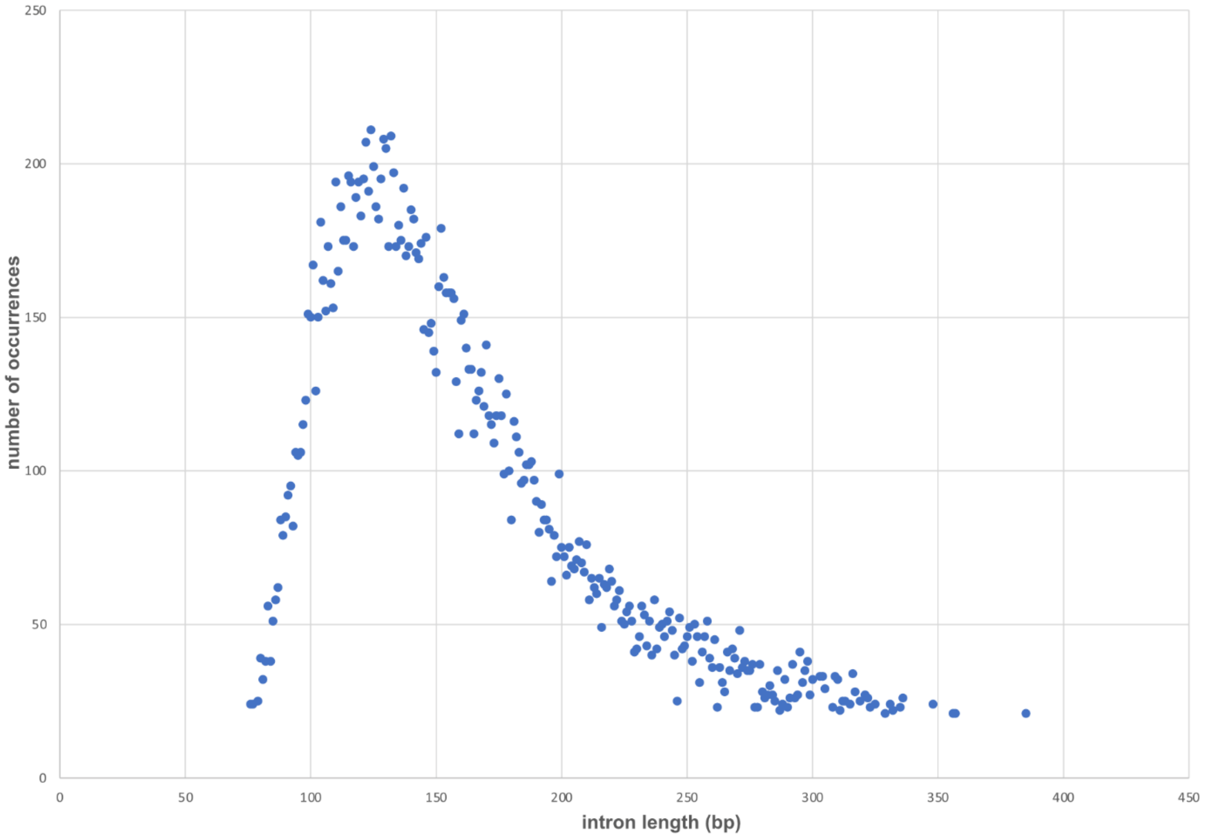


**Figure S16B** Intron lengths in *A. flamelloides* BUSSELTON2.

c.

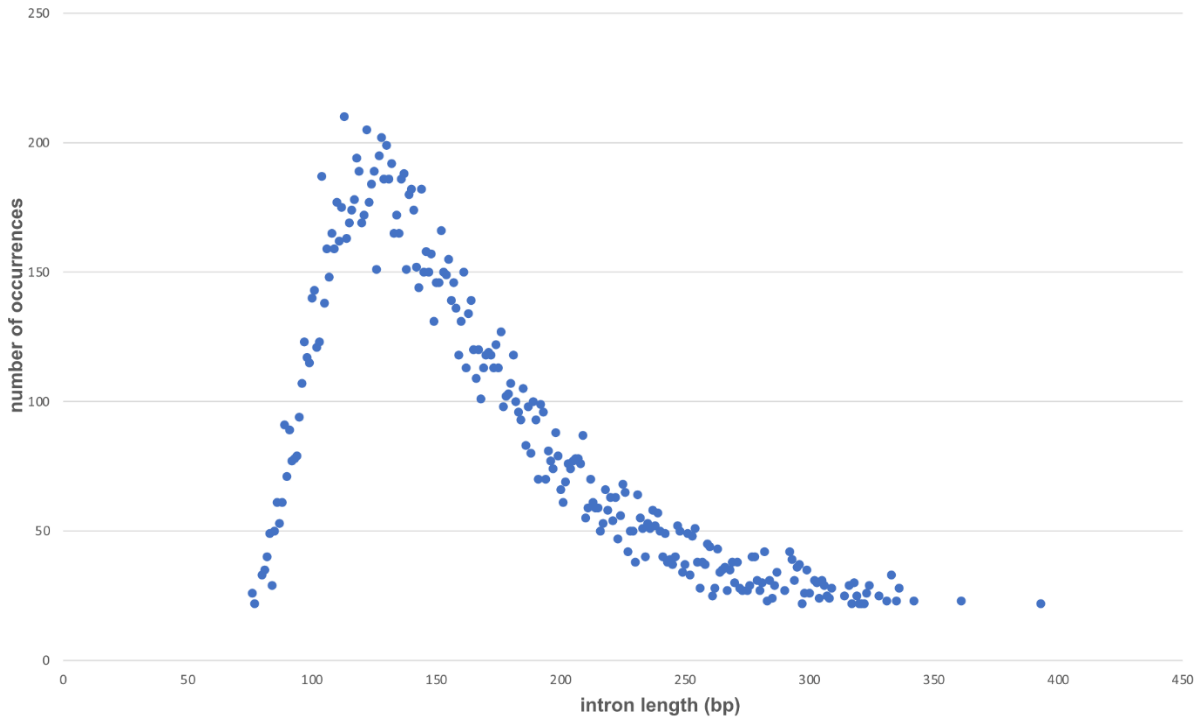


**Figure S16C** Intron lengths in *A. flamelloides* SCHOONER1.

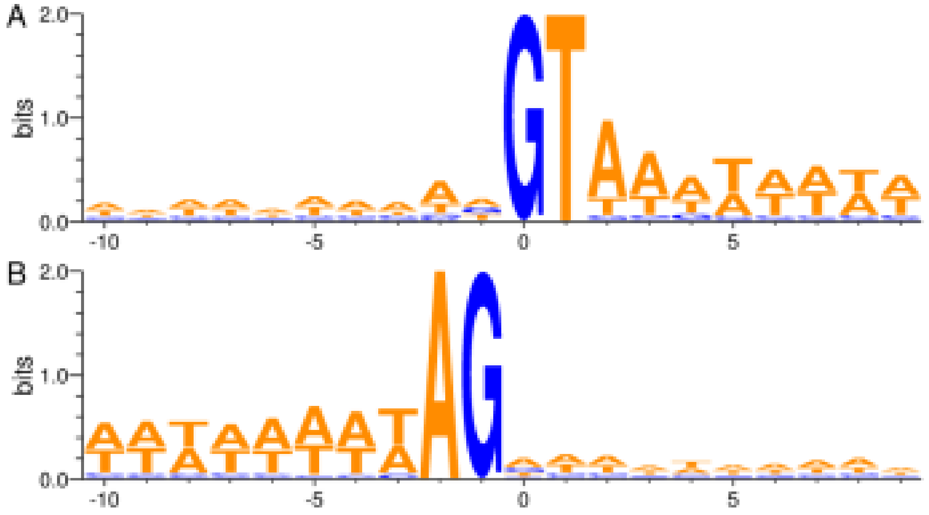

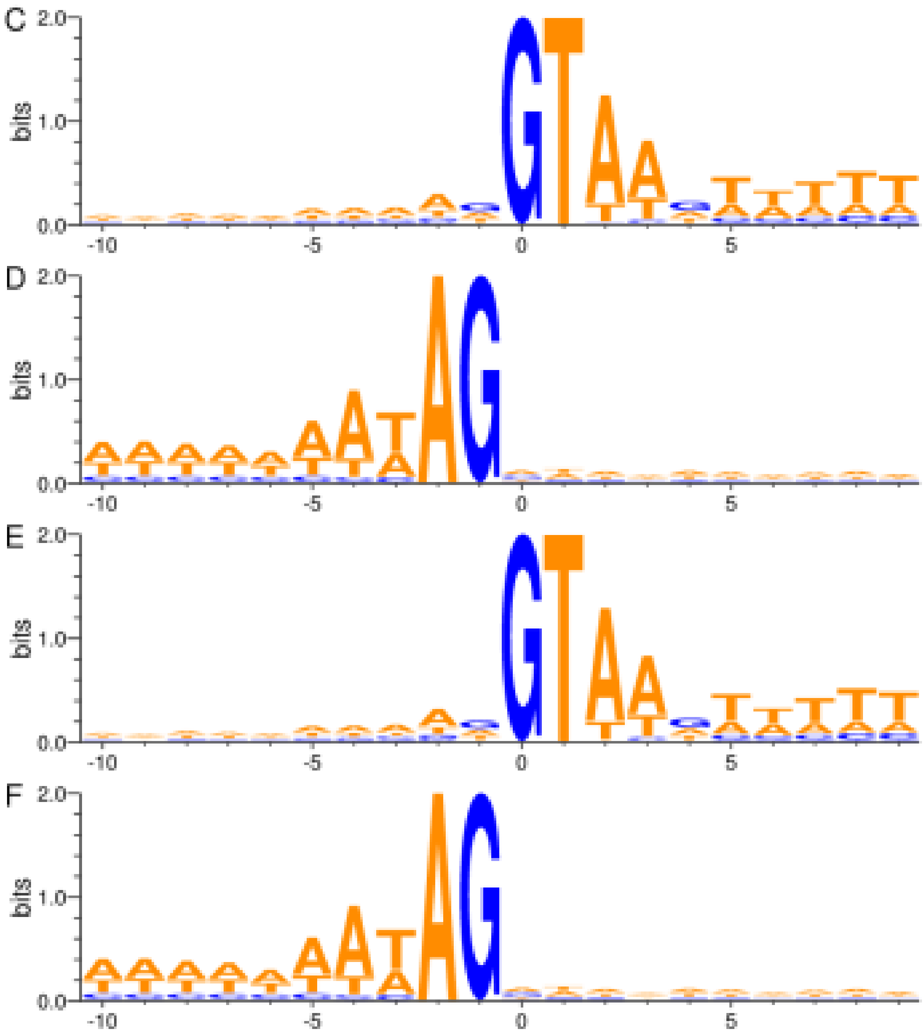


**Figure 16D Characteristics of intron splice sites.** Sequence logo plots showing subpanel **da.**, 10 bases upstream of donor sites and 8 downstream in BMAN, subpanel **db.**, 8 bases upstream of acceptor site and 10 bases downstream in BMAN, subpanel **dc.**, 10 bases upstream of donor sites and 8 downstream in BUSSELTON2, subpanel **dd.**, 8 bases upstream of acceptor site and 10 bases downstream in BUSSELTON2, subpanel **de.**, 10 bases upstream of donor sites and 8 downstream in SCHOONER1, subpanel **df.**, 8 bases upstream of acceptor site and 10 bases downstream in SCHOONER1. The sequence logos were generated using the WebLogo3 (Crooks et al. 2004) server.

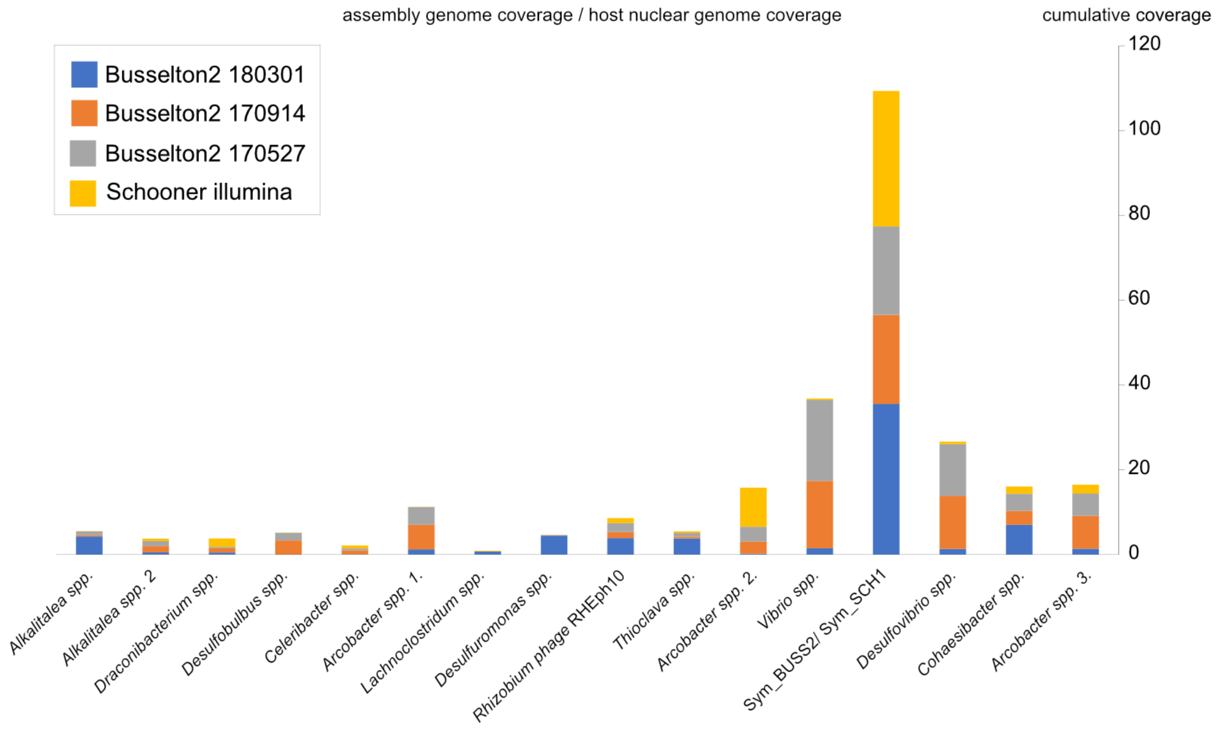


**Figure S17: Sym_BUSS2 / Sym_SCH1 are consistently associated with *Anaeramoeba flamelloides*.** Four sequencing datasets generated from DNA from four individual *A. flamelloides* cell isolations were mapped using minimap2 against either *A. flamelloides* BUSSELTON2 or SCHOONER1 genome and the total metagenome assemblies. The assembly genome / host nuclear genome coverages were calculated from bam-files using coverage statistics from pileup.sh in the BBMap package v38.20 (sourceforge.net/projects/bbmap/). The cumulative coverage was plotted with coverage from the different replicates indicated by blue (*A. flamelloides* BUSSELTON2 180301), orange (*A. flamelloides* BUSSELTON2 170914), grey (*A. flamelloides* BUSSELTON2 170527) and yellow (Schooner illumina). The different labels on the x-axis are major assembly components derived from metagenomic assemblies of *A. flamelloides* BUSSELTON2 and SCHOONER1. The assembly component annotations were assigned using the top blastn hit in NCBI.
